# Chemotherapy-induced intestinal injury promotes Galectin-9-driven modulation of T cell function

**DOI:** 10.1101/2023.04.30.538862

**Authors:** Suze A. Jansen, Alessandro Cutilli, Coco de Koning, Marliek van Hoesel, Leire Saiz Sierra, Stefan Nierkens, Michal Mokry, Edward E.S. Nieuwenhuis, Alan M. Hanash, Enric Mocholi, Paul J. Coffer, Caroline A. Lindemans

## Abstract

The intestine is vulnerable to chemotherapy-induced toxicity due to its high epithelial proliferative rate, making gut toxicity an off-target effect in several cancer treatments, including conditioning regimens for allogeneic hematopoietic cell transplantation (allo-HCT). In allo-HCT, intestinal damage is an important factor in the development of Graft-versus-Host Disease (GVHD), an immune complication in which donor immune cells attack the recipient’s tissues. Here, we developed a novel human intestinal organoid-based 3D model system to study the direct effect of chemotherapy-induced intestinal epithelial damage on T cell behavior. Chemotherapy treatment using busulfan, fludarabine, and clofarabine led to damage responses in organoids resulting in increased T cell migration, activation, and proliferation in ex-*vivo* co-culture assays. We identified galectin-9 (Gal-9), a beta-galactoside-binding lectin released by damaged organoids, as a key molecule mediating T cell responses to damage. Increased levels of Gal-9 were also found in the plasma of allo-HCT patients who later developed acute GVHD, supporting the predictive value of the model system in the clinical setting. This study highlights the potential contribution of chemotherapy-induced epithelial damage to the pathogenesis of intestinal GVHD through direct effects on T cell activation and trafficking promoted by galectin-9.

## Introduction

The intestinal epithelium is comprised of a variety of secretory and absorptive intestinal epithelial cell (IEC) types that derive from the intestinal stem cells (ISC) at the bottom of intestinal epithelial crypts. As the IECs divide, they differentiate into their destined lineage along the crypt-villus axis^1^. The intestinal epithelium forms a physical barrier between the gut lumen and milieu interior, and as such provides the first layer of defense against harmful luminal components and pathogens^2^. Damage to the intestinal epithelium has been linked to immune activation with a T cell component in multiple disease settings, including inflammatory bowel disease (IBD), coeliac disease, and acute intestinal graft-versus-host disease (GVHD) after allogeneic hematopoietic stem cell transplantation (HSCT)^2–9^. How epithelial damage may directly influence T cell activation still needs to be completely elucidated.

Mechanisms of T cell activation after epithelial damage are thought to involve innate immune cell activation including neutrophils, monocytes and macrophages, leading to migration of T cells, and both local antigen presentation, as well as antigen presentation by professional antigen-presenting cells (APCs) in nearby lymphoid organs^10^. Chemotherapy used in the treatment of malignancies^11, 12^ and conditioning regimens prior to HSCT, damage IECs and disrupt the epithelial barrier^13–15^. After HSCT, the barrier breach may cause local inflammation and activation of T cells supporting the development of GVHD^16, 17^. The type and intensity of the conditioning regimen and related regimen-related toxicity correlate with the development and severity of GVHD, and related outcomes in patients^18–21^.

It is currently unclear how conditioning-induced IEC damage directly affects T cell behavior, and as such could contribute to the development of GVHD. There is evidence that the intestinal epithelium and T cells closely interact under both homeostatic and pathogenic conditions, modulating T cell recruitment, differentiation, and function^22–30^. T cells exhibit dynamic behavior within the IEC compartment which adapts to intraluminal epithelial cell exposure to pathogens^31, 32^. During both homeostatic and inflammatory conditions intestine-derived IL-18 modulates inflammation by suppressing Th17 cells and stimulating T regulatory (Treg) cell differentiation^27, 28^. While Treg cell-derived IL-10 can support Lgr5+ intestinal stem cell (ISC) self-renewal^26^. Gut-directed α4β7-expressing T cells are preferentially recruited to intestinal crypts due to clustering of MAdCAM-1 expression on the endothelium of capillaries in the lower small intestinal crypt region making ISCs prone to damage^30, 33^. Here, both CD4^+^ and CD8^+^ donor T cells cause non-specific intestinal epithelial crypt damage through the secretion of apoptosis-inducing interferon-gamma (IFNγ) in GVHD^33^. IECs have also been shown to play an essential role as APCs in murine GVHD^24^ thereby having the potential to locally activate CD4^+^ T cells. MHC-II expression by IECs is regulated through the presence of IFNγ and was shown to be indispensable for the initiation of lethal acute GVHD in the GI tract^24^.

Until now most studies have focused on murine model systems. The identification of factors influencing these direct interactions between IECs and T cells is relevant in multiple settings, from the development of novel therapeutic strategies for the prevention or treatment of GVHD, to the identification of novel targets for cell- and immunotherapy. To this end it is important to develop ex-*vivo* human model systems to interrogate these interactions. Here, we have investigated the direct effects of intestinal epithelial damage caused by chemotherapy exposure on T cell behavior using a human intestinal organoid-based damage model. Intestinal organoids are 3D epithelial cultures that self-organize from isolated intestinal crypts when supplied with essential growth factors and have the potential to differentiate into all IEC types^34, 35^. We and others have demonstrated organoids to be a relevant ex-*vivo* proxy for studying *in vivo* intestinal epithelial-immune cell interactions^33, 36–39^. Here, we show that chemotherapy-induced epithelial injury increases T cell migration, proliferation and activation. In addition, utilizing this damage model we identify galectin-9 (Gal-9), a beta-galactoside-binding lectin released by the damaged epithelium, to play a role in IEC-mediated modulation of T cell behavior. Furthermore, Gal-9 was detectable in the plasma of clinical HSCT patients, and levels were increased in patients that eventually developed GVHD of the gut. Taken together, this study highlights the potential contribution of chemotherapy-induced epithelial damage to directly promoting T cell activation and trafficking. Furthermore, we have identified Gal-9 as a damage-associated molecule modulating T cell migration, proliferation, and IFNγ production potentially contributing to the pathogenesis of immune disorders affecting the gut.

## Methods

### Please refer to supplemental materials for standard protocols

#### Intestinal organoid cultures

Healthy human small intestinal epithelial organoids were established and cultured as previously described^35^. In short, organoids were generated from biopsies of individuals initially suspected of coeliac disease, but declared free of pathology, and stored in a biobank. All individuals provided written informed consent to participate in the study as approved by the medical ethical review board of the UMC Utrecht (METC) (protocol METC 10-402/K; TCBio 19-489). Organoids (>passage 7) were passaged via single cell dissociation using 1x TrypLE Express (Gibco) and resuspended in Advanced DMEM/F12 (Gibco), 100U/ml penicillin-streptomycin (Gibco), 10mM HEPES (Gibco) and Glutamax (Gibco) (GF- medium), and 50-66% Matrigel (Corning). After plating and Matrigel polymerization, human small intestinal organoid expansion medium (hSI-EM) was added consisting of GF-, Wnt-3a conditioned-medium (CM) (50%), R-spondin-1 CM (20%), Noggin CM (10%), murine EGF (50ng/ml, Peprotech), nicotinamide (10mM, Sigma), N-acetyl cysteine (1.25mM, Sigma), B27 (Gibco), TGF-β inhibitor A83-01 (500nM, Tocris), p38 inhibitor SB202190 (10µM, Sigma), Rho-kinase/ROCK inhibitor Y-27632 (10µM, Abcam, for the first 2-3 days of culture), and Primocin (optional) (100µg/ml, Invitrogen). Medium was refreshed every 2- 3 days. For indicated timepoints, treatment wells received different concentrations of Busulfan (Busilvex or TEVA), Fludarabinephosphate (Aerobindo), Clofarabine (Evoltra or Mylan) and rhIFNγ (R&D systems).

#### T cell activation co-culture assay

For the evaluation of T cell activation and proliferation in the presence of chemotherapy treated organoids, organoids were cultured and treated with the indicated condition for 24h. Subsequently, organoids were mechanically disrupted. A portion of each condition was used to dissociate into single cells to infer the cell number and normalize between different conditions. T cells were isolated and stained with CTV. Co-cultures were set up in a 96-well plate, with 200,000 T cells and the equivalent of 1/6^th^ of untreated organoids per well, in no SB medium with 10% BME. In activating conditions, wells had previously been coated with anti-CD3 and the medium was supplemented with anti-CD28. After 4 days of co-culture, the BME was disrupted in situ and the plate was cooled at 4°C for 30 min, before centrifugation at 500g 5 min at 4°C. When performing intracellular staining, GolgiStop (containing Monensin, BD Biosciences) was added to the medium 4h before staining. The pellets were dissociated to single cells with TrypLE, washed with PBSO and consequently stained for FC analysis as described above. For the Gal-9 blocking assays, 10µg/ml anti-gal-9 mAb (BioLegend) was added to the co-culture from the start of the assay.

#### RNA sequencing

For RNA sequencing of treated organoids, mRNA was isolated using Poly(A) Beads (NEXTflex). Sequencing libraries were prepared using the Rapid Directional RNA-Seq Kit (NEXTflex) and sequenced on a NextSeq500 (Illumina) to produce 75 base long reads (Utrecht DNA Sequencing Facility). Sequencing reads were mapped against the reference genome (hg19 assembly, NCBI37) using BWA41 package (mem –t 7 –c 100 –M –R)^40^. RNA sequencing was analyzed using DESeq2^41^ in the R2: Genomics Analysis and Visualization Platform (http://r2.amc.nl). A principle component analysis (PCA) was performed, and a list of differentially expressed genes (padj<0.1) was generated. Gene Ontology (GO) term analysis was done using either upregulated (logFoldChange>0) or downregulated (logFoldChange<0) DE genes as compared to control per condition using 2X2 contingency table analysis chi-square with continuity correction (padk<0.1). REVIGO software was used to visualize GO term clusters (http://revigo.irb.hr/). Gene Set Enrichment Analysis Pre-ranked analysis was performed with the GSEA software probing for enrichment of genes belonging to Hallmark datasets in the GSEA software^42^. A Venn Diagram was constructed using the webtool provided on (https://bioinformatics.psb.ugent.be/webtools/Venn/) and remade using Biorender.

#### Olink proximity extension proteomic analyses

Organoids were cultured in 24-well plates and treated for 48h with indicated conditions. After treatment, the medium was refreshed with hSI-EM without p38 inhibitor (no SB) for 24h. Consequently, the CM was harvested and centrifuged 1000 g for 5 min. The supernatant was transferred to fresh Eppendorf tubes and stored at -80 °C until analysis. The Olink Target 96 Immuno-Oncology panel (v.3112) from Olink (Uppsala, Sweden) was used to quantify 92 immuno-oncology related proteins in each sample (Supplementary Table 10) Multiplex proximity extension assay panels were used to quantify each protein, as previously described^43^. The raw quantification cycle values were normalized and converted into normalized protein expression (NPX) units. The NPX values were expressed on a log2 scale in which one unit higher in NPX values represents a doubling of the measured protein concentration. Quality control of the measured samples was performed by using the standard quality control protocol of OLINK.

#### Statistical analysis

Data are presented as mean ± S.E.M. To take into account intra-individual and intra-experimental variation experiments were performed at least twice with several wells per condition, and sample material coming from at least two different human donors. Statistical significance was determined at *P ≤* 0.05 using 2-way analysis of variance (ANOVA) with Tukey’s multiple comparison test, 1- way ANOVA with Šidák’s multiple comparison test, row-matched (RM) 1-way ANOVA with Dunnett’s multiple comparison test, a Mann-Whitney *U* test, or a Student *t* test where appropriate. Significance is indicated as *P ≤* 0.05 (*), *P ≤* 0.01 (**), or *P ≤* 0.001 (***) or *P* < 0.0001 (∗∗∗∗).

#### Data sharing statement

For original data please contact:

### Results

#### Modeling chemotherapy-induced damage in human small intestinal epithelial organoids

To study the direct effects of chemotherapy-induced epithelial damage on T cell behavior, we developed a novel damage model by exposing human intestinal epithelial organoids to three chemotherapeutics that are frequently used in HCT conditioning regimens: busulfan (Bu), fludarabine (Flu) and clofarabine (Clo)^44–51^. Busulfan is an alkylating agent that crosslinks DNA strands, inhibiting DNA replication^52^, while fludarabine and clofarabine are both nucleoside analogs (NA), which are incorporated into the DNA during normal synthesis and thereby stall replication^53^. We evaluated chemotherapy concentrations that are used within the range of preclinical studies conducted in leukemia cell lines that eventually formed the basis for their clinical application^45, 54^. Four-day chemotherapeutic exposure resulted in visible morphological changes, with smaller organoid size, a condense/folded phenotype and shedding of dead cells and cell debris both inward into the organoid lumen and outward into the organoid surroundings (**Figure 1a**). Sites of stalled DNA replication and DNA damage can be identified by nuclear foci of phosphorylated (γ) histone (H)2AX complexes^55^. Indeed, exposure to all three chemotherapeutics caused a significant increase in γH2AX complexes per nucleus in treated organoids (**Figure 1b, Suppl. Fig. 1a**). In treated organoids the induced replicative stress correlated with a reduction in proliferation compared to untreated organoids (**Figure 1c, Suppl. Figure 1b**). A dose-dependent decrease in mitochondrial membrane potential was also observed (**Figure 1d**) indicative of increased oxidative stress and apoptosis. Indeed, chemo-exposure induced apoptosis, as measured by a dose-dependent increase of caspase 3/7 activity (**Figure 1e**) and an increased percentage of Annexin-V positive cells (**Figure 1f**). Functionally, the number of organoids generated from single cells was reduced, showing that chemo-treatment impaired their ability to regenerate (**Figure 1g**). Taken together, the tissue-damaging effects of chemotherapeutic-exposure on human intestinal epithelia can be modeled ex-*vivo* using organoids.

**Figure 1.**
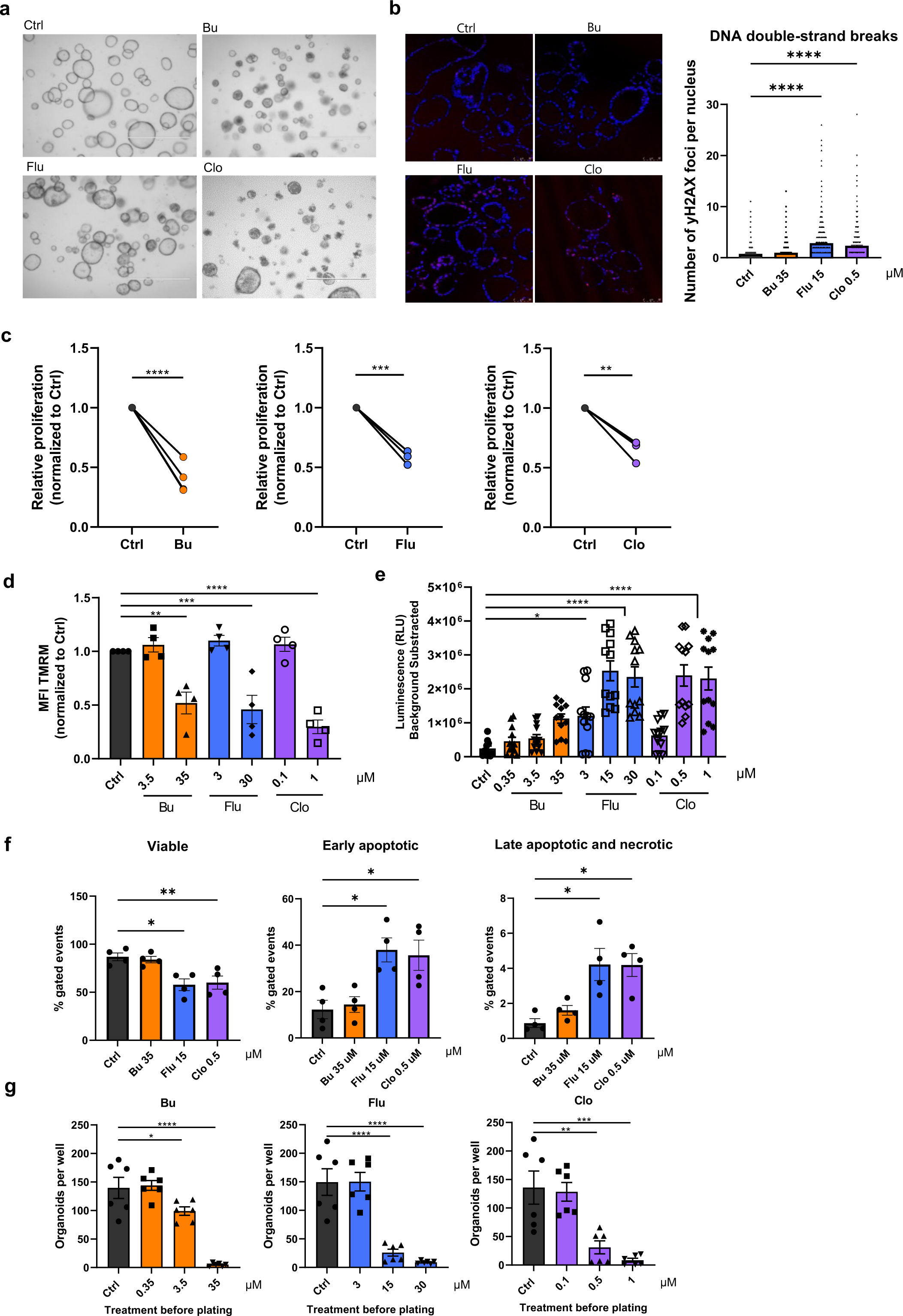
Modeling chemotherapy-induced damage in human small intestinal epithelial organoids. **(a)** Representative EVOS images of organoids treated for 96h with Busulfan (35μM), Fludarabine (15μM) or Clofarabine (1μM), scale bar = 1000μm. **(b)** Representative confocal images of organoids stained with DAPI (blue) and γH2AX-complexes (red/AF647) 24h after chemo- treatment, at 40X, scale bar = 50μm (left panel), quantification of γH2AX foci per nucleus (right panel) (n=2 donors, >700 nuclei analyzed per condition), mean with SEM, ANOVA. **(c)** Relative proliferation of chemotherapy-treated organoids, quantified as CellTrace Violet MFI ratio by FACS after 72h of busulfan 35 μM treatment (n=4 donors), fludarabine 7.5μM (n=2 donors), clofarabine 0.25μM (n=2 donors), Paired t test **(d)** Normalized MFI of TMRM staining for functional mitochondria after 48h of indicated chemo treatment damage, measured by FACS, n=2 donors, mean with SEM, ANOVA. **(e)** Levels of cleaved caspase 3/7 as measured by CaspaseGlo assay after 48h of indicated chemotherapy treatment, n=4 donors, mean with SEM, ANOVA. **(f)** Annexin-V and PI staining for early apoptotic (Annexin-V^+^) and late apoptotic or necrotic cells (Annexin-V^+^PI^+^) after 48h of indicated chemo treatment damage (right panels), measured by FACS. Representative FACS, n=2 donors, mean with SEM, ANOVA. **(g)** Reconstitution of single organoid cells into organoids after treatment with indicated chemo for 48h, n=2 donors, mean with SEM, ANOVA.

#### Chemotherapy treatment modulates intestinal transcriptional programs

To evaluate transcriptional responses to chemotherapeutic exposure, bulk RNA sequencing (RNA-seq) of three independent organoid donors was performed. Treatment of organoids with Bu, Flu or Clo for 24 hours resulted in relatively subtle transcriptional changes with 66, 106 and 118 differentially expressed genes (DEGs) respectively (**Figure 2a, Figure Suppl. 2a**). A concise list of the top 20 most differentially expressed genes identified for each individual treatment and detected in at least two samples is shown in **Figure 2b**. While there are considerable differences between chemotherapeutics, all share certain similarities, in particular the two nucleoside analogs Flu and Clo, reflecting their common mechanism of action (**Figure 2c**, **Suppl. Figure 2b, Suppl. Table 1**). Gene Ontology (GO) term analysis of upregulated genes indicated a similar over- representation of upregulated gene sets in the nucleoside analog Flu- and Clo-treated organoids, including (mitotic) cell cycle processes, metabolic processes, and programmed cell death (**Figure 2d**; a complete list of GO gene sets is provided in **Suppl. Table 2**, and a visual representation is shown in **Suppl. Figure 3a**). Reflecting their mechanism of action, over-representation of genes related to DNA damage and repair as well as the p53 pathway was predominant in Flu- and Clo- treated organoids. Differences between the chemotherapy types, nucleoside analog or alkylating agent, were also observed by Gene Set Enrichment Analysis (GSEA) of chemo-treated organoids (**Figure 2e**). For example, a negative association between Bu-treatment and the p53 pathway was observed, in contrast to Flu- and Clo-treatment. However, a transcriptional change associated with inflammatory response was observed for all treatments. In summary, chemotherapeutics can evoke distinct transcriptional responses in the intestinal epithelium with nucleoside analogs Flu and Clo being more similar when compared with the alkylating agent Bu.

**Figure 2.**
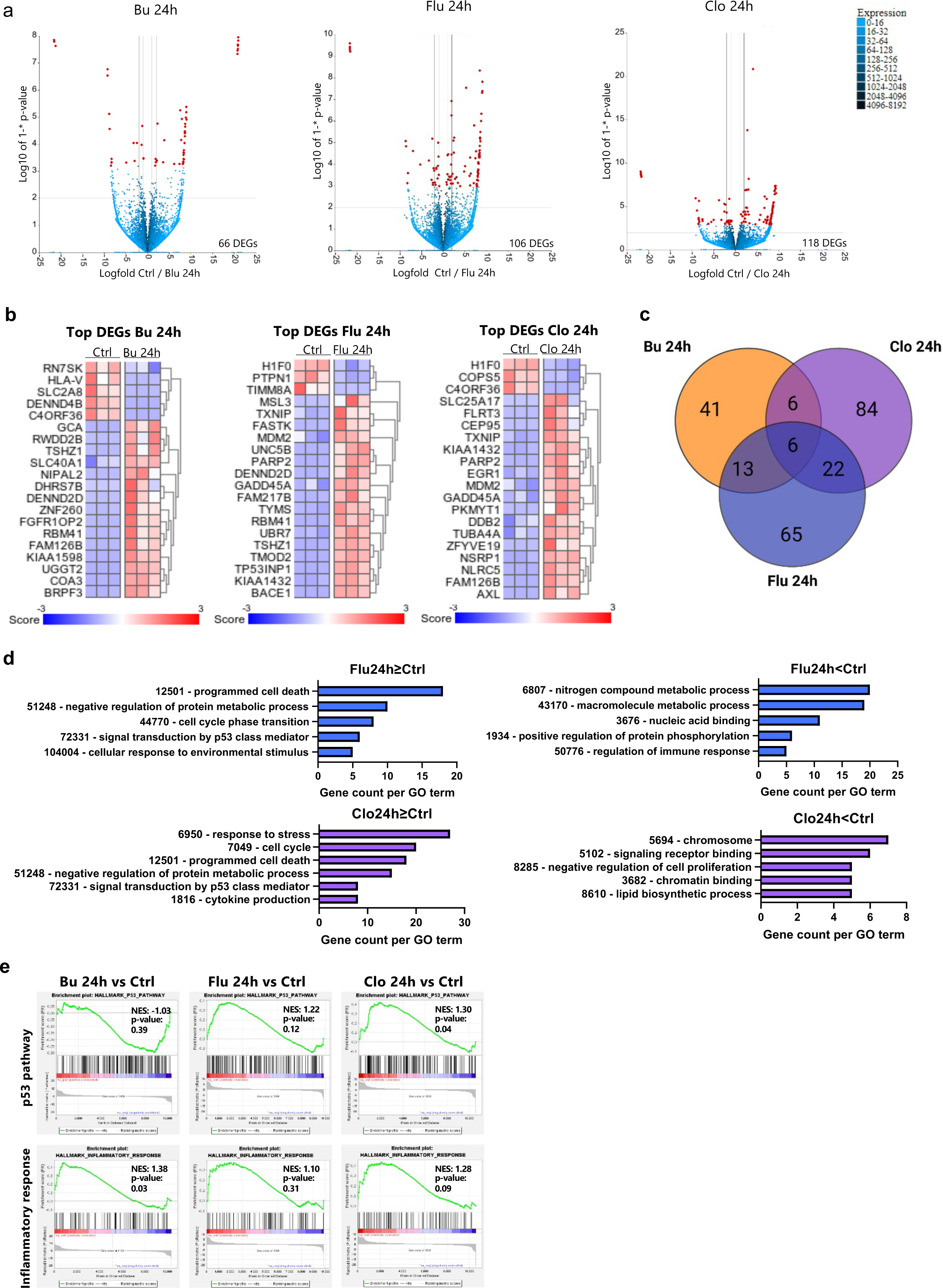
Chemotherapy conditioning specifically reprograms the intestinal epithelial transcriptome. **(a-e)** Human intestinal epithelial organoids (n=3 donors) were treated with busulfan (3.5μM), fludarabine (15μM), clofarabine (0.5μM) for 24h.RNA was isolated and subjected to bulk RNA-sequencing, after which bioinformatics analyses was performed. **(a)** Volcano plots indicating differentially expressed (padj<0.1) genes of 24h Bu- (left), Flu- (middle) and Clo- (right) treated organoids versus control in red. **(b)** Heatmaps of top 20 most different differentially expressed genes in 24h Bu- (left), Flu- (middle) and Clo- (right) treated organoids versus control (DESeq2, padj<0.1). Minimal number of samples containing a present call was set to 2. **(c)** Venn-diagram showing numbers of overlapping and distinct DE genes between different chemotherapeutics (DESeq2, pdaj<0.1). (**d**) Gene Ontology (GO) term analysis of Biological Processes in genes upregulated (LFC>0) or downregulated (LFC<0) by 24h Bu- (top), Flu- (middle) and Clo- (bottom) treatment of organoids (pdaj<0.1). **(e)** Gene Set Enrichment Analysis (GSEA) on all genes in each indicated treatment-condition.

#### Chemotherapy-damaged organoids directly promote T cell activation

To evaluate whether chemotherapy-damaged organoids can directly influence T cell responses, we first evaluated migration (**Suppl. Figure 4a**). Intermediate chemotherapeutic concentrations were chosen to model epithelial damage. In order to quantify T cell migration, we made use of a transwell system with 3µm-pored inserts. Organoids were treated with chemotherapeutics for 48 hours and subsequently cultured in drug-free medium for an additional 24 hours. Either unstimulated or polyclonally pre-activated CTV-stained human peripheral blood CD4^+^ and CD8^+^ T cells were subsequently added to the transwell insert **(Suppl. Figure 4b-c)**. After 24 hours, the number of T cells that had migrated to the lower compartment was evaluated. Both unstimulated and pre-activated CD4^+^ and CD8^+^ T cells demonstrated significantly increased migration towards chemotherapy-treated organoids (**Figure 3a, Suppl. Figure 4d**).

Subsequently, the effect of chemotherapy-damaged epithelium on the polyclonal activation of T cells was evaluated (**Suppl. Figure 4e**). CD4^+^ and CD8^+^ T cells were isolated, stained with CTV, and activated by incubation with plate-bound anti-CD3 and soluble anti-CD28, in the presence of untreated or chemotherapy-treated organoids (representative images of organoids after treatment and before washing are shown in **Suppl. Figure 4f**). This co-culture system allows measurement of the effects of epithelial damage exposure on T cell activation, as T cells received activation stimuli in presence of damaged organoids. Proliferation and activation markers were assessed by flow cytometry after four days of co-culture (**Suppl**. **Figure 4b**). Co-culture with chemotherapy-treated epithelial organoids led to both increased IFNγ-production (**Figure 3b**) and proliferation (**Figure 3c**). However, membrane expression levels of activation markers CD25 and CD69 (**Figure 3d-e**) were not significantly changed between conditions, with the exception of CD4^+^ T cells co-cultured with Clo-treated organoids which expressed higher CD25. In conclusion, Bu-, Flu- and Clo-damaged intestinal epithelium promote T cell migration, and potentiated T cell activation, supporting T cell expansion and IFNγ production.

**Figure 3.**
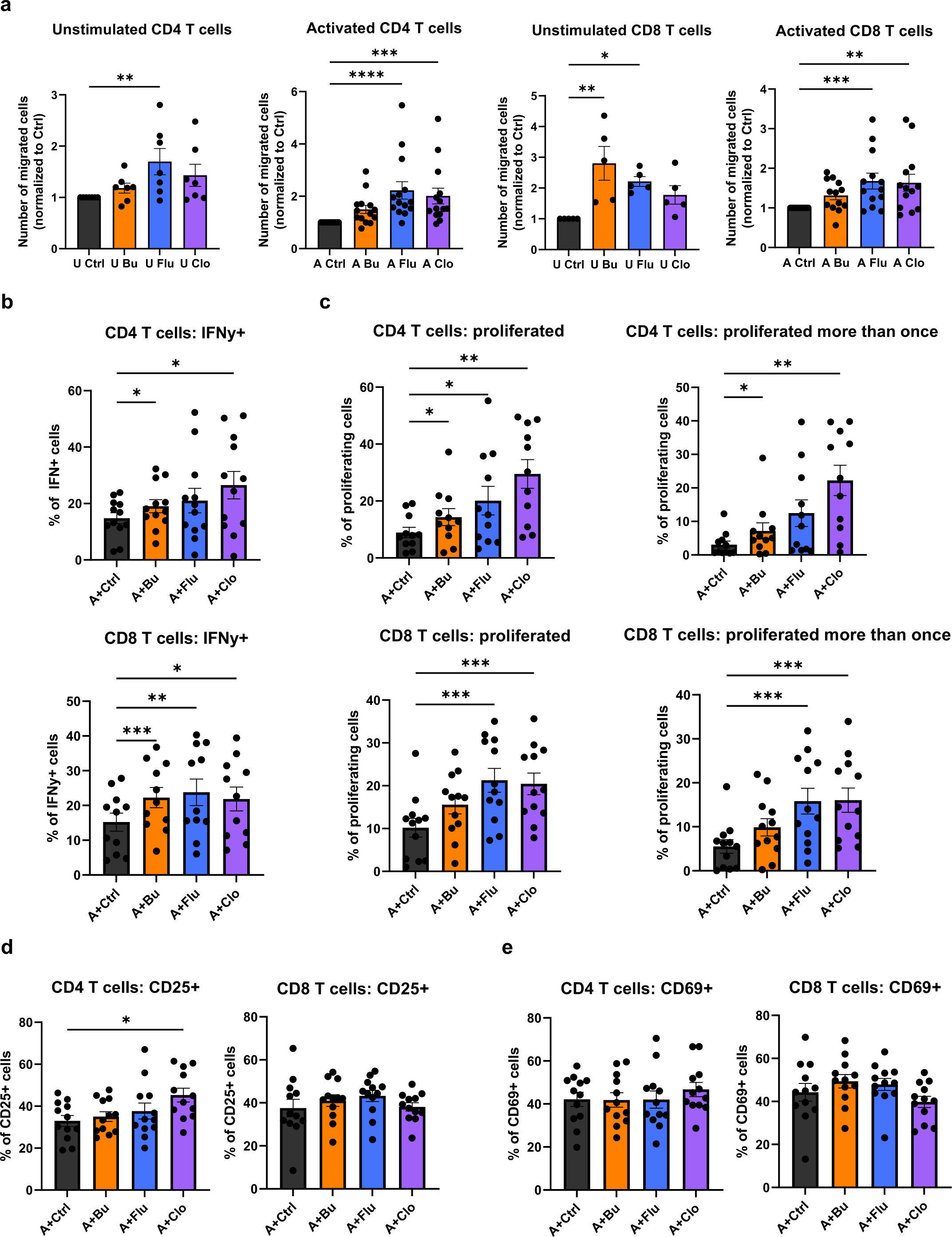
Chemotherapy-induced damage promotes T cells migration and activation. **(a)** Normalized number of unstimulated and pre-activated CD4^+^ and CD8^+^ T cells that have migrated overnight from a 3 μm-pore sized insert (upper compartment) to the lower compartment containing organoids that were treated with busulfan (35μM), fludarabine (15μM), clofarabine (0.5μM) for 48h, and then refreshed for 24h, counted by FACS. n≥5 T cell donors with 1 organoid donor, each data point indicates a T cell donor, mean with SEM, ANOVA. **(b-d)** (Membrane) activation marker expression and proliferation of CD4^+^ and CD8^+^ T cells as measured by CTV- dilution after 4-day co-culture with organoids that were previously treated with busulfan (35μM), fludarabine (15μM), clofarabine (0.5μM) for 24h before the start of the assay. Organoids were washed and disrupted mechanically before replating in co-culture with T cells. n≥11 T cell donors with 1 organoid donor, each data point indicates a T cell donor, mean with SEM, ANOVA.

#### Intestinal organoid-derived galectin-9 modulates T cell migration and activation

To gain mechanistic insight as to how epithelial damage can increase the migration and proliferation of T cells, we utilized the Olink proteomics platform. Here, we evaluated conditioned medium (CM) from both chemotherapy-damaged and untreated intestinal organoids^43^. 44 proteins were above level of detection (LOD), including chemokines such as C-X-C motif chemokine ligand 9 (CXCL9), CXCL10 and CXCL11; cytokines such as IFNγ and IL-8; immune checkpoint molecules such as Programmed death-ligand 1 (PD-L1), and growth factors including EGF among others (**Figure 4a**). In the conditioned media from chemotherapy-damaged organoids the levels of galectin-9 (Gal-9), a beta-galactoside-binding lectin, were increased in CM predominantly from Flu- and Clo-treated organoids (**Figure 4b**). To determine whether Gal-9 may play a role in mediating the effects of chemotherapy-damaged organoids on T cell responses, an anti-Gal-9 blocking monoclonal antibody was utilized. Anti-Gal-9 added to the lower compartment significantly inhibited CD8^+^ T cell migration and showed a trend towards decreased migration of CD4^+^ T cells (**Figure 4c**). Furthermore, the presence of anti-Gal-9 abrogated both increased proliferation (**Figure 4d**) and IFNγ levels (**Figure 4e**) observed in the presence of Bu-, Flu- and Clo-treated organoids. In contrast, T cells preserved the ability to express activation markers CD69 and, to a certain extent, CD25 (**Suppl. Figure 5a**). These data suggest that Gal-9 released by chemo-damaged epithelium can play an important role in T cell migration towards chemotherapy-damaged epithelium, consequently potentiating T cell activation supporting both expansion and IFNγ production.

**Figure 4.**
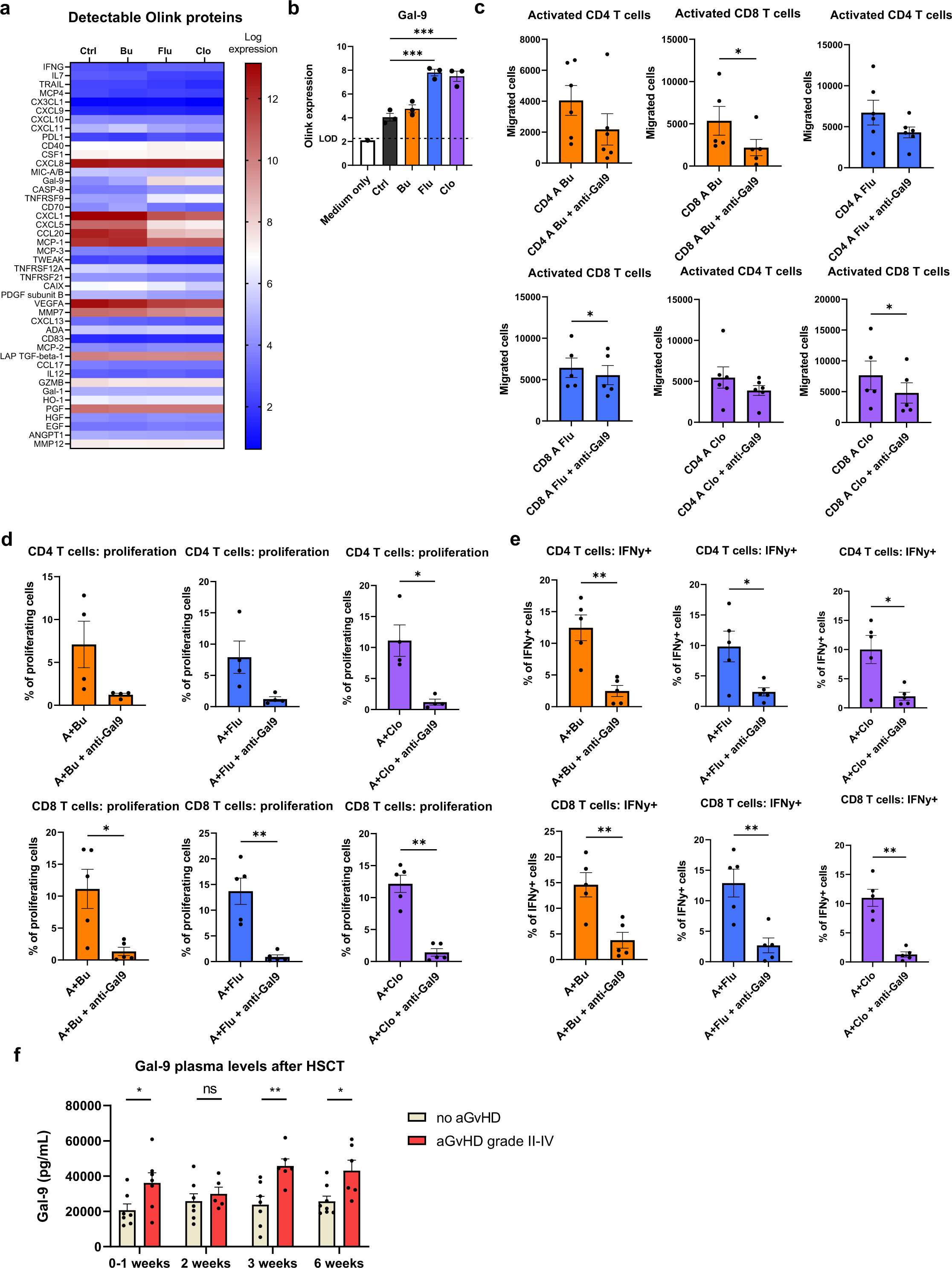
Galectin-9 is released by chemo-damaged organoids and contributes to T cell migration, activation and proliferation. **(a-b)** Analysis of proteins present in conditioned media of organoids treated with chemotherapeutics as in the ex-*vivo* T cell migration assay (Olink Proteomics) (n=3 donors). Each data point indicates the levels of Gal-9 in the CM from each organoid condition (log scale), mean with SEM, ANOVA. **(c)** Migration of activated CD4^+^ and CD8^+^ T cells towards treated organoids in the presence of anti-gGal-9 blocking mAb. n≥5 T cell donors with 1 organoid donor, each data point indicates a T cell donor, mean with SEM, paired t-test. **(d)** CD4^+^ and CD8^+^ T cell activation after co-culture with treated organoids in the presence of anti-Gal-9 mAb. n≥4 T cell donors with 1 organoid donor, each data point indicates a T cell donor, mean with SEM, paired t-test. **(f)** Galectin-9 levels in plasma of HSCT patients with and without acute grade II-IV (gut) GVHD as measured by Luminex. n≥5 patients per condition.

To evaluate whether Gal-9 levels following conditioning are also increased in a clinical setting, we measured Gal-9 levels in stored plasma samples of 17 pediatric transplant patients aged 12-18 at multiple time points after HCT. All patients received a conditioning regimen with Bu and Flu, and, in addition, some patients Clo, before undergoing transplantation with a cord blood graft. Seven patients developed acute GVHD grade II-IV with gastrointestinal involvement. Median time to GVHD was 24 days (range 15-45) and all patients had a follow-up of more than 100 days. Gal- 9 levels were measurable in the plasma of all patients (**Figure 4f**) and were increased at the day of, or early after, transplant in patients that later on developed grade II-IV acute GVHD. Furthermore, Gal-9 levels remained elevated at several time points after HCT in grade 2-4 GVHD patients. While correlative, this observation suggests that our model system can have predictive value in identifying novel intestinal damage biomarkers. Taken together, our data suggests that Gal-9 may be a biomarker for the development of GVHD in HCT patients. Our functional data further support a pathogenic role for Gal-9, perhaps by increasing migration and activation of T cells at the site of the conditioning-damaged intestinal epithelium.

## Discussion

Immune activation after intestinal epithelial cell injury is a well-known phenomenon, but how such damage can directly affect T cell function remains unclear. Here we have developed a novel ex- *vivo* intestinal organoid model to mimic chemotherapy-induced damage which can occur during cancer treatment or conditioning preceding HSCT. This ex-*vivo* human model has allowed us to study the direct effect of epithelial damage on T cell homeostasis, without possible confounding or species-specific interactions that may occur in an *in vivo mouse* setting. In addition to developing our understanding of the biology of T cell responses during sterile inflammation, the model has the potential to serve as a platform for the discovery of new targets and testing of therapeutics in multiple intestinal disease settings including the field of HCT and GVHD, as well as in the tailored development of autologous immunotherapy and CAR T cell therapy for solid tumors in sequence with chemo or radiotherapy^56–59^.

This is the first study comparing epithelial damage response to different clinically relevant chemotherapeutics. We observed that exposure of intestinal epithelial organoids to chemotherapeutics resulted in chemotherapy-distinct transcriptional responses. Their main chemotherapeutic mechanism of action is chain termination when incorporated in place of natural purine nucleosides, with stalling of the cells in S-phase resulting in induction of apoptosis. Clofarabine affects cells with lower proliferation rates as well. Busulfan on the other hand is an alkylating agent that crosslinks guanine bases in the DNA, therefore, making it impossible for DNA strands to unfold, thereby also resulting in apoptosis. Although we cannot completely exclude the contribution of the chemotherapeutic concentrations utilized, we observed a similar gene expression pattern and related T cell responses between Flu and Clo, the purine nucleoside analogs we included in our study. While it is currently not possible to reduce chemotherapy regimens, it is important to understand the differential consequences of exposure to each individual chemotherapeutic, and a further argument for pursuing the development of antibody-based conditioning for the future^60^.

We found that epithelial damage caused by chemotherapy increases the migration of both resting and polyclonally pre-activated T cells toward the site of injury. The migration assay studies the cells in two compartments, suggesting a soluble molecule gradient to be responsible for this. Interestingly, in our setting, migration occurred in the absence of endothelium, or any previously known intestinal-epithelial-derived chemokine. In particular, we did not observe an increase in the concentration of the chemokines CXCL9, CXCL10 and CXCL11. However, we observed decreased T cell migration when blocking Gal-9 during the migration assays, suggesting a possible new role for Gal-9 in modulating T cell trafficking to the damaged epithelium.

Besides the capacity of chemotherapy-induced epithelial damage to increase T cell migration, we also demonstrate a direct influence on the proliferation and activation of T cells. Besides peripheral and local hematopoietic APC-mediated activation, we have shown for the first time that damaged epithelium can directly additionally stimulate tissue-recruited activated T cells, which likely further propagates intestinal damage through local IFNγ production. While the precise mechanism by which this occurs remains to be fully elucidated, we have identified Gal-9 as novel driver. Gal-9 is a beta-galactoside-binding lectin with immunomodulatory properties widely expressed by a variety of tissues^61–63^. The carbohydrate domains of Gal-9 can bind ý- galactosides, such as lactose, found on O-glycans and N-glycans of glycosylated proteins or lipids^64^. Gal-9 expression can be nuclear as well as cytoplasmic, or extracellular where it is membrane-bound or in the extracellular matrix. It has many different receptors, among which TIM- 3, DR3, 4-1BB, CD44, PD1 and Protein Disulfide Isomerase are expressed on T cells ^65–70^. Interestingly, the expression of soluble TNFRSF9 (CD137 or 4-1BB), one of the receptors for Gal- 9, was also significantly upregulated in chemo-treated organoids (**Supp. Figure 5b**). The reported effects of Gal-9 on T cells *in vitro*, *in vivo,* and deduced from human biomarker studies are pleiotropic and range from immunosuppressive to pro-inflammatory. The complexity of reported effects may be related to the Gal-9 doses utilized, species differences, as well as the cellular or tissue context. Gal-9 at low doses has been shown to increase T cell expansion and activation ex-*vivo*, inducing a Th1 phenotype with IFNγ production^71^. This Th1-type response by Gal-9 has been linked to a pathway mimicking antigen-specific activation of the TCR resulting in cytosolic calcium mobilization^72–74^. However, which receptor mediates these effects remains unknown. The binding of Gal-9 to TIM3 on conventional T cells has been reported to induce apoptosis^65^, which we have not observed in our studies. However, Gal-9 TIM-3-induced apoptosis has only been observed under supraphysiological concentrations of Gal-9, and the relevance of this for human T cells is unclear. These effects are also likely to be immune context-dependent since CD4^+^ T cells from rheumatoid arthritis patients also do not undergo apoptosis when exposed to increasing concentrations of Gal-9, compared to cells from healthy controls^75^.

Damage to the gastrointestinal tract plays a major role in the morbidity and mortality associated with GVHD. Currently, it remains difficult to predict the onset of intestinal GVHD due to a lack of early biomarkers. Our data show increased plasma levels of Gal-9 early after conditioning which persist in patients that develop GVHD after HCT. Gal-9 levels have also been associated with intestinal inflammation and correlate with disease severity in inflammatory bowel disease^76^. Gal- 9 levels are also increased in patients with atopic dermatitis^77^. In contrast to our own data, a recent study has shown that Gal-9 intraperitoneal administration reduced acute GVHD in a murine model^78^. However, in this study Gal-9 was administered only after the initial onset of GVHD and of note there was no improvement in overall survival. Despite the differences often observed between species, a murine study showed that perturbation of TIM3/Gal-9 interaction increased GVHD lethality, but reduced it when Treg cells were concomitantly depleted from the graft, resulting in improved survival^79^. Gal-9 has also recently received attention in the context of cancer immunotherapy^80, 81^ where anti-Gal-9 blocking antibodies are being developed as checkpoint inhibitors to induce the anti-tumor immune response. Ongoing clinical trials (NCT0466668) will provide important information regarding the safety and effectiveness of Gal-9 blockade, potentially helping the translation of Gal-9 targeting intervention for GVHD patients^82^. However, given the pleiotropic and diverse effects reported, strict control of Gal-9 perturbation must be in place in respect to timing, location and cell types involved.

In conclusion, we have modeled chemotherapy-induced damage to the human intestinal epithelium and studied its effects on T cell homeostasis. T cells demonstrate increased migration to chemotherapy-treated organoids, proliferated more and expressed higher levels of IFNγ. We propose that damaged-organoid-derived Gal-9 may be a novel damage-associated molecule responsible for these effects, and we suggest that Gal-9 may be a possible biomarker for GVHD development in HCT patients. In addition, treatment aimed at blocking/potentiating Gal-9 may be a new damage-dampening, preventive therapy in the context of immune-mediated diseases in which there is damage in the gut, such as GI-GVHD.

## Acknowledgments

We would like to thank all members of the Lindemans and Coffer groups for helpful discussions. Elsbeth van Liere, Bart Westendorp, Thomas Brand and Jet Segeren for help with the yH2AX stainings and analysis. Sabine Middendorp for generation of organoid lines. Maaike de Vries for help in acquiring yH2AX confocal images. We thank the Hubrecht Institute FACS facility, the Olink facility at the UMC Utrecht, Noortje van den Dungen of the Epigenetics facility of the UMC Utrecht, USEQ, and Richard Volckmann from R2- support. We would also like to thank Nienke Vriesekoop for discussions on immune cell migration.

Funding: Alexander Suerman Stipend UMC Utrecht; NWO Aspasia grant; WKZ foundation grant, Marie S. Curie Co-fund RESCUE grant No 801540; NIH 2020 R01 HL145631 and 2018 R01HL146338.

## Author Contributions

SAJ and AC designed, performed, and analyzed experiments, and wrote the manuscript. MH, LSS, CK and SN performed experiments. MM assisted with RNA sequencing analysis. EM assisted with experimental design. PJC and CAL supervised the research, helped in experimental design, and wrote the manuscript. EES and AMH provided expert advice. All authors contributed to the manuscript.

## Supplemental Methods

### T cell isolation and activation

T cells were isolated from peripheral blood of healthy donors in the UMC Utrecht as approved by the METC (protocol 07/125) or from buffy coats (Sanquin, NL). After Ficoll-Paque (GE Healthcare) gradient separation, CD8^+^ T cells were isolated from the peripheral blood mononuclear cells (PBMCs) in MACS buffer (2% heat-inactivated FBS, 2% 0,1M EDTA in PBSO) using the CD8^+^ Dynabead isolation kit (Thermo Fisher) and BD IMag Cell Separation Magnet (BD Biosciences). CD4^+^ T cells were isolated from the CD8-depleted PBMC fraction using the MagniSort human CD4^+^ T cell enrichment kit (Thermo Fisher). T cell purity was checked by flow cytometry (routinely >80%, Suppl. Figure 4). T cells were activated using plate-bound functional grade anti-human CD3 (1.6μg/ml in PBSO overnight at 4°C or 2h at 37°C, eBioscience) and soluble functional grade anti-human CD28 (1μg/ml, eBioscience) for 3 or 4 days as indicated at a concentration of 1 million cells/ml in T cell medium (TCM) (RPMI Medium 1640+GlutaMAX-I, Gibco, with 100U/ml pen-strep and 10% heat-inactivated FBS).

### Migration transwell assay

Organoids were cultured in 24-well plates and treated for 48h with indicated conditions. After treatment, the medium was refreshed with hSI-EM without p38 inhibitor (no SB) for 24h. Simultaneously, isolated T cells were activated for 3 days or left resting in TCM. At the start of the assay, the T cells were stained with CTV and added to 3μm-pored transwell inserts (Greiner Bio-One) (400.000 T cells in 200μl) that were placed in the wells with organoids. After overnight incubation, inserts were removed and the contents of each well was dissociated with TrypLE and reconstituted in 300μl. The number of CTV+ events per 150μL sample was counted using flow cytometry. For the Gal-9 blocking assays, 2μg/ml anti-Gal-9 mAb (BioLegend) was added to the lower compartment prior to start of the assay.

### Immunofluorescent stainings

For the γH2AX-staining of organoids, treated organoids were harvested, washed in PBSO and fixated in formalin 4%/eosin 0.1% for 1h at room temperature and transferred to 70% ethanol. Consequently, the organoids were embedded in agarose (2.5%, Eurogentec, EP-0010-05), processed (Leica ASP 300 S) and embedded in paraffin (Surgipath Paraplast, Leica). The FFPE organoids were sectioned at 4μm, dried at 55 °C overnight and then deparaffinized in xylene and rehydrated in decreasing concentrations of alcohol (by using the leica autostainer). For antigen retrieval, the slides were incubated in sodium citrate buffer (10mM, pH 6, Merck) and washed with PBS/Tween20 (PBST). The slides were blocked with normal goat serum (10% in PBST) for 30 min and incubated with rabbit anti-phospho-histone H2A.X (Ser139) (20E3) (1:200, Cell Signaling, 9718) overnight at 4°C. The slides were washed with PBST and then incubated with goat anti-rabbit Alexa 647 (1:200, Thermo Fisher, a21244) for 1h at RT. After additional washing, the slides were mounted with fluoroshield with DAPI mounting medium (F6057, Sigma Aldrich).

### Imaging of organoids

Bright field (co-culture) images were acquired using an EVOS FL Cell Imaging System (Thermo Fisher Scientific). Fluorescence images were generated with a Leica (Wetzlar, Germany) SP8X laser-scanning confocal microscope.

### Quantification of **γ**H2AX-staining

For the quantification of γH2AX-staining in organoids, images acquired by the confocal microscope were processed in Fiji (ImageJ 1.53q) using a script^1^. In short, the channels of the RGB images were split, nuclei defined based on DAPI staining and maxima in the γH2AX channel measured and quantified per nucleus. The number of foci per nucleus has then been analyzed.

### CaspaseGlo assay

Organoids were cultured in 96-well plates and treated with indicated conditions for 48h. Subsequently, the Matrigel was dissolved with GF- and the samples were transferred to a white opaque 96-well plate. Caspase-Glo 3/7 Reagent (Promega) was prepared as per protocol and added to the organoids in a 1:1 ratio to GF- up to a total volume of 100μl. The assay was incubated for 40 min and luminescence was measured with a TriStar2 Multimode plate reader LB942 (Berthold Technologies).

### Flow Cytometry

T cells were stained with live/dead marker Zombie NIR (Biolegend) or Fixable Viability Dye eFluor 780 (Affymetrix eBioscience) and directly conjugated antibodies anti-CD3- PE and anti-CD4- or anti-CD8-FITC (BioLegend) either in FACS buffer (PBSO, 2mM EDTA, 0.5% BSA, Sigma) or MACS buffer (PBS0, 2mM EDTA, 2% FBS). For assessing T cell activation, anti-CD25-APC (BioLegend), anti-CD69-BV605 (BioLegend) were added to the staining. Intracellular IFNγ-staining was performed using the Intracellular Fixation & Permeabilization Buffer Set (eBioscience Thermo Fisher) with anti-IFNg-PECy7 (BD Biosciences). For analysis of proliferation T cells were stained before start of the assay with CellTraceViolet (CTV) (Invitrogen, 5 µM in PBSO) according to manufacturer’s protocol. A Dead Cell Apoptosis Kit with Annexin V-FITC and propidium iodide was used according manufacturer for Annexin V FACS staining. FC data were acquired with a BD LSRFortessa Cell Analyzer (BD Biosciences) using FACSDiva (BD Biosciences) software and a CytoFLEX Flow Cytometer (Beckman Coulter) with CytExpert software. The data were analyzed with FlowJo (Treestar, 10.6.2) or CytExpert software (2.4).

### CellTraceViolet proliferation assay

For evaluation of the effect of chemotherapy treatment on organoid proliferation, organoids were dissociated into single cells and stained with CTV before plating. After 5 days of culture, organoids were harvested, processed into single cells, stained with Zombie NIR, and analyzed by FC.

### Mitochondrial damage assay

For the quantification of mitochondrial damage in chemotherapy treated organoids, organoids were incubated with MitoTracker Green FM (100nM, ThermoFisher) and Tetramethylrhodamine methyl ester perchlorate (TMRM) (150nM, ThermoFisher) for 45 min at 37 °C after treatment. After incubation organoids were dissociated into single cells, stained with Zombie NIR and analyzed by FC.

### Luminex

Biobanked plasma samples from HSCT patients at specific time points related to their HSCT date were used for luminex analysis. The data were collected after patients provided written informed consent (HSCT Biobank, local IRB approval 05-143 and 11- 063k) in accordance with the Helsinki Declaration. Plasma had been stored at -80°C until analysis. With Luminex multiplex immunoassay technology, a total of 60 plasma proteins were measured; Gal9, IL1RA, IL2, IL3, IL4, IL5, IL6, IL7, IL10, IL15, IL17, IL18, IL22, TNFα, IFNα, IFNψ, APRIL, OSM, LAG3, Follistatin, I309, MIP1a, MIP1b, IL8, MIG, IP10, BLC, OPG, OPN, G-CSF, M-CSF, GM-CSF, SCF, HGF, EGF, AR, VEGF, CD40L, sPD1, FASL, IL1R1, IL1R2, ST2, TNFR1, TNFR2, sIL2Rα, sCD27, IL7Rα, sSCFR, Elastase, S100A8, Ang1, Ang2, LAP, TPO, sICAM, sVCAM, MMP3, Gal3, C5a. The multiplex immunoassay was performed according to the protocol from the MultiPlex Core Facility of the UMCU^2^.

### Illustrations

All illustrations were made with BioRender.com.

## Supplementary Figures

**Supplementary Figure 1.**
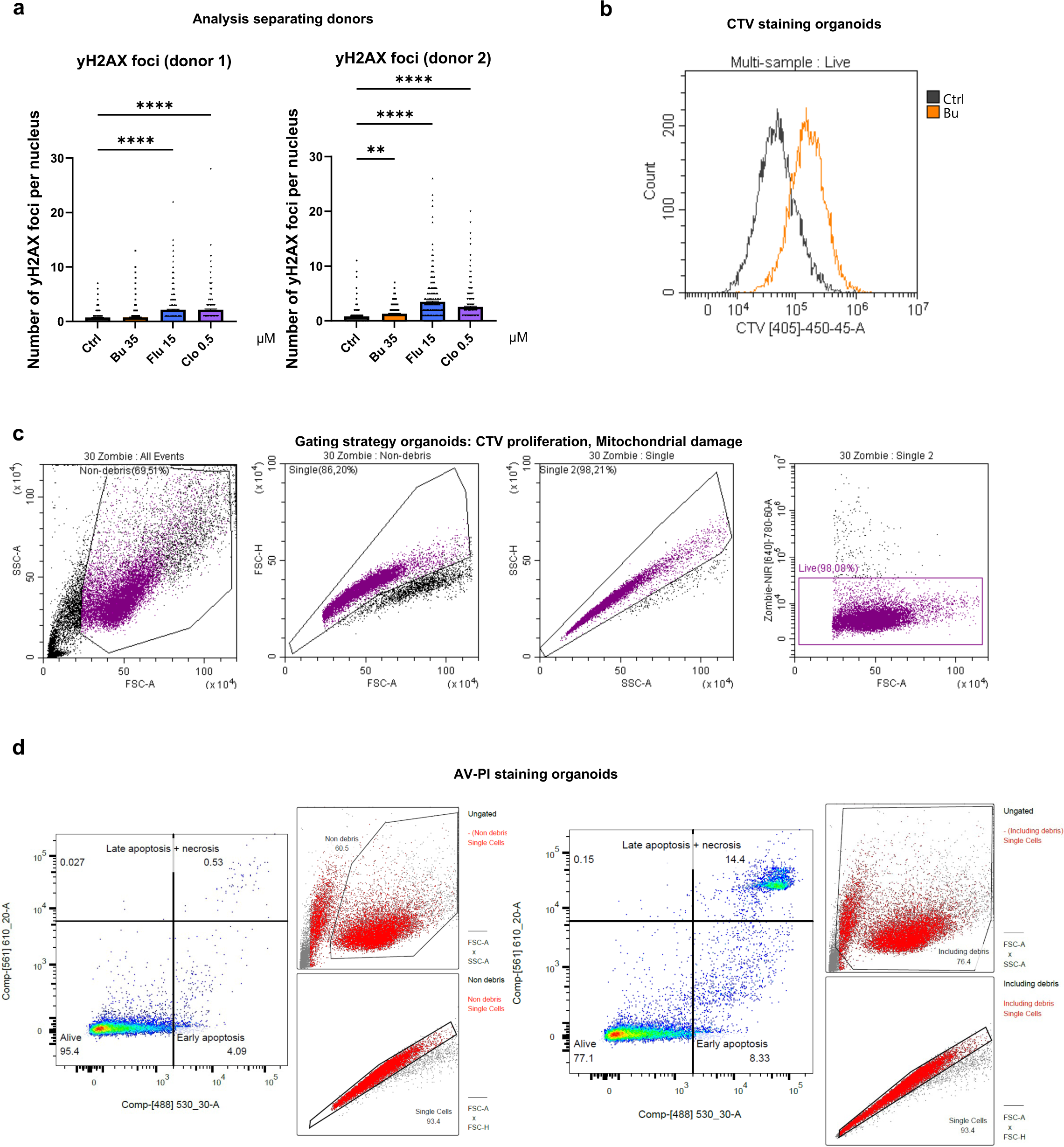
**(a)** Quantification of yH2AX foci per nucleus per donor. **(b)** CTV signal by FACS of organoids on day 5 of culture after treatment with Bu for 4 days. **(c)** Gating strategy of single organoid cells in flow cytometry, as used in Figure 1c and 1d **(d)** Gating strategy for Annexin-V-PI staining to indicate viable, early and late apoptotic/necrotic cells by FACS. The right panel includes debris characterized by a low FSC-A and high SSC-A, included here to illustrate gating decisions. The analysis of apoptotic cells has been conducted excluding debris as depicted in the left panel.

**Supplementary Figure 2.**
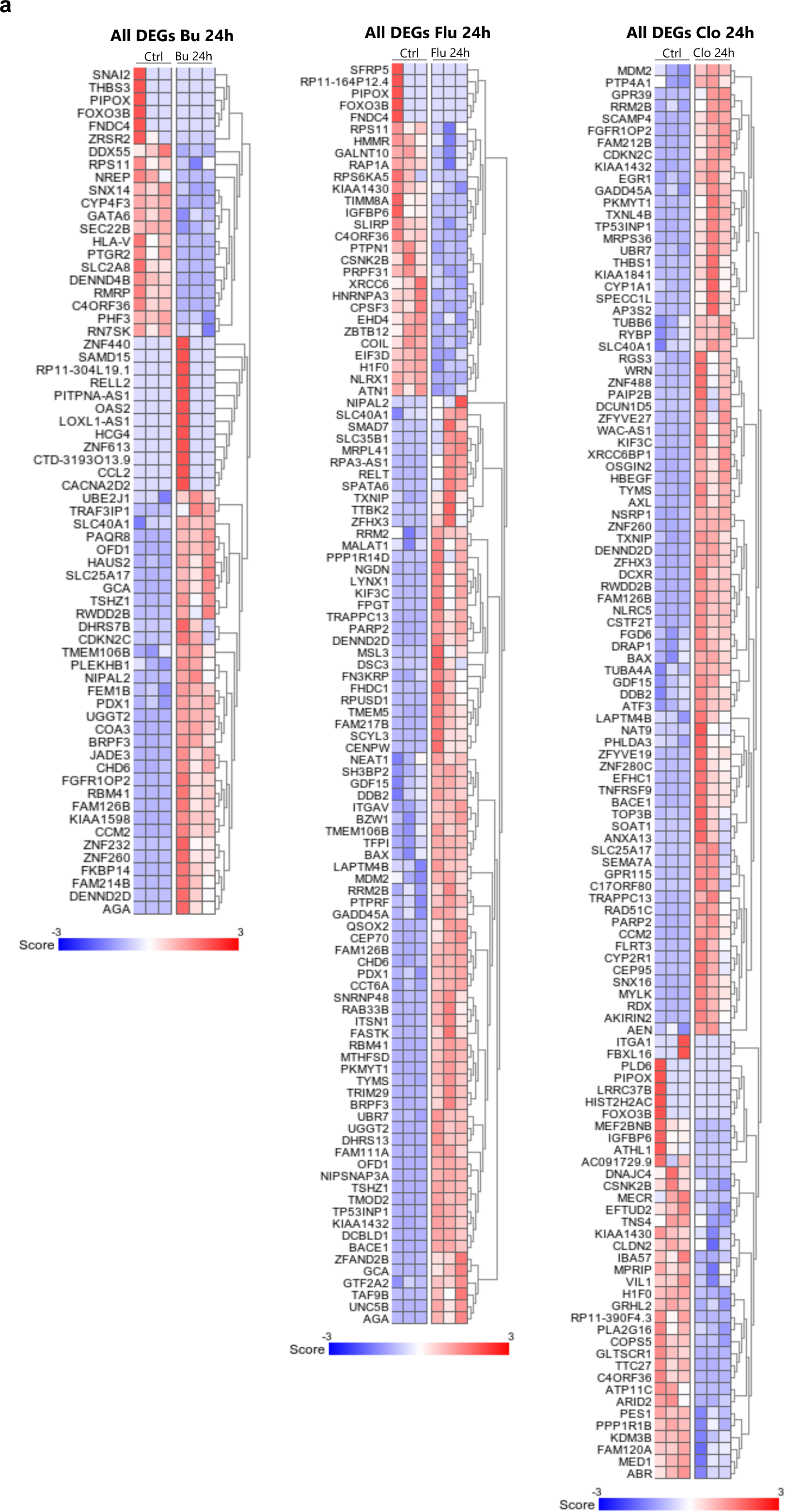
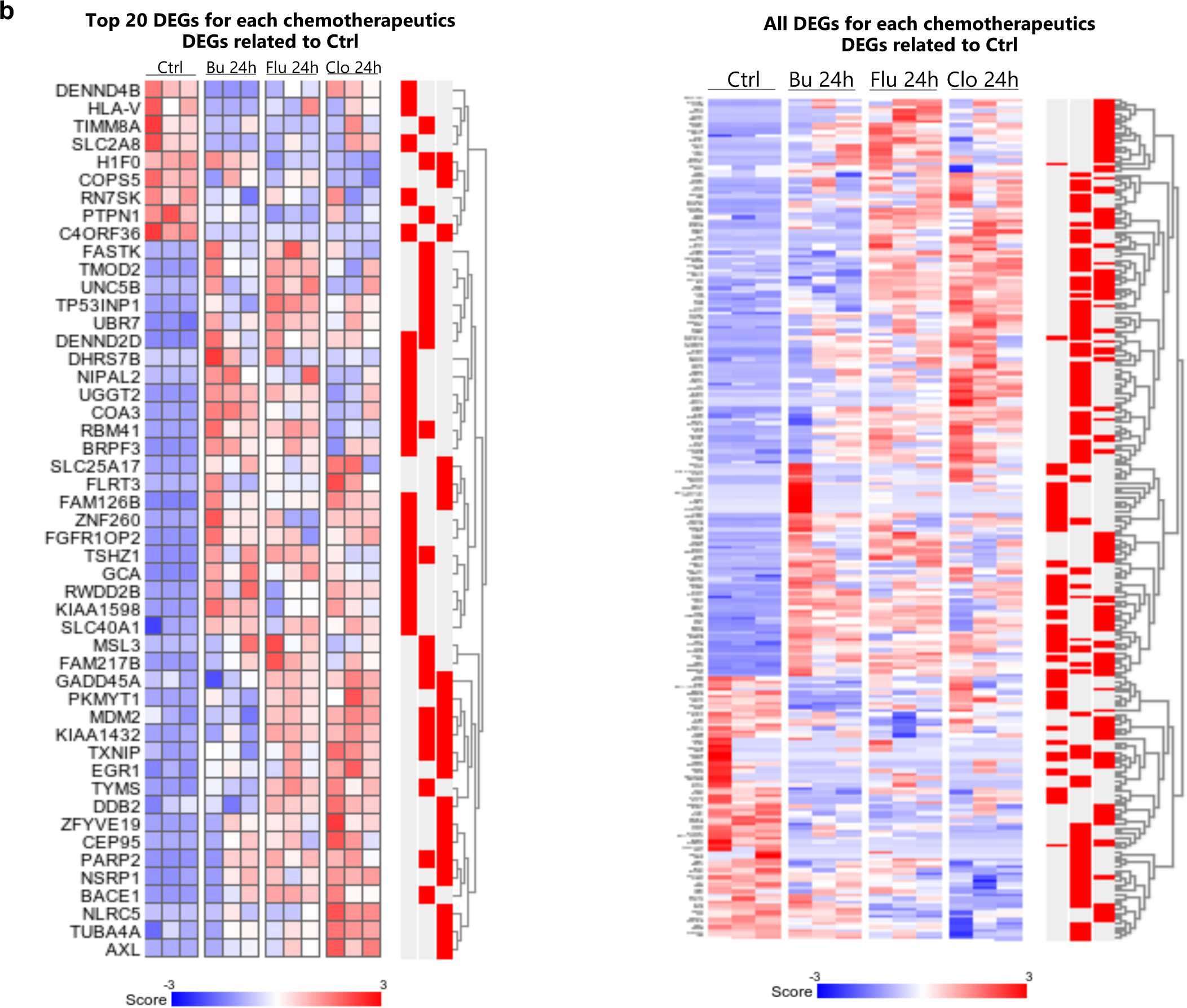
**(a)** PCA clustering of chemo-treated samples, indicating sample condition (left) and sample donor (right). **(b)** Heatmaps of all different differentially expressed genes (DEGs) in 24 hour Bu- (left), Flu- (middle) and Clo- (right) treated organoids versus control (padj<0.1). **(c)** Heatmaps of a combined list made from the top 20 DEGs of 24 hour Bu-, Flu- and Clo- treated organoids versus control showed in Figure 2b (left) or all DEGs (right).

**Supplementary Figure 3.**
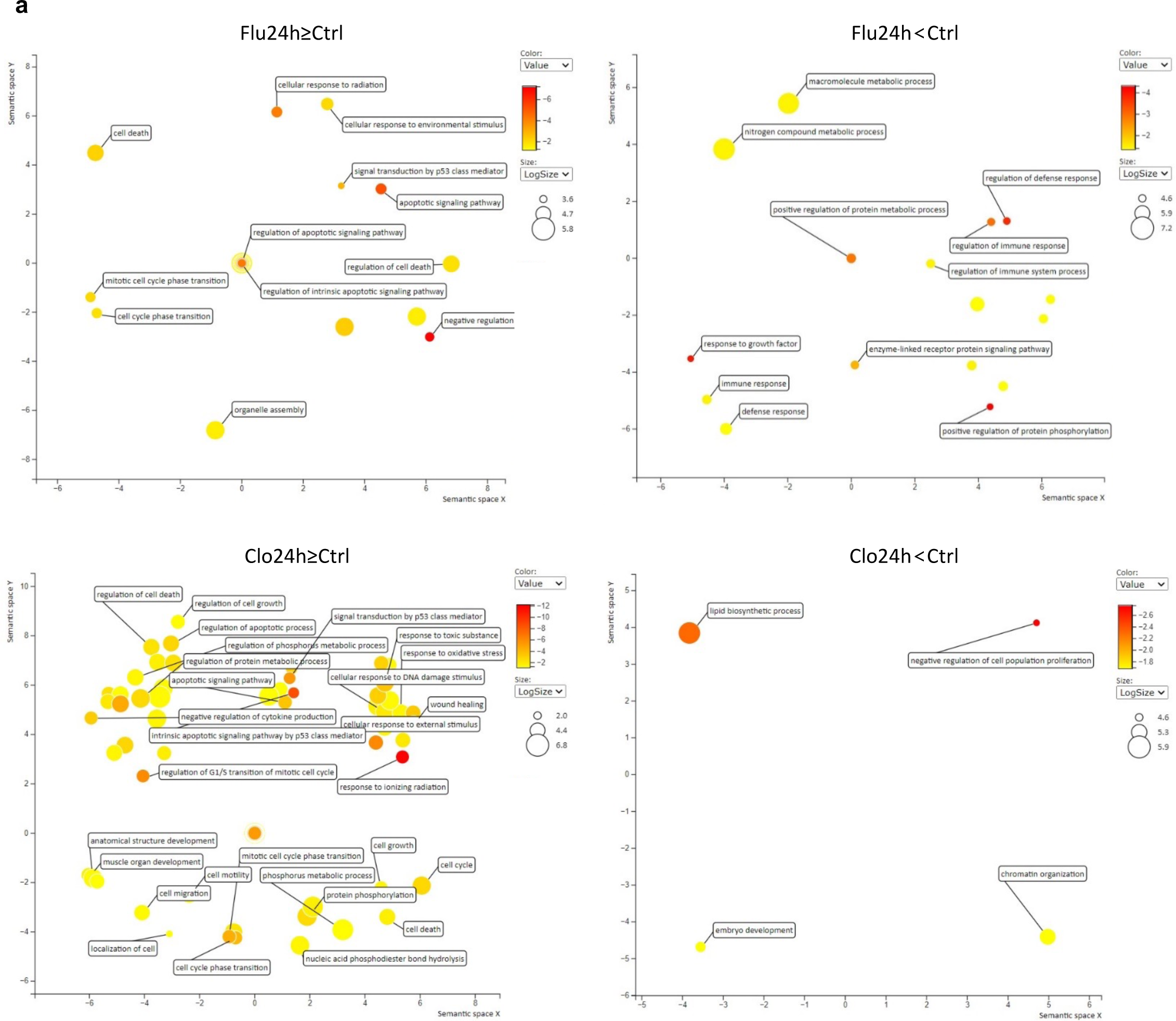
(a) REVIGO analysis of GO gene sets up- or downregulated compared to control obtained from R2 analysis (padj<0.1) reported in Suppl. Table 2.

**Supplementary Figure 4.**
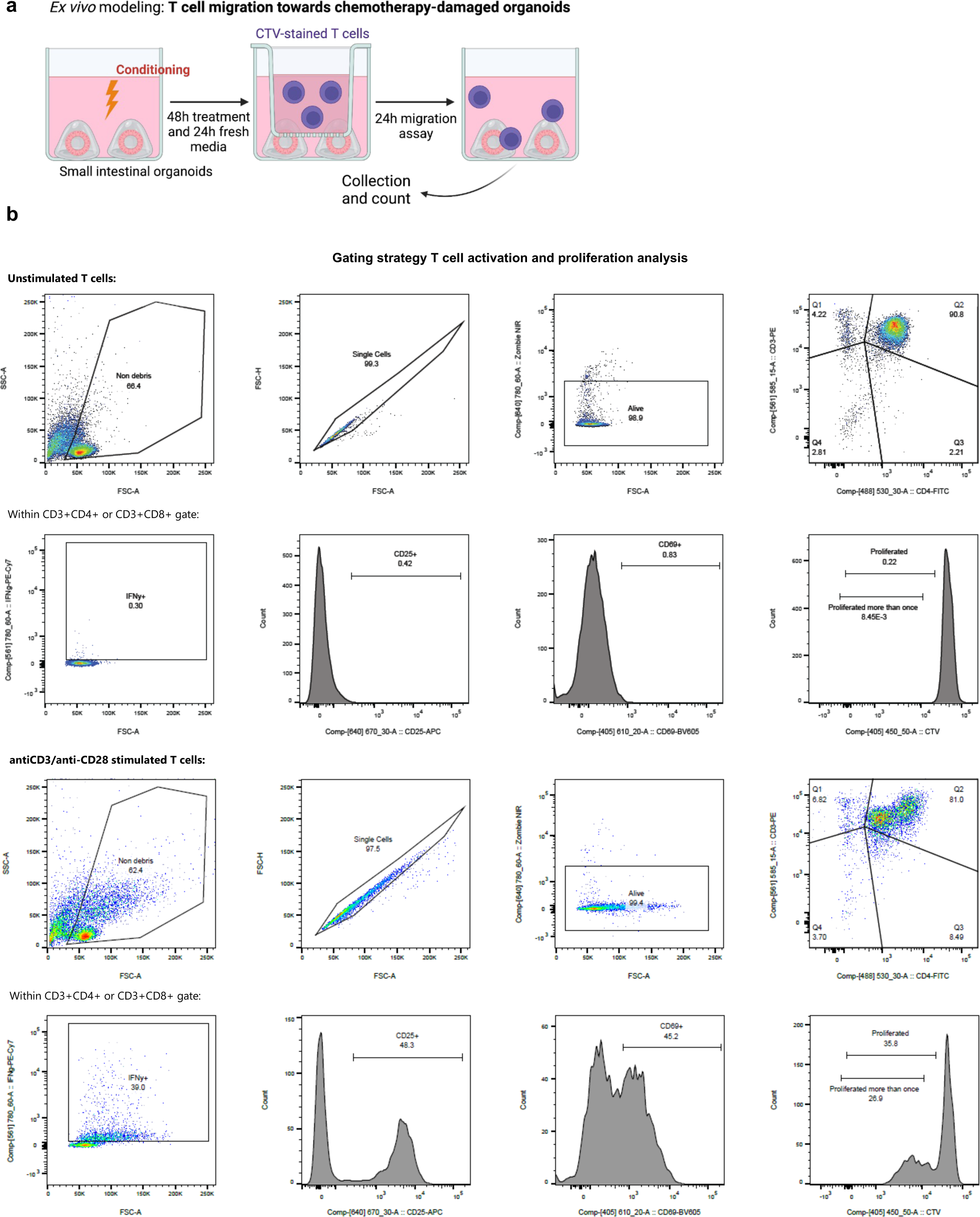
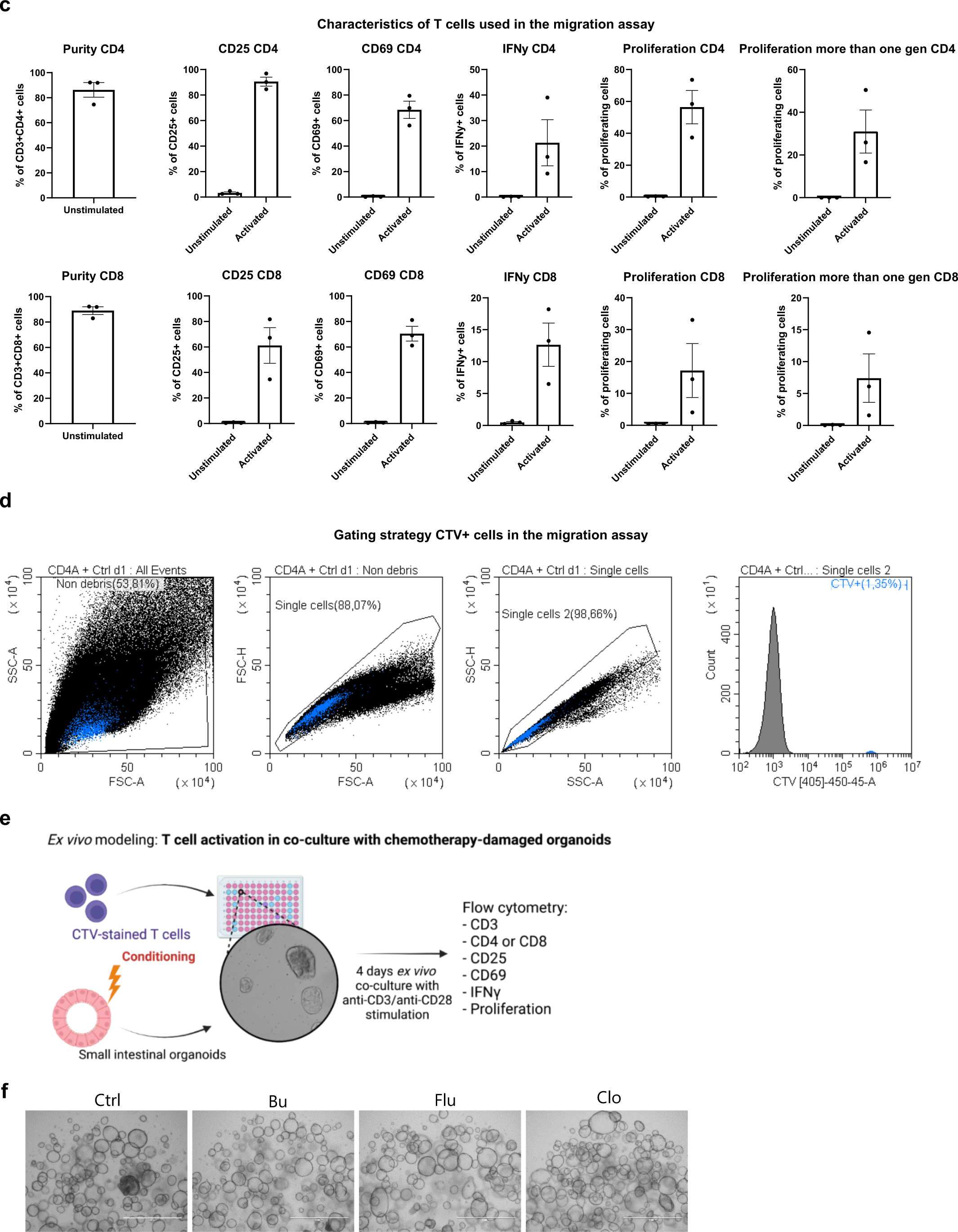
**(a)** Schematic overview of the migration assay, with organoids cultured in the lower compartment and CTV-stained T cells added in the upper compartment of a transwell 3μm-pore sized insert system. Organoids were treated for 48 hrs with chemotherapeutics and cultured additionally in fresh drug-free media for 24 prior to assay start. 24 hrs after the migration assay, cells were collected and CTV+ cells were counted. **(b)** Gating strategy of T cell proliferation and activation markers (example is CD4^+^ T cells), in unstimulated condition (upper panel) and when polyclonally activated (lower panel). **(c)** Purity, activation markers and proliferation of CD4^+^ and CD8^+^ T cells polyclonally activated for 3 days before start of the migration assay. **(d)** Gating strategy of CTV+ T cells collected from the lower compartment in the ex-*vivo* migration assay. **(e)** Schematic overview of the co-culture system to evaluate T cell activation by polyclonal stimulation in presence of organoids. CTV-stained T cells were co-cultured with organoids previously treated with chemotherapeutics for 24 hrs and mechanically disrupted prior to co-culture start. 4 days after plating, T cells were analyzed by FACS for membrane activation marker expression and proliferation.**(f)** Representative EVOS images of organoids treated for 24h with Busulfan (35μM), Fludarabine (15μM), Clofarabine (0.5μM). Images are taken before the start of the co-culture assay, scale bar = 1000 um.

**Supplementary Figure 5.**
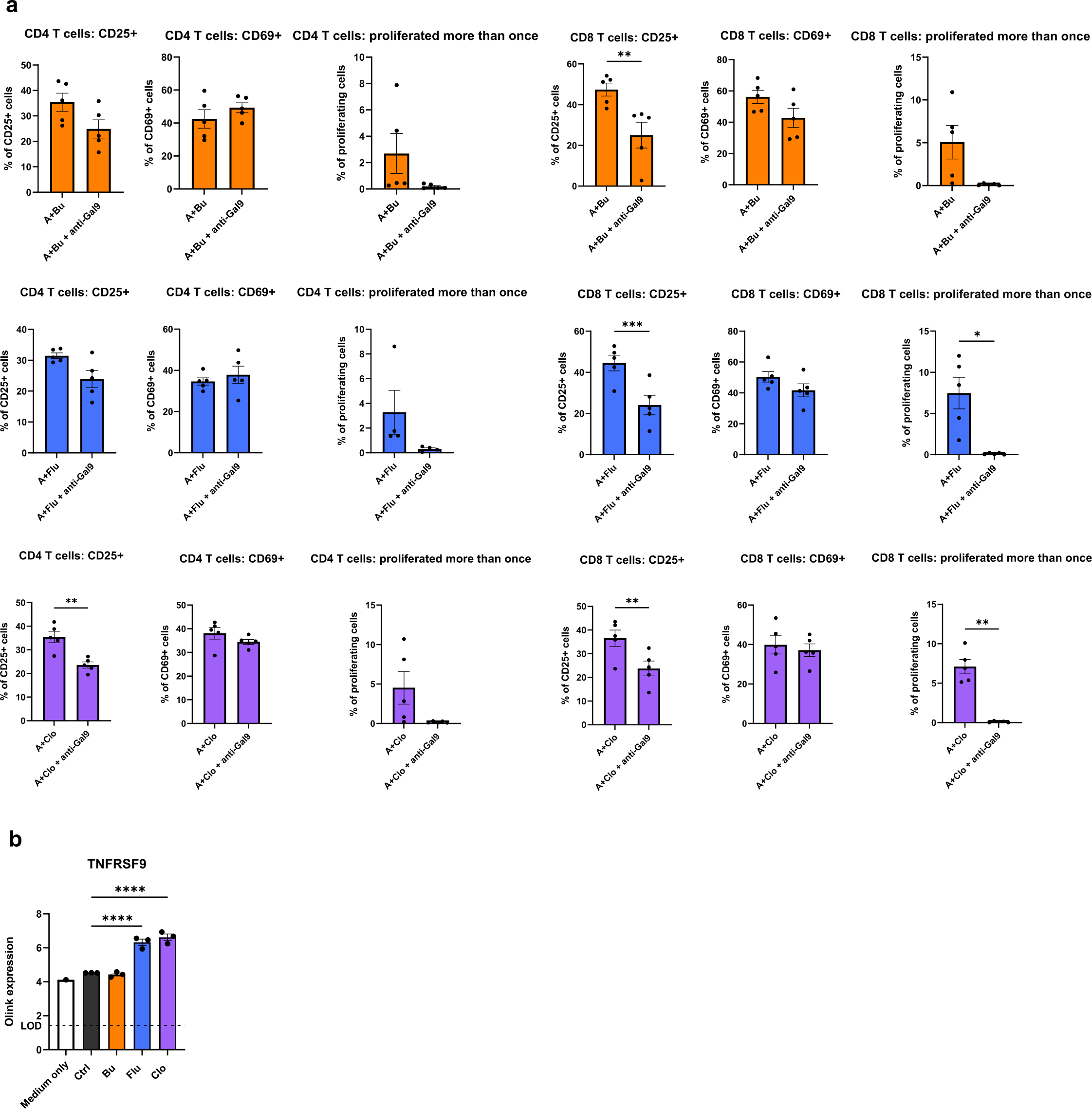
**(a)** CD4^+^ and CD8^+^ T cell activation after co-culture with treated organoids in the presence of anti- Gal-9 mAb. N≥5 T cell donors with 1 organoid donor, each data point indicates a T cell donor, mean with SEM, paired t-test. **(b)** Soluble-TNFRSF9 levels detected by Olink proteomics. Each data point indicates the levels of the detected protein in the CM from each organoid condition (log scale), mean with SEM, ANOVA.

**Supplementary Table 1.**
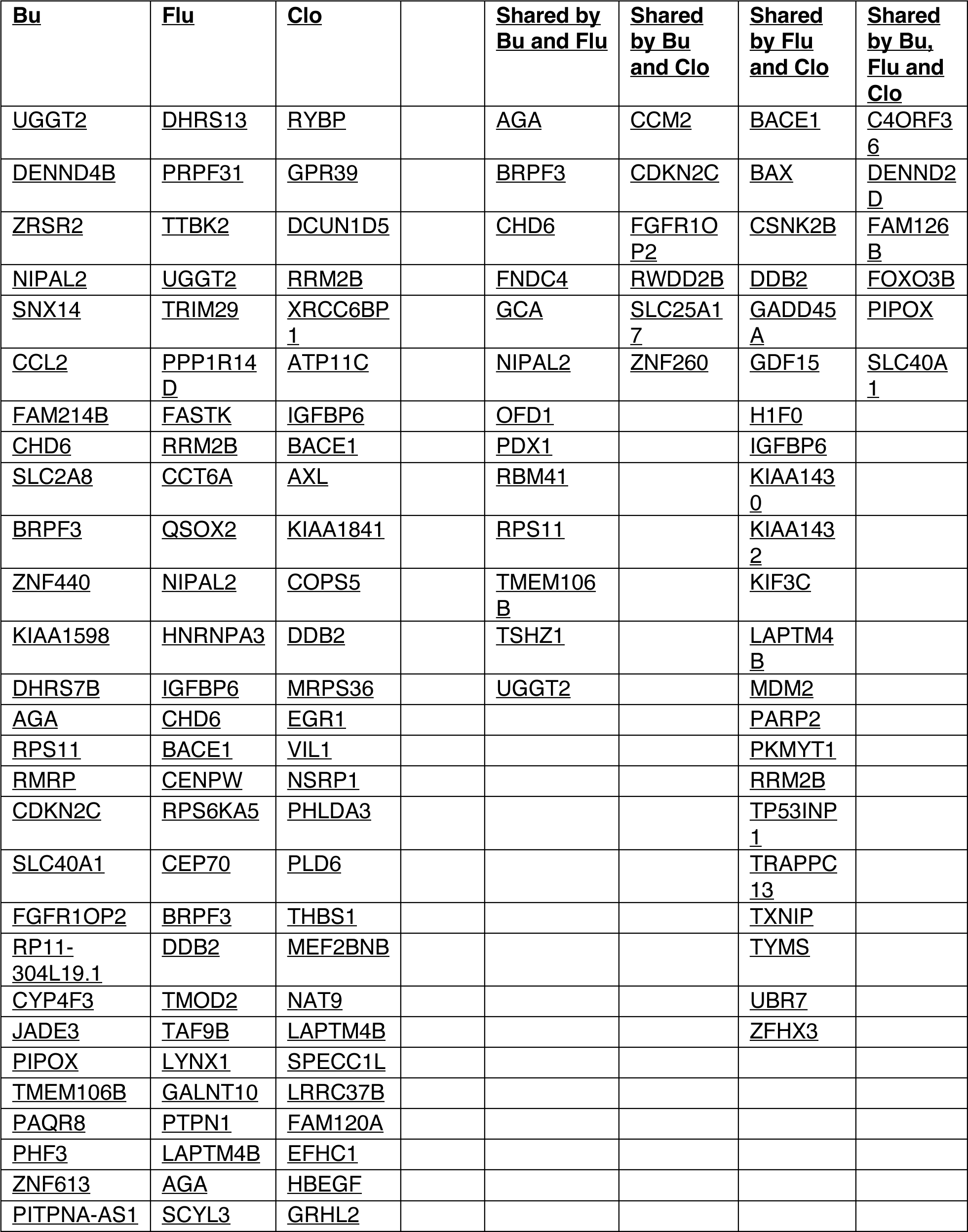
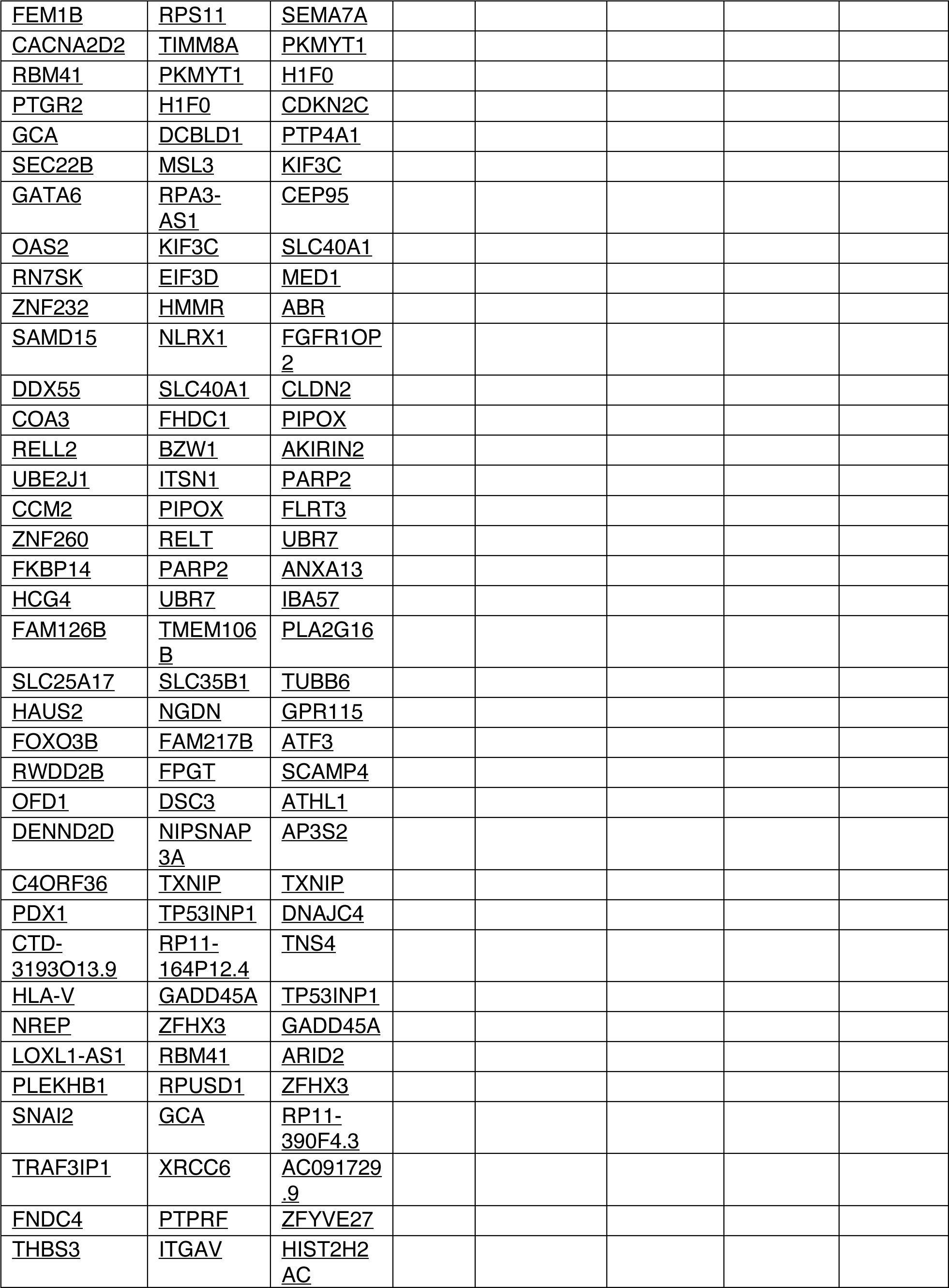
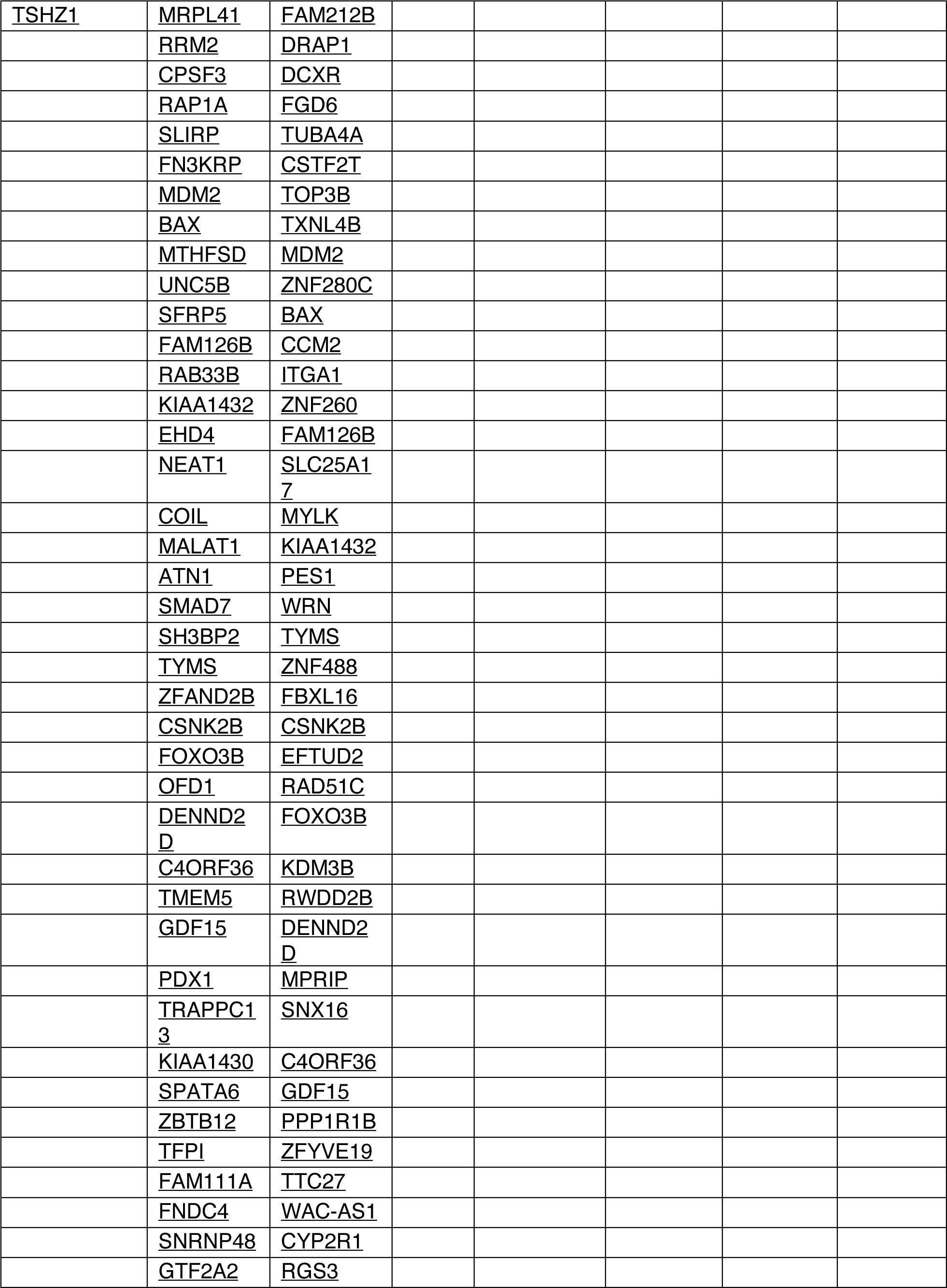
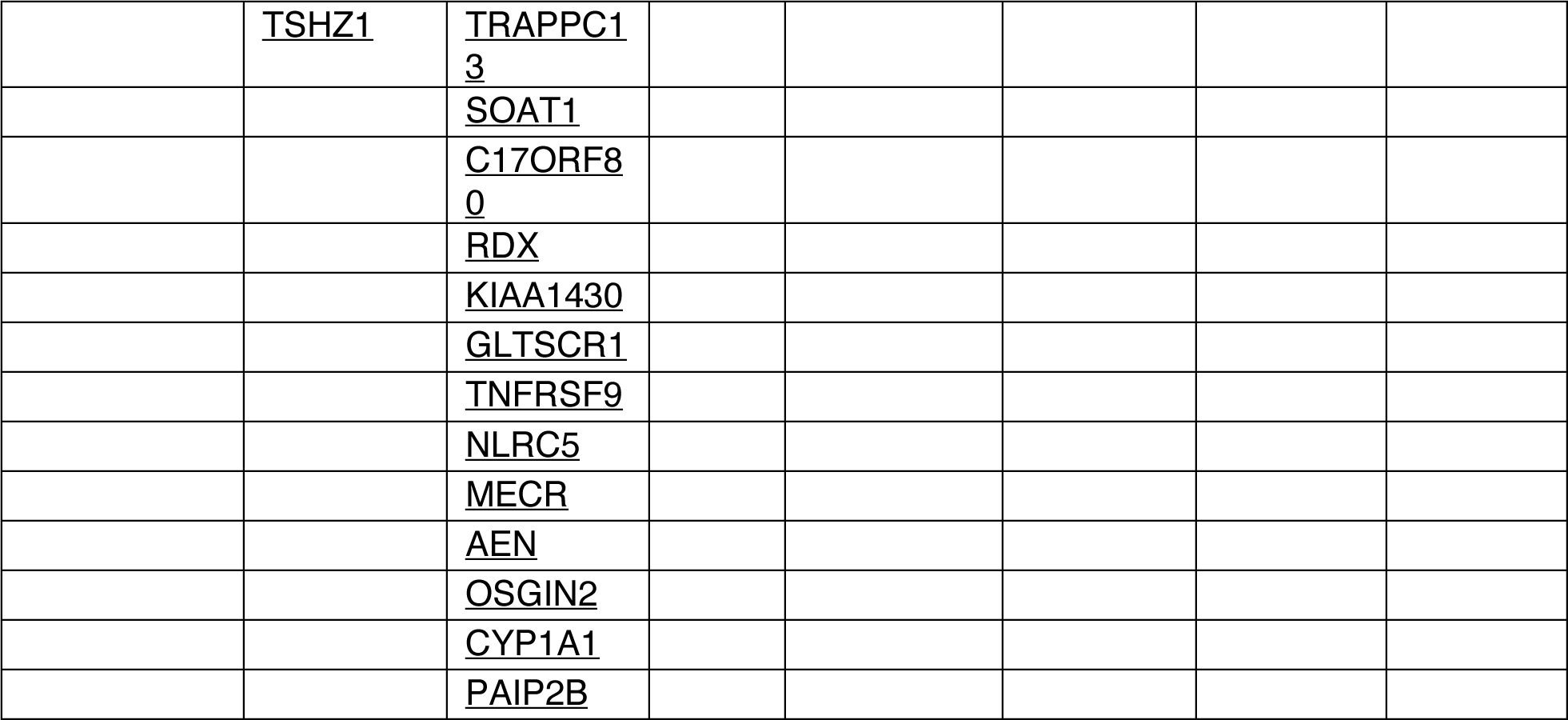
Differentially Expressed Genes

**Supplementary Table 2.**
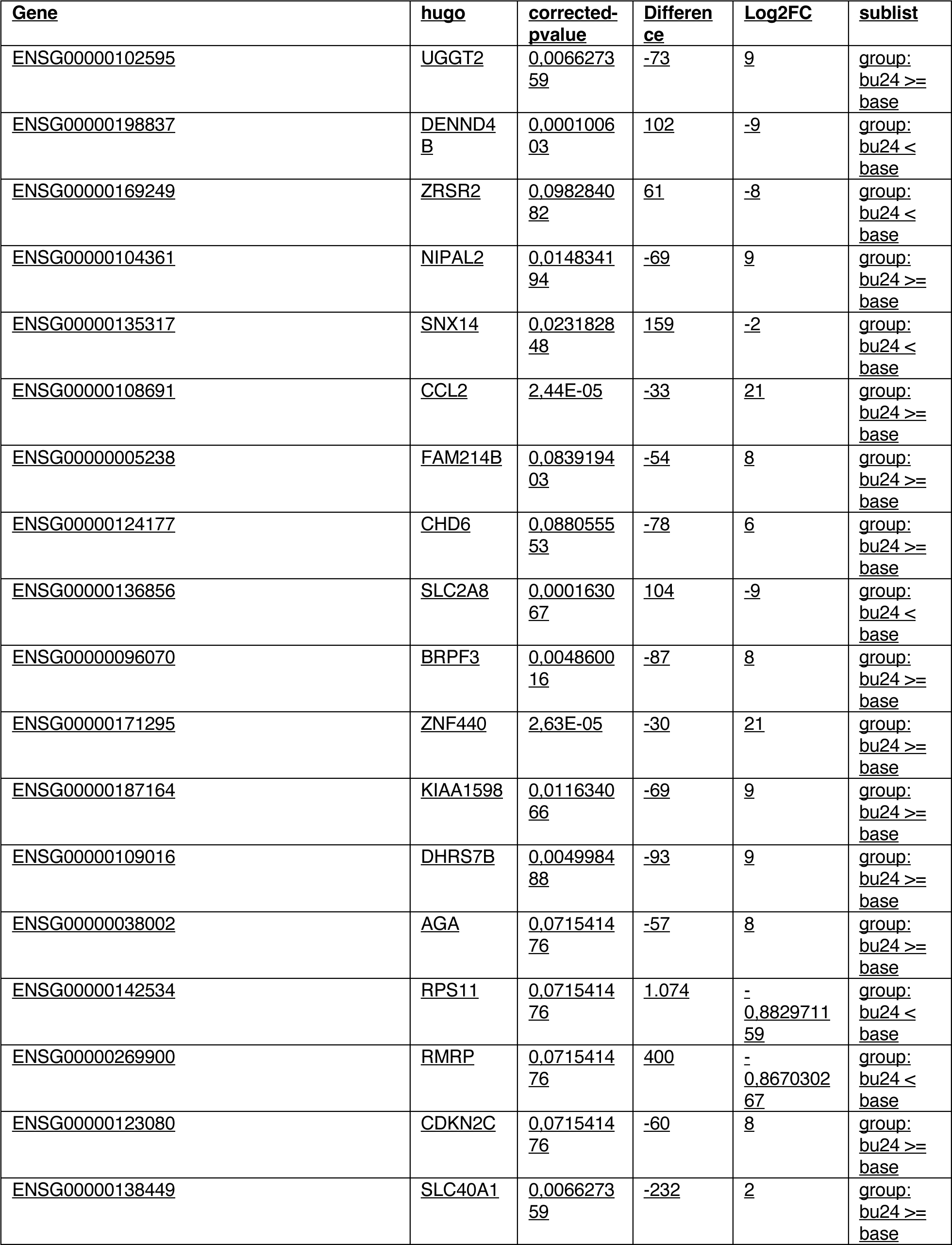
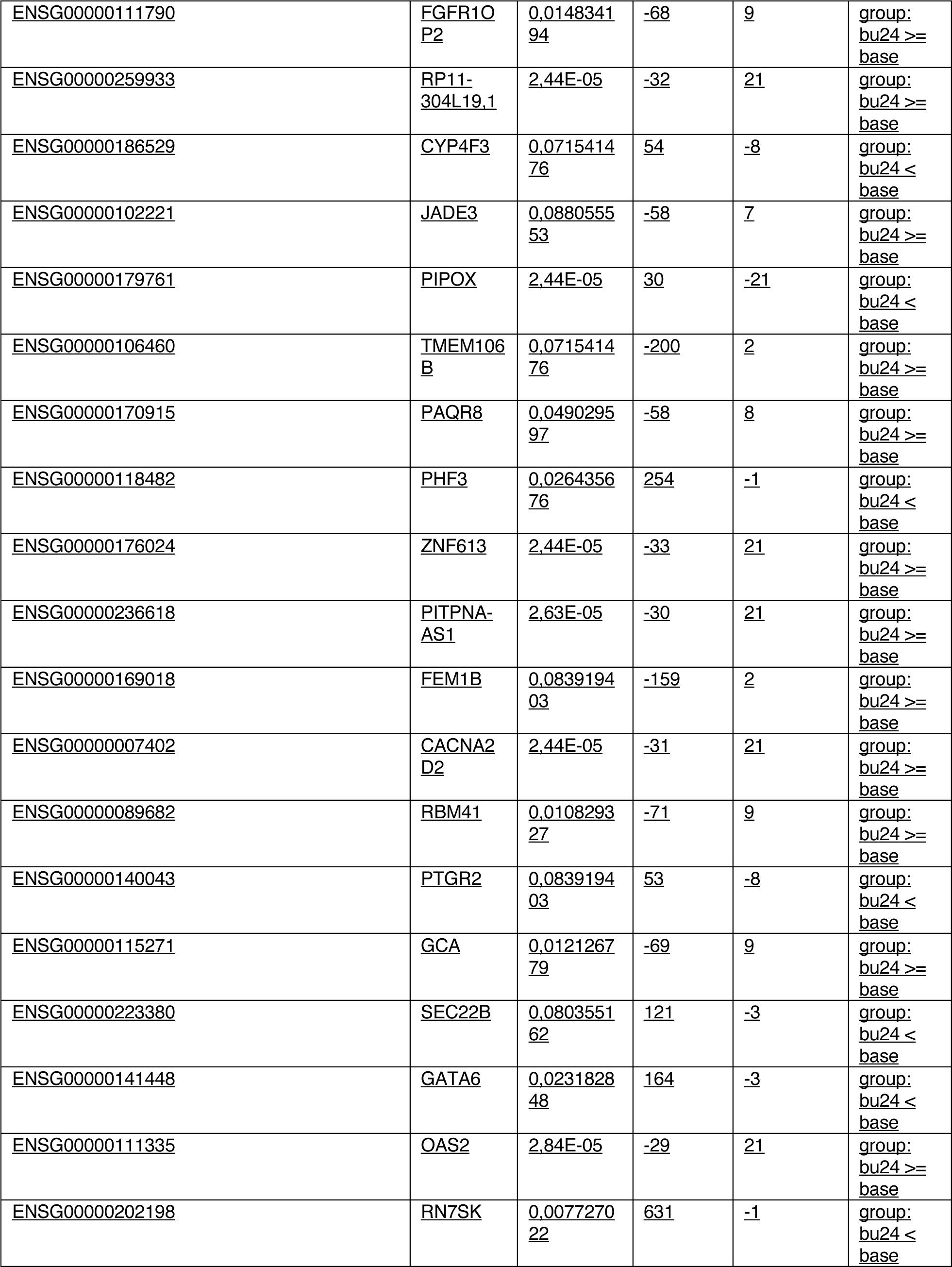
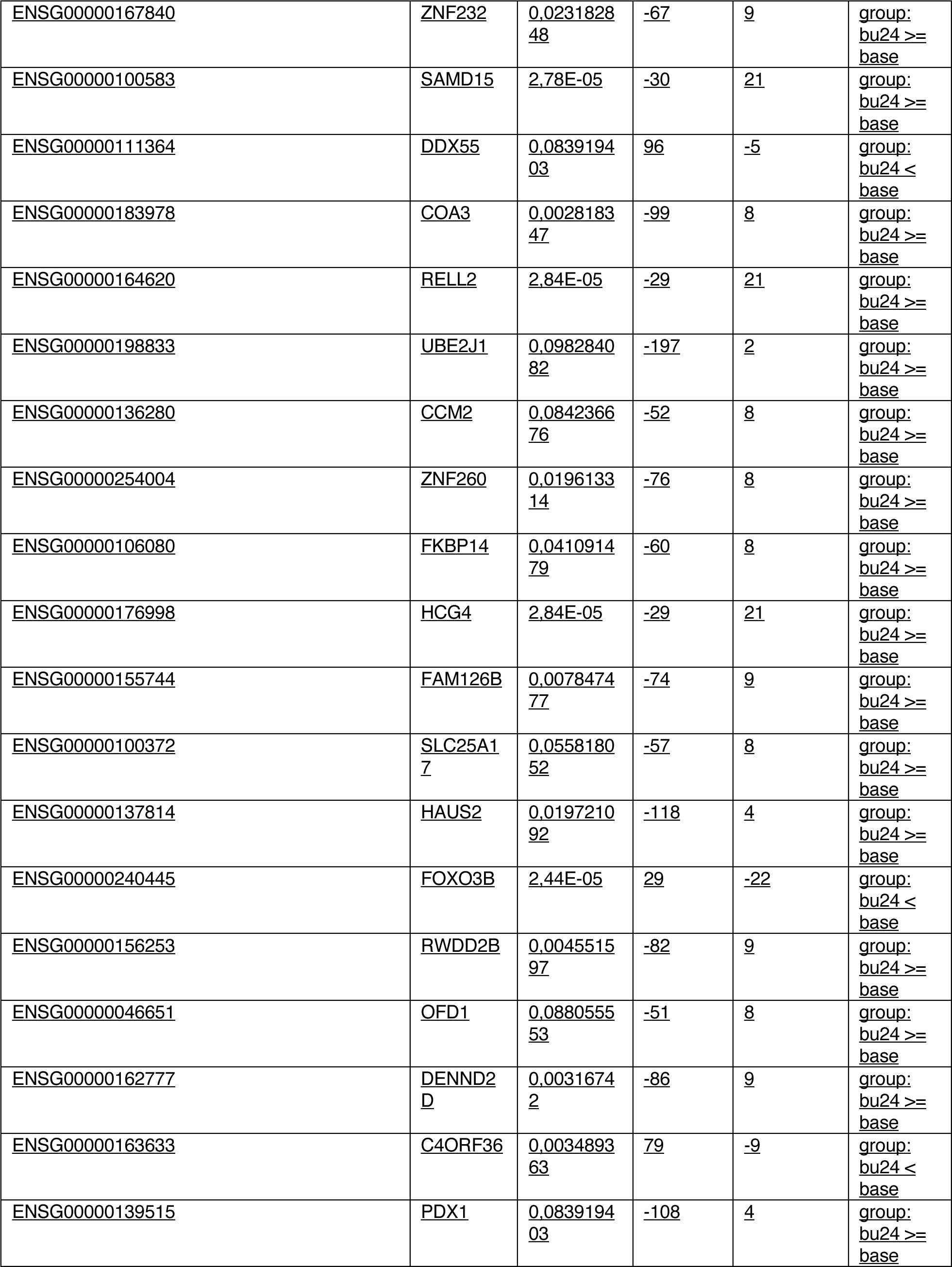
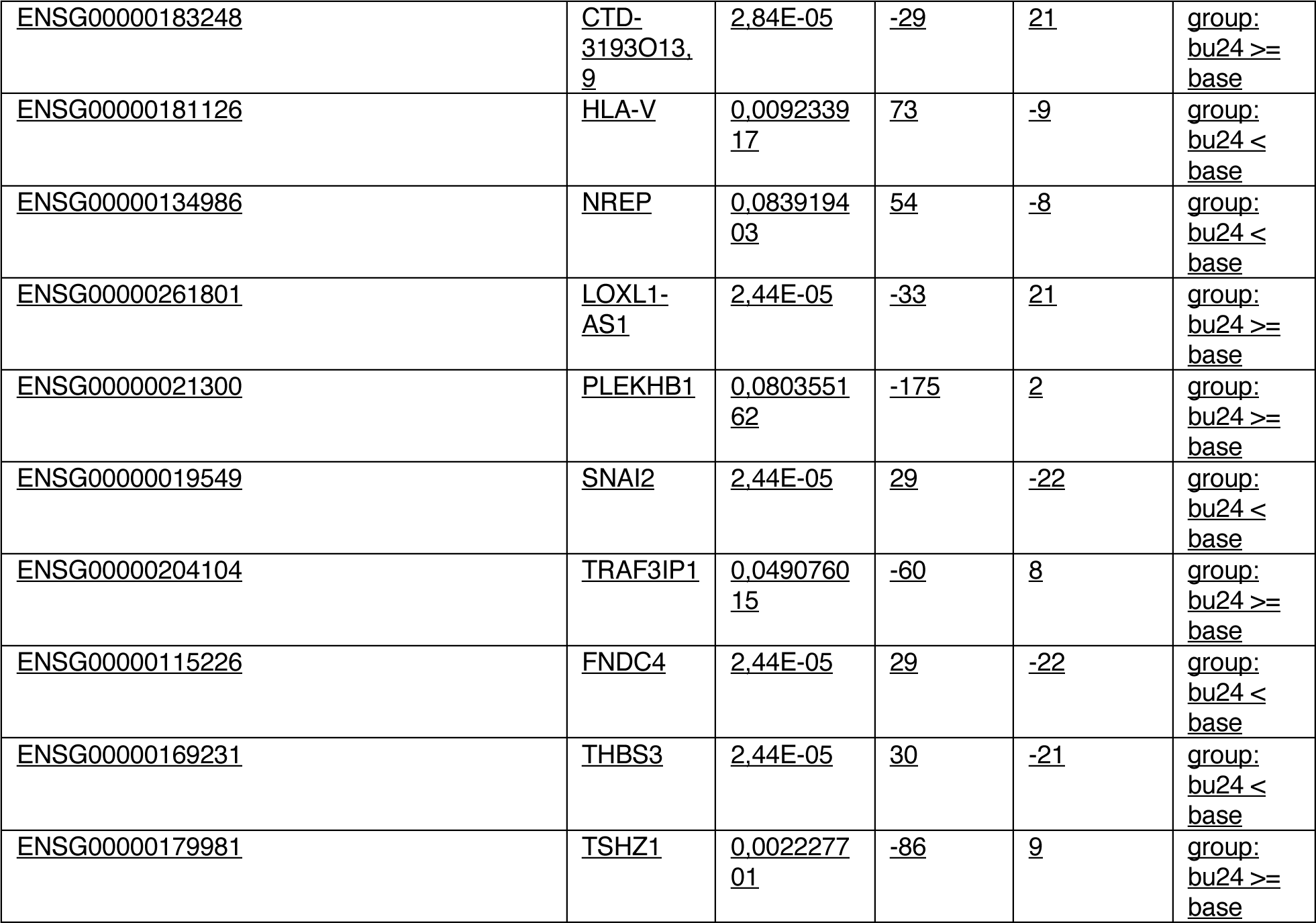
Differentially Expressed Genes after treatment with busulfan 24h

**Supplementary Table 3.**
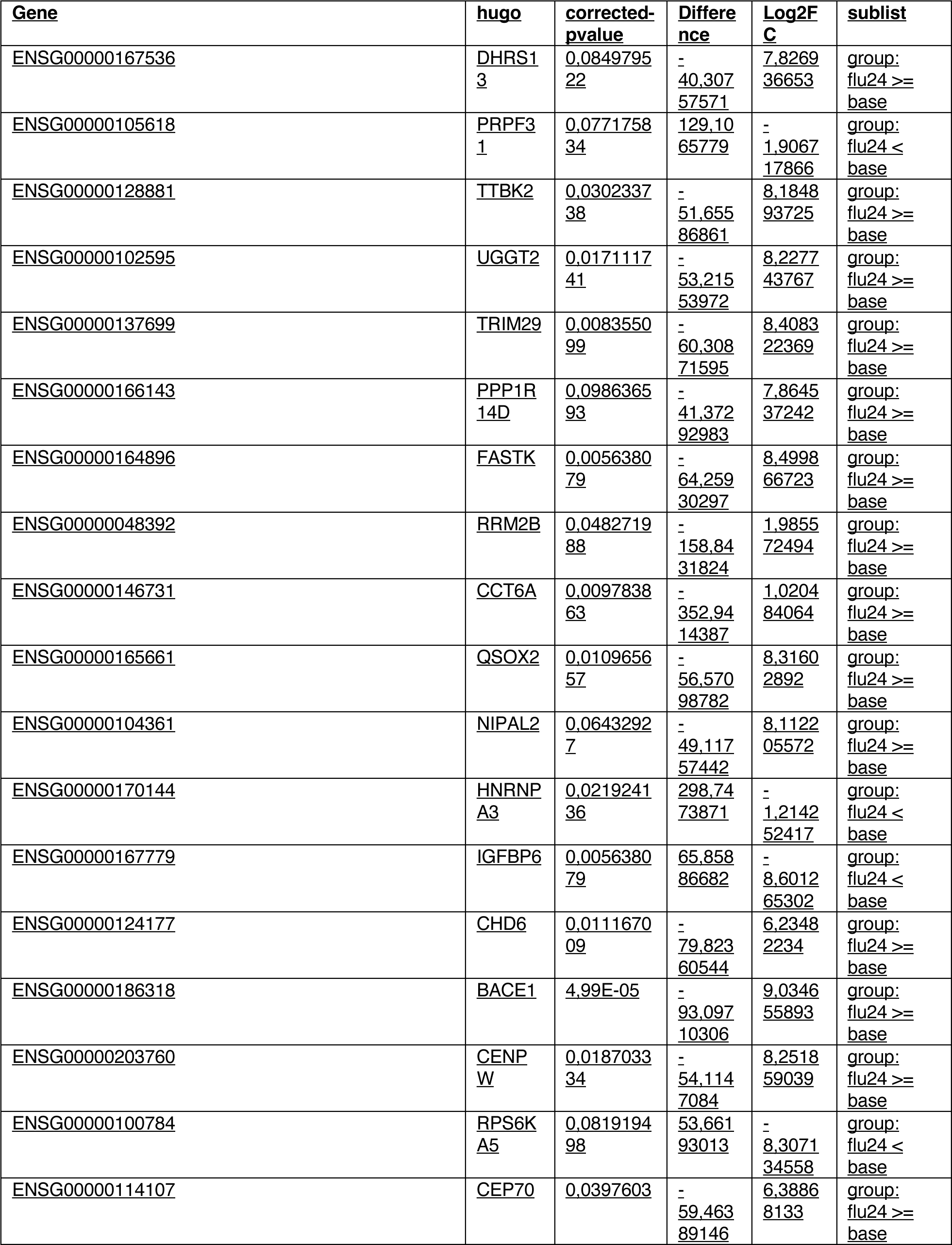
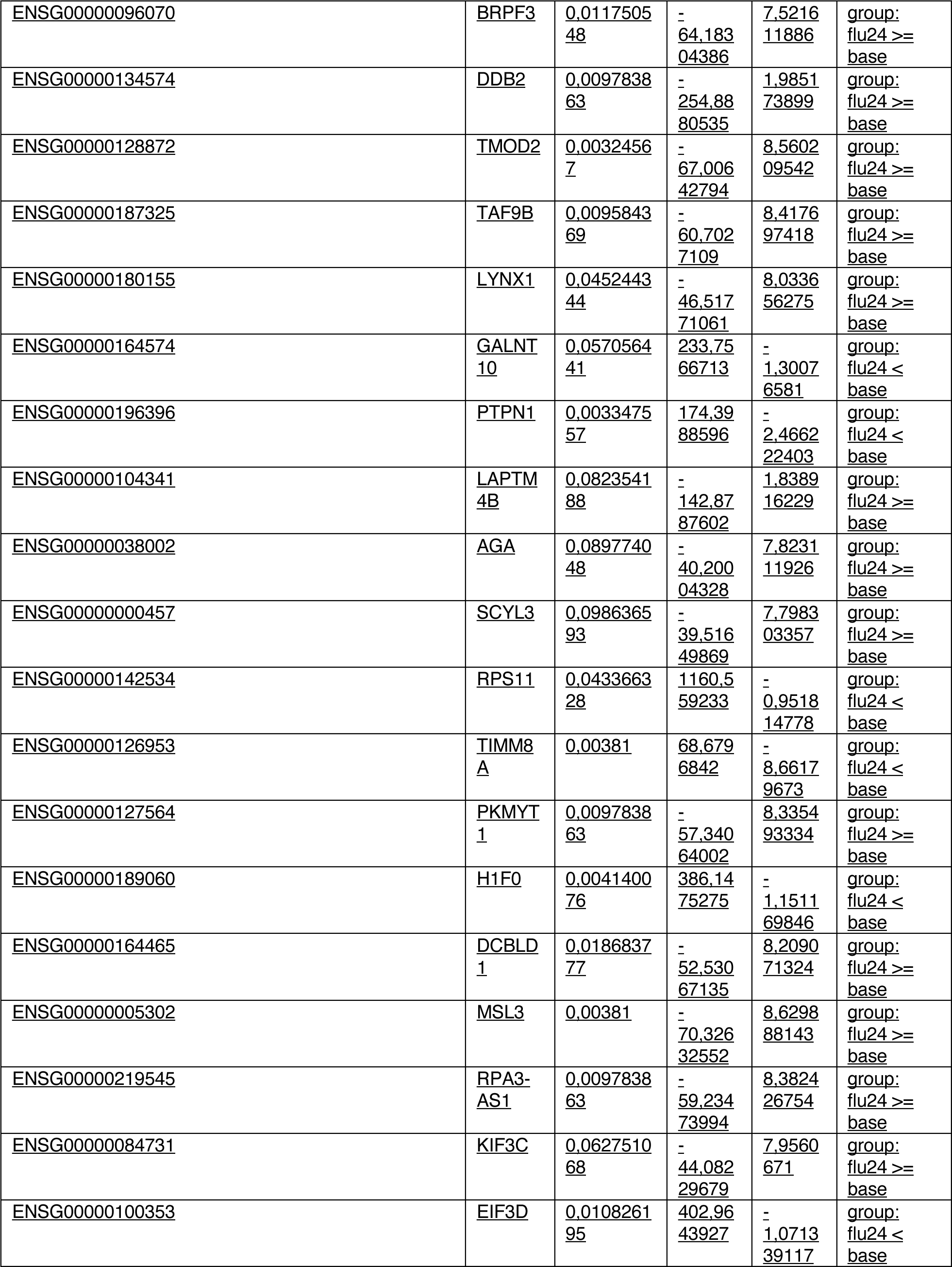
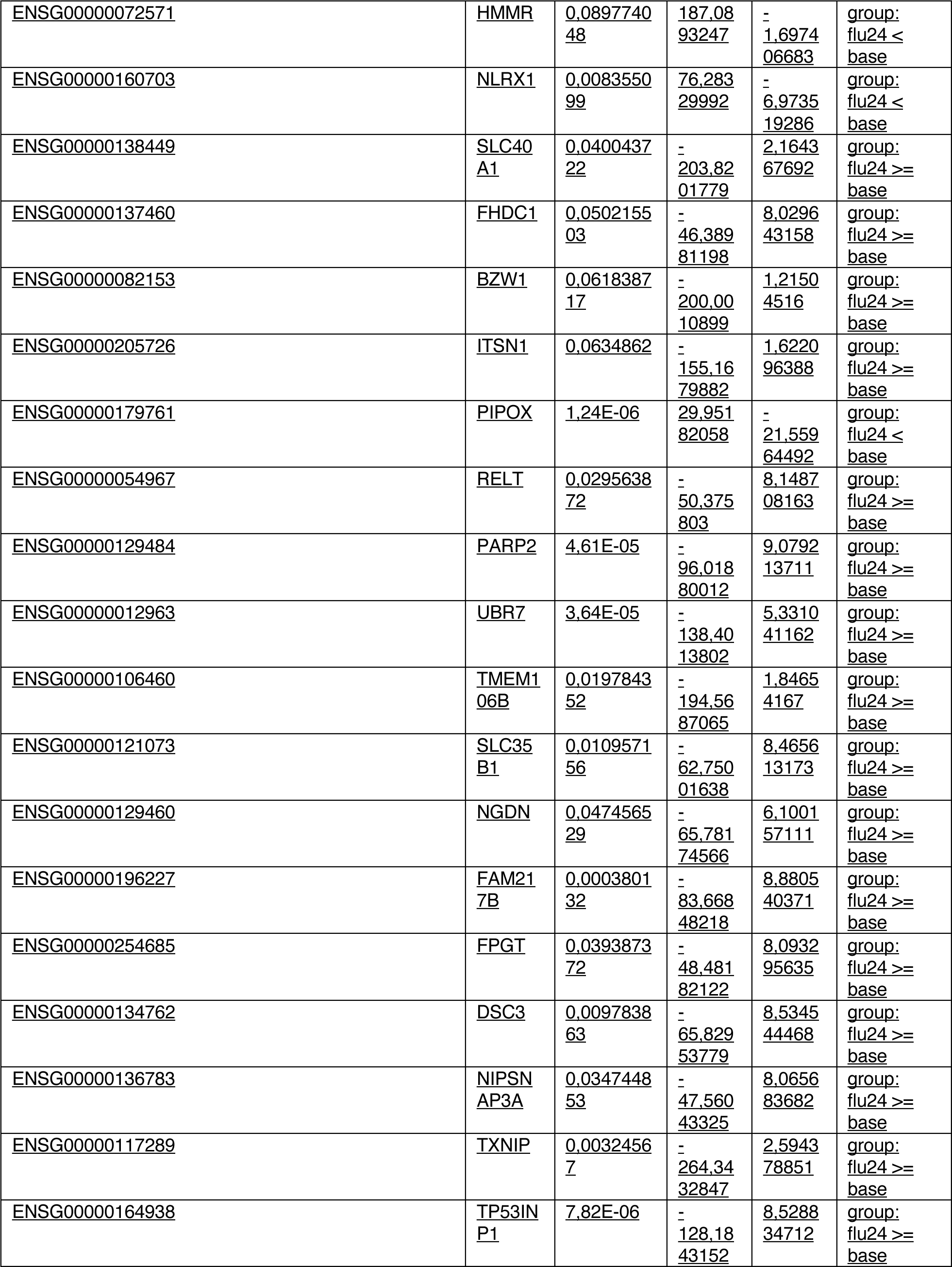
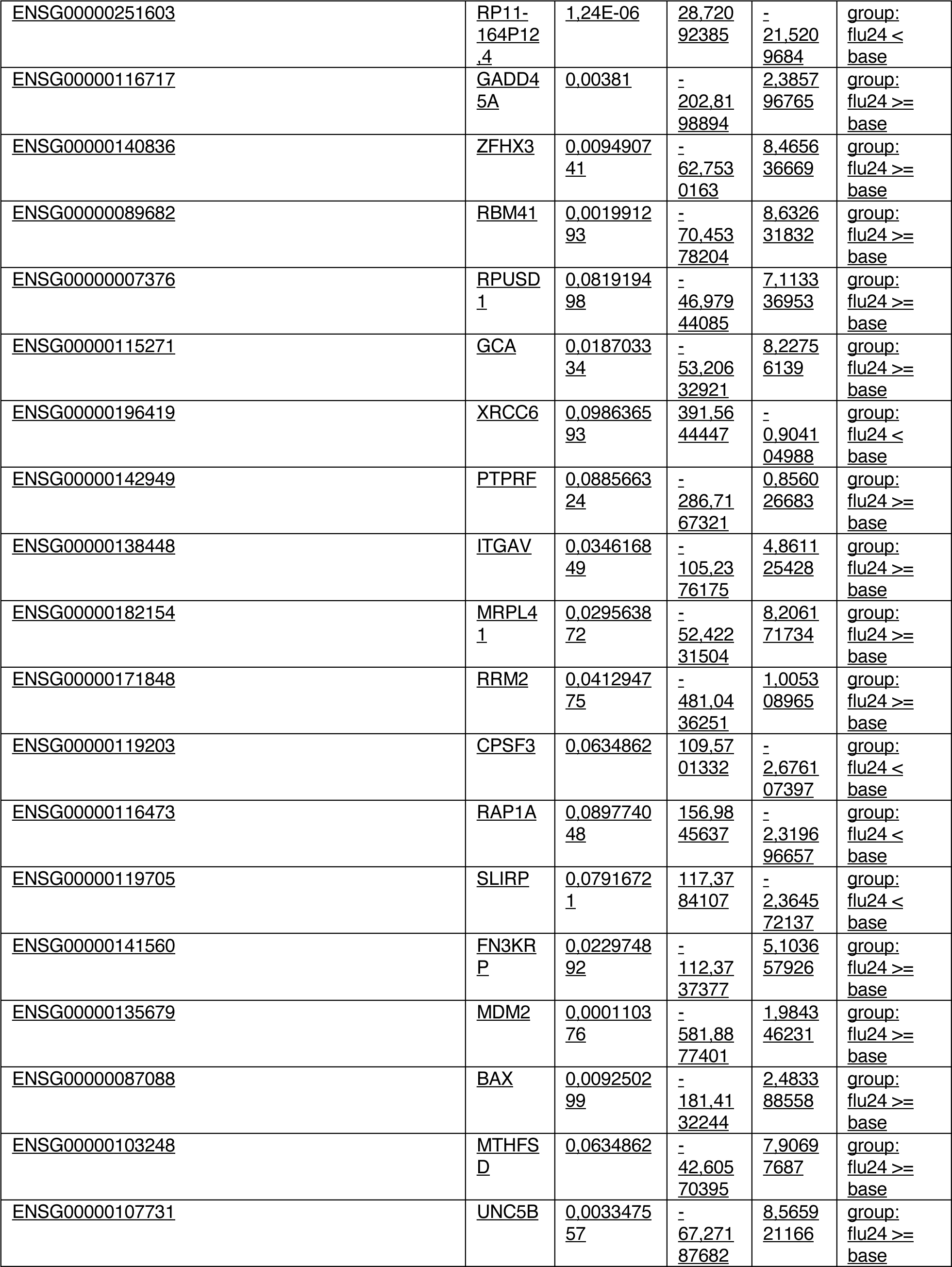
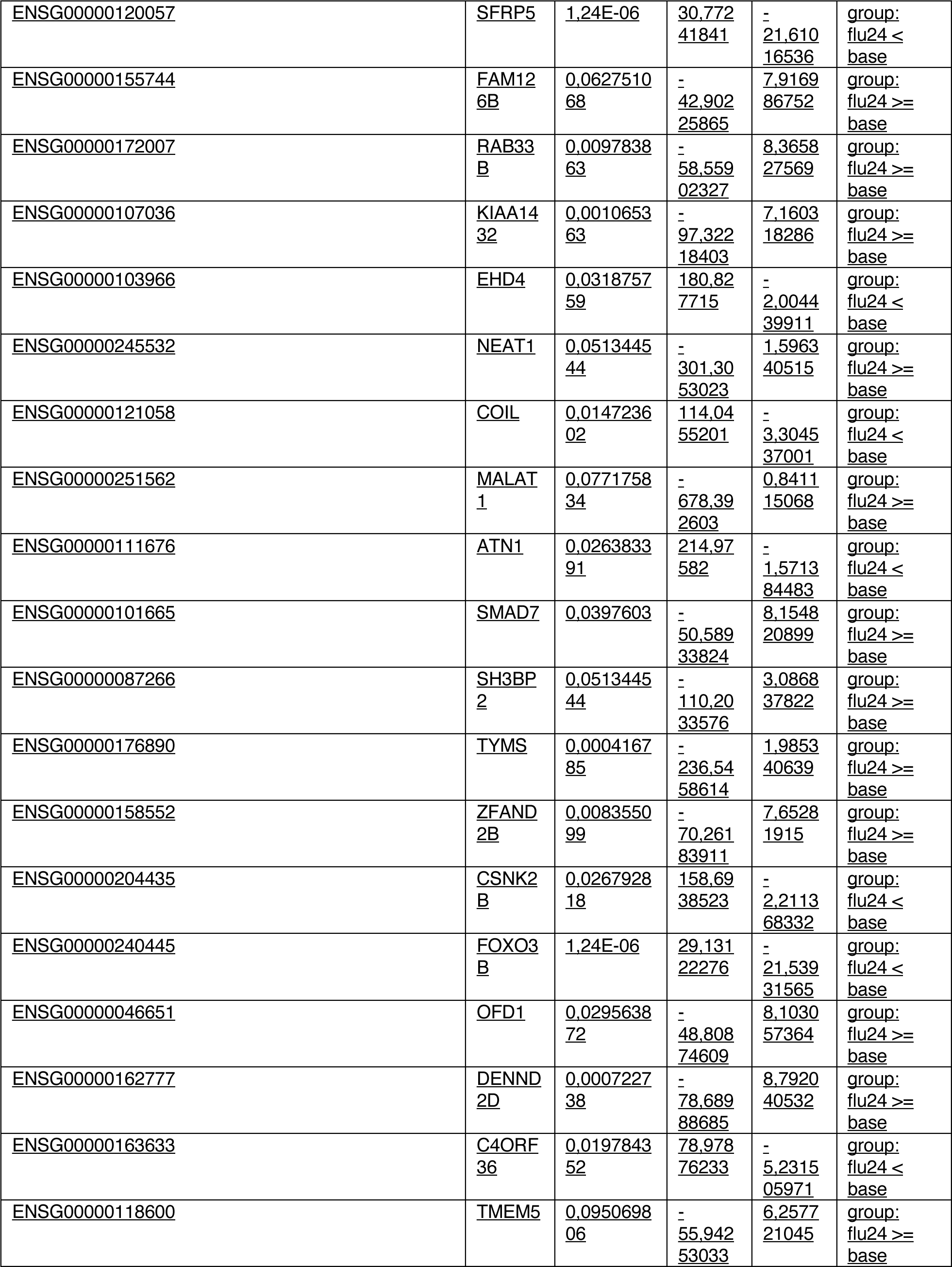
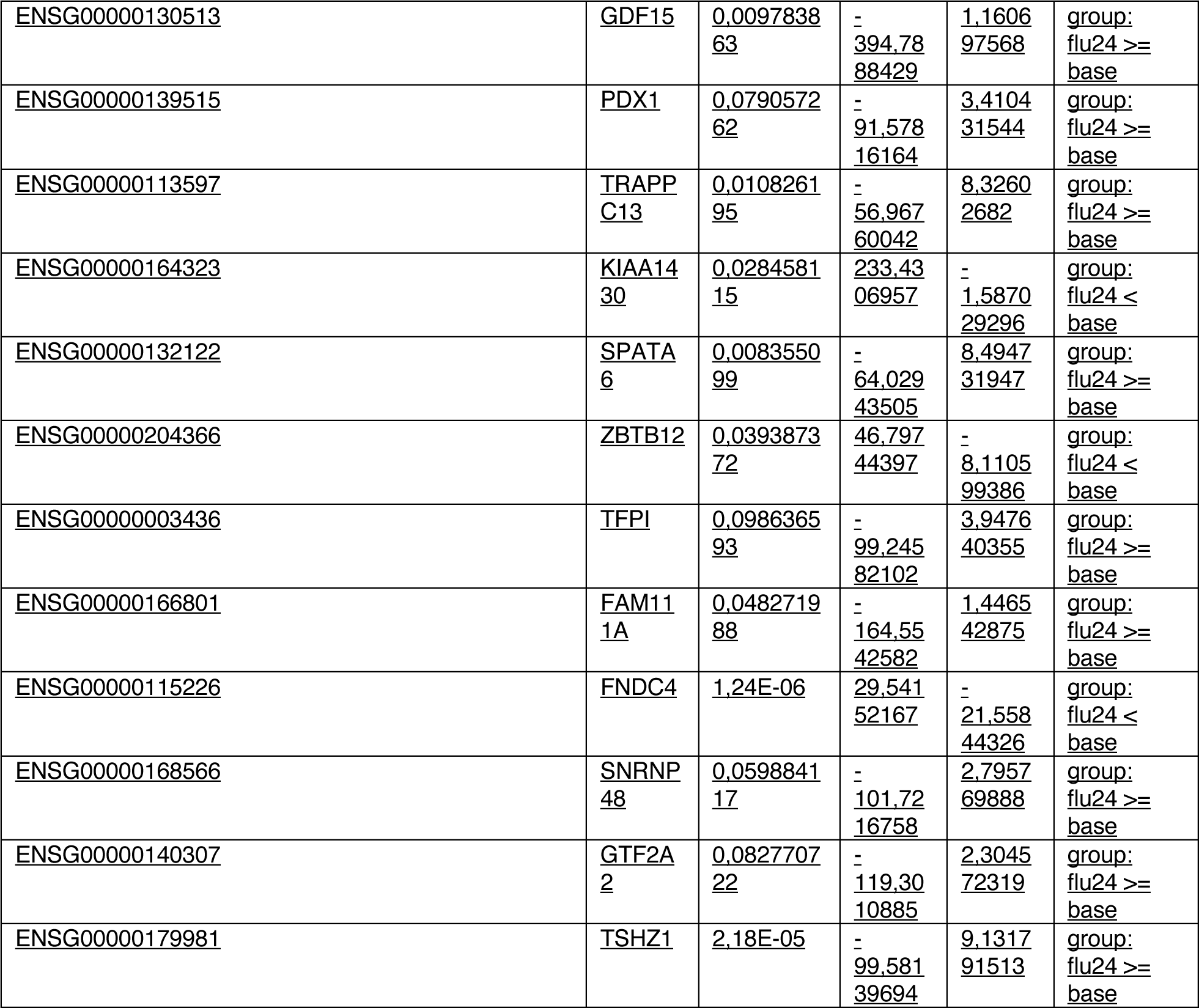
Differentially Expressed Genes after treatment with fludaribine 24h

**Supplementary Table 4.**
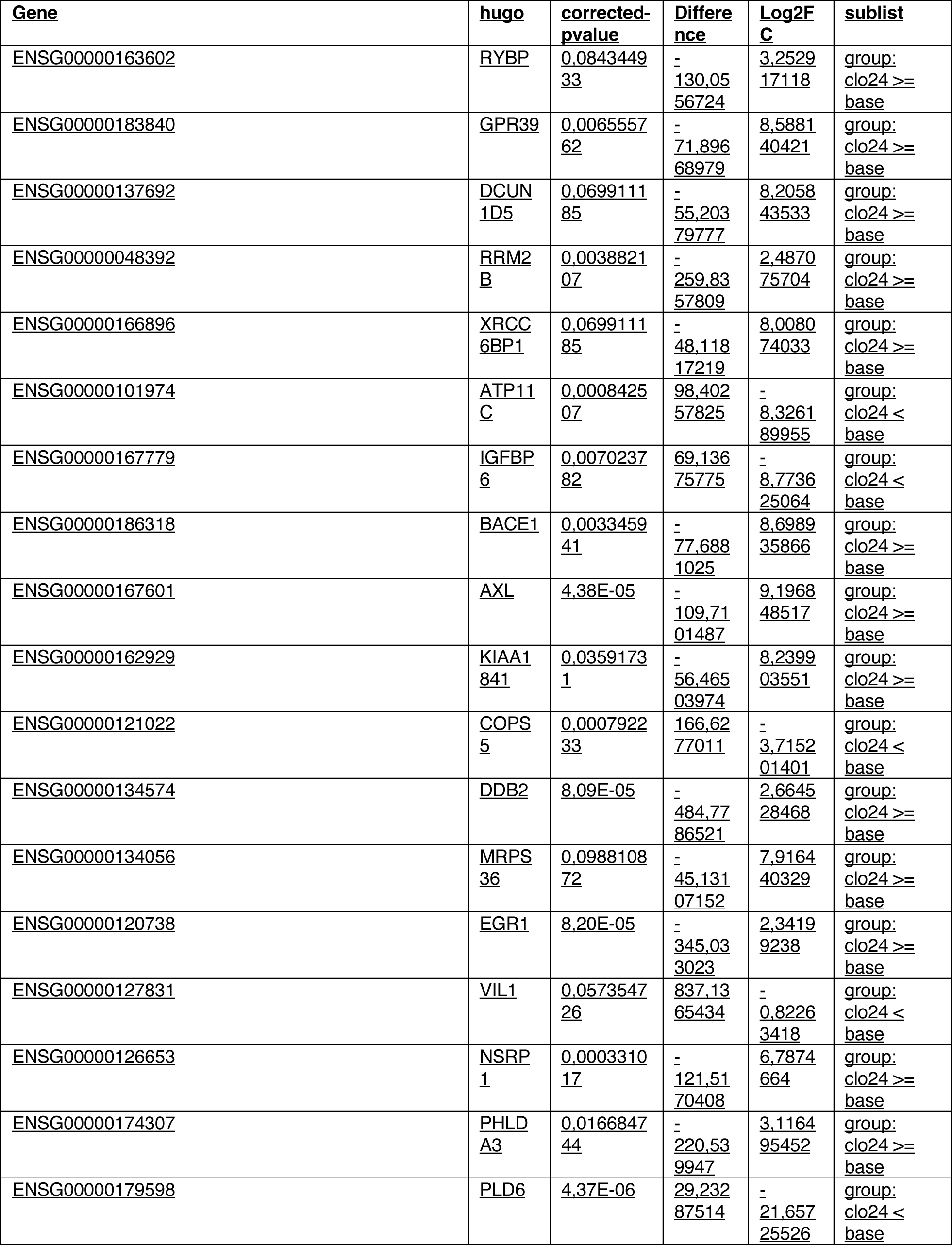
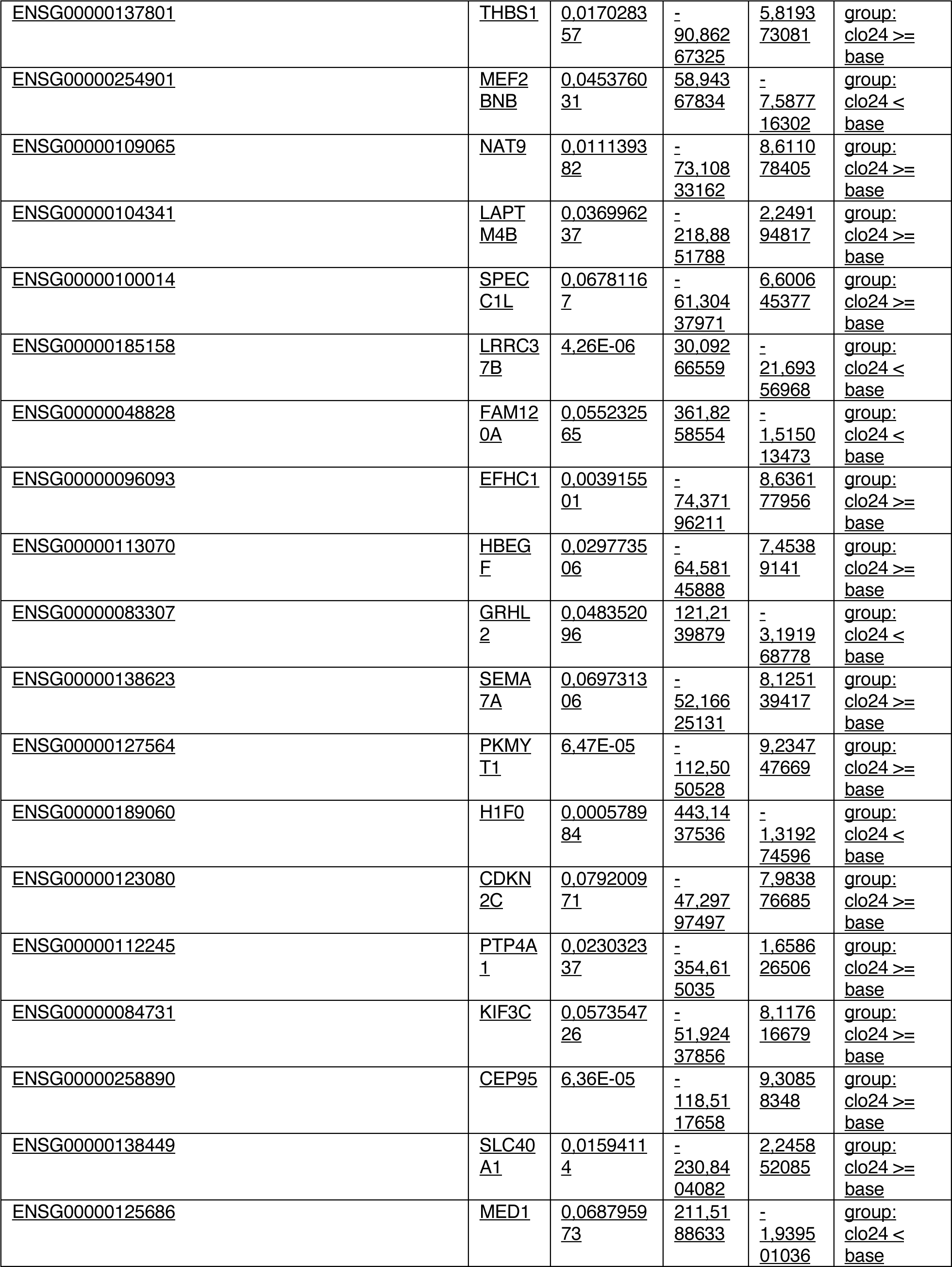
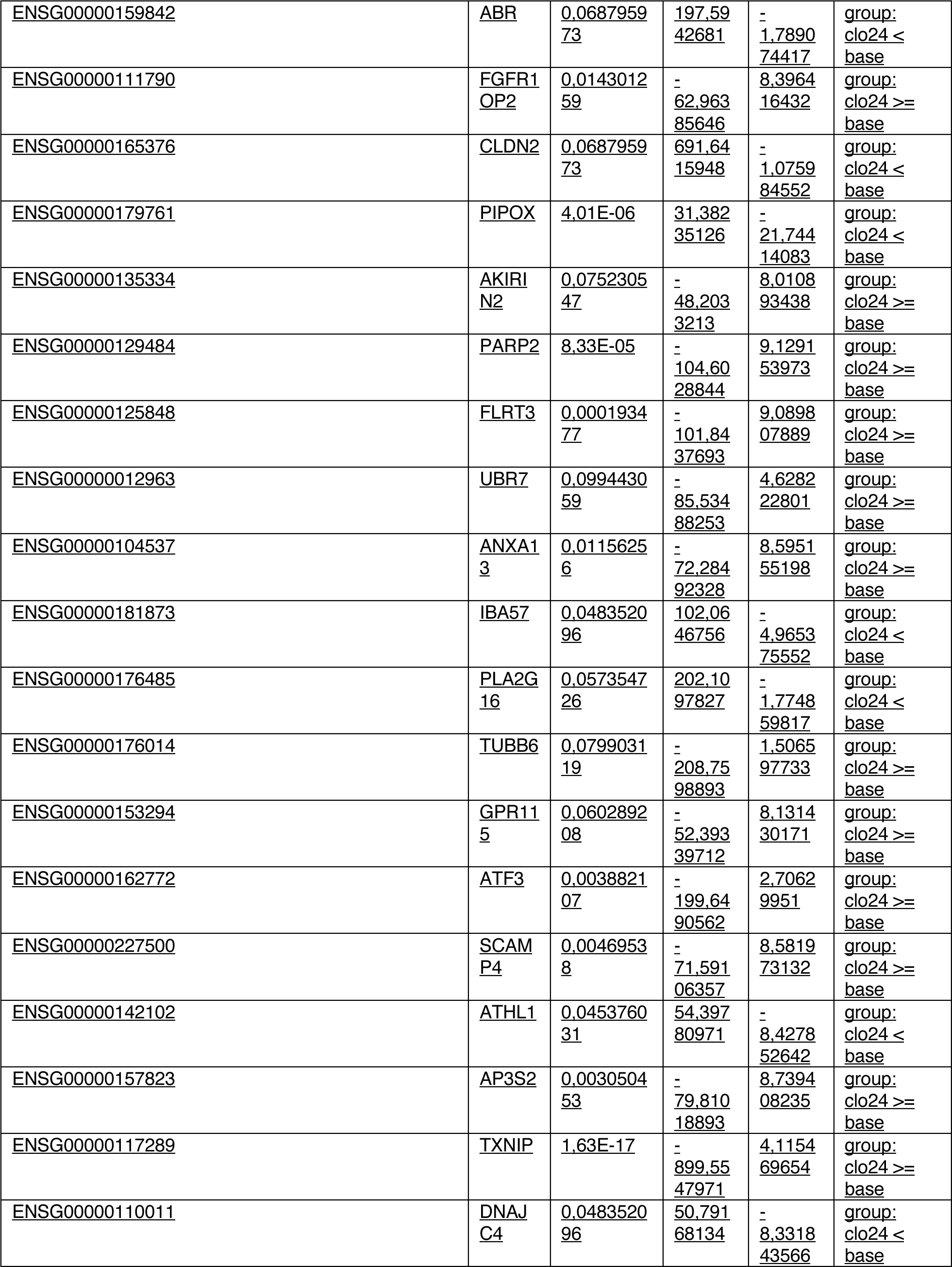
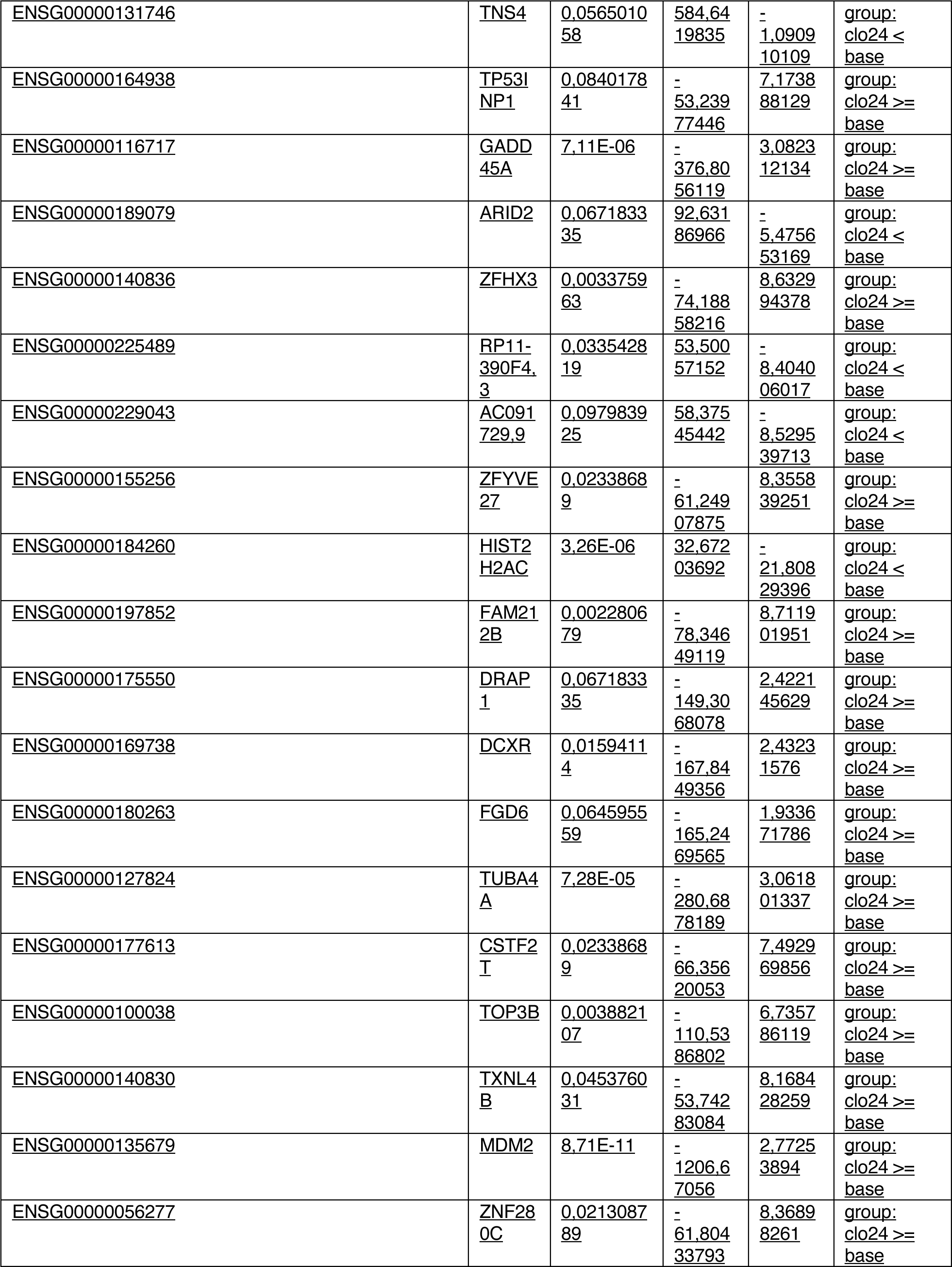
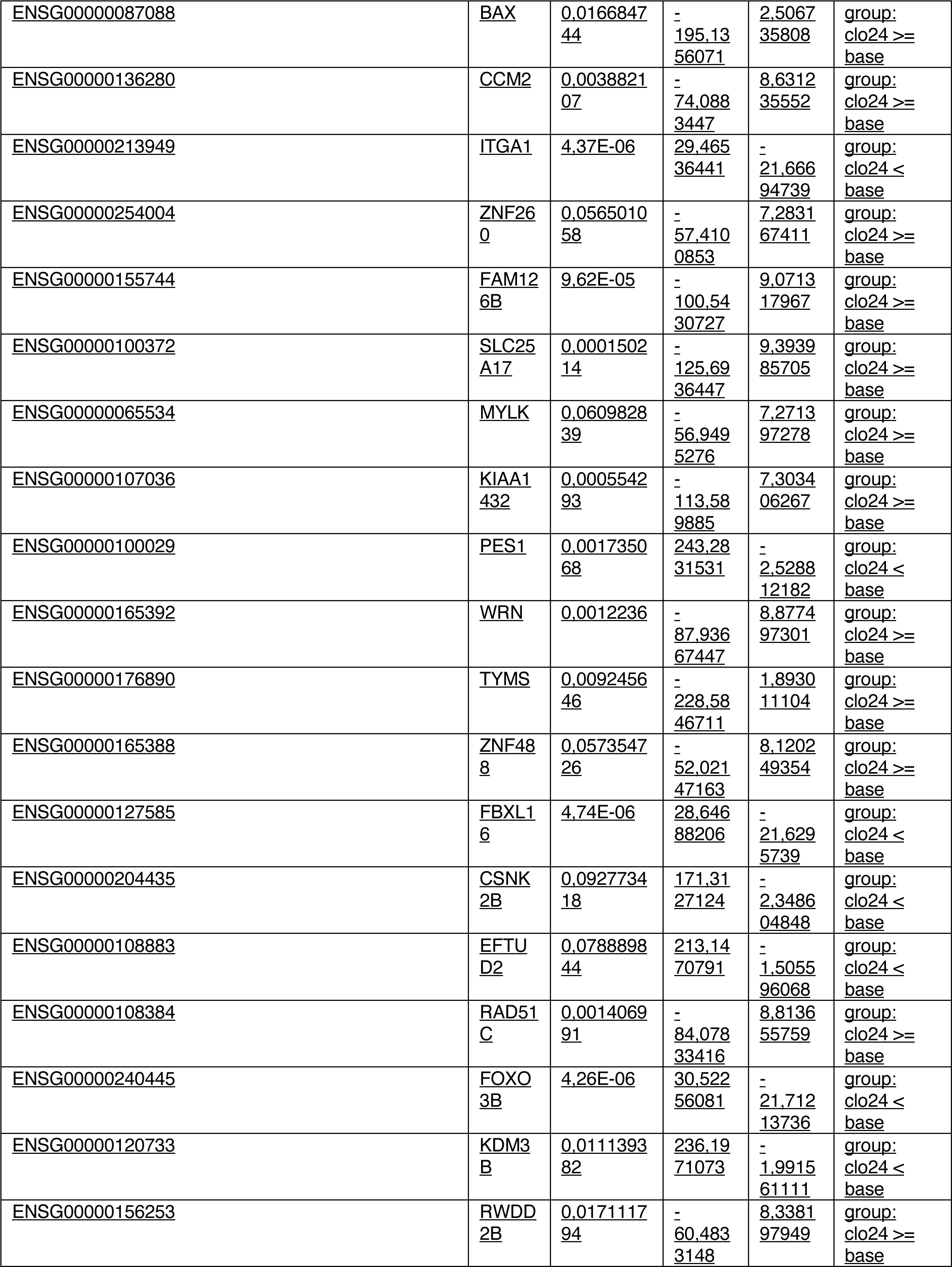
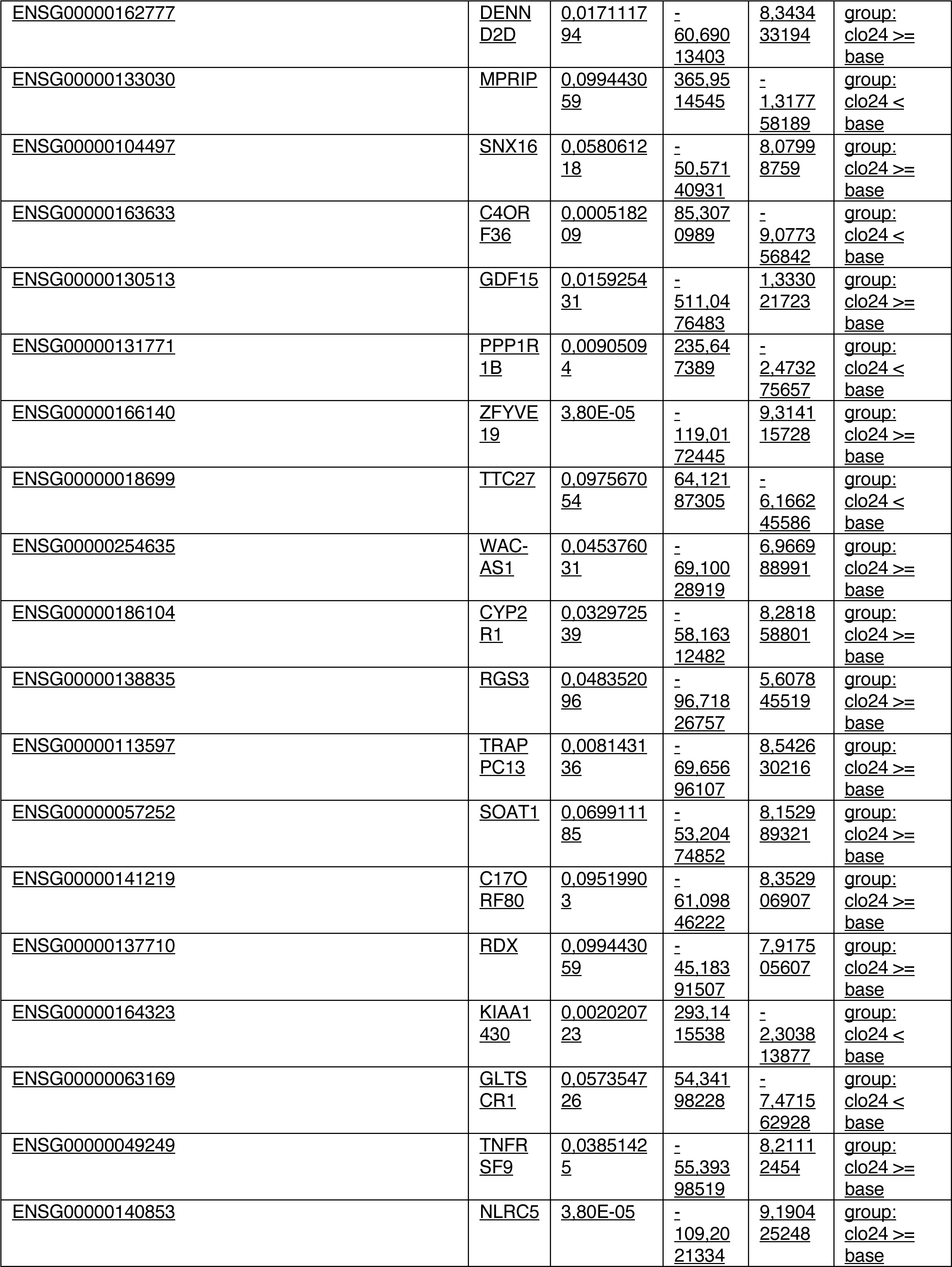
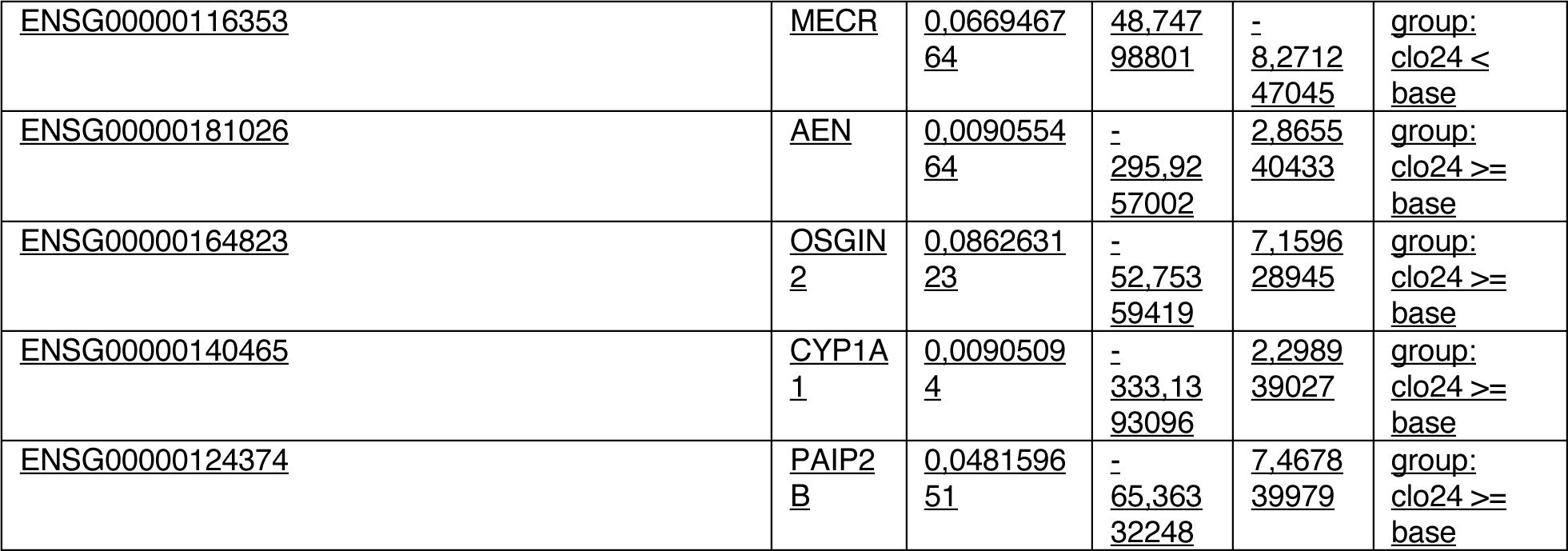
Differentially Expressed Genes after treatment with clofarabine 24h

**Supplementary Table 5.**
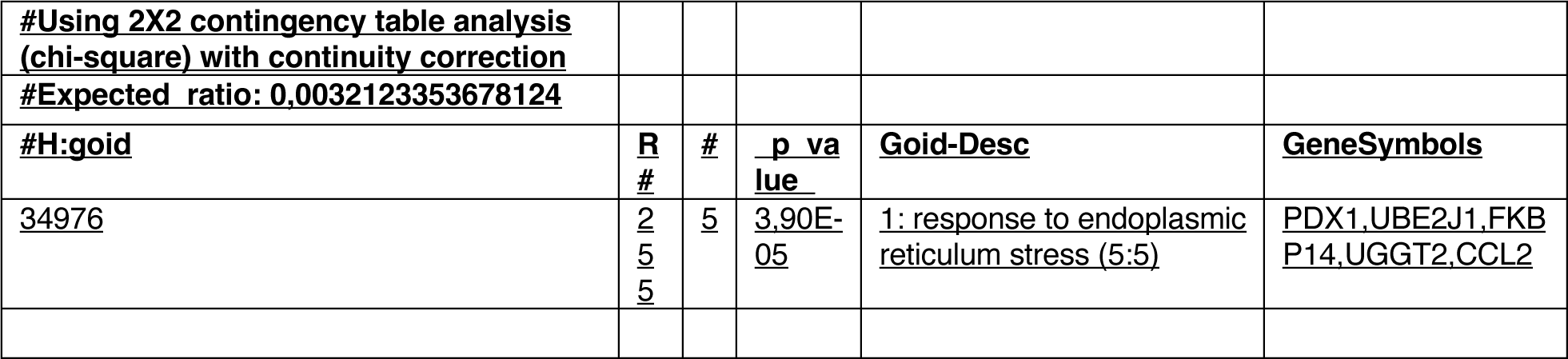
GO term analysis Bu≥Ctrl

**Supplementary Table 6.** GO term analysis Flu≥Ctrl

|  |  |  |  |  |  |
| --- | --- | --- | --- | --- | --- |
| #Using 2X2 contingency table analysis (chi-square) with continuity correction |  |  |  |  |  |
| #Expected ratio: 0.00594282043045294 |  |  |  |  |  |
| **#H:goid** | **R#** | **#** | **p value** | **Goid-Desc** | **GeneSymbols** |
| 48523 | 3214 | 27 | 0.05 | 1: negative regulation of cellular process (3:5) | TSHZ1, TXNIP, MSL3, TTBK2, GADD45A, UNC5B, TRIM29, TMOD2, SLC40A1, PDX1, ITGAV, SMAD7, MDM2, ZFH3, RRM2B, TAF9B, PPP1R14D, LAPTM4B, RIC1, BAX, RRM2, ITS1, TFPI, TYMS, RAB33B, PKMYT1, TP53INP1 |
| 12501 | 1489 | 18 | 1.90E-03 | 1: programmed cell death (4:4) | PARP2, TXNIP, FASTK, GADD45A, DSC3, UNC5B, SLC40A1, PDX1, ITGAV, MDM2, RRM2B, TAF9B, BAX, ITS1, MRPL41, RELT, TP53INP1, GDF15 |
| 31324 | 1811 | 18 | 0.03 | 1: negative regulation of cellular metabolic process (4:6) | TSHZ1, TXNIP, MSL3, GADD45A, TRIM29, PDX1, ITGAV, SMAD7, MDM2, ZFH3, TAF9B, PPP1R14D, LAPTM4B, RIC1, BAX, TFPI, TYMS, PKMYT1 |
| 8219 | 1581 | 18 | 4.50E-03 | 1: cell death (3:3) | PARP2, TXNIP, FASTK, GADD45A, DSC3, UNC5B, SLC40A1, PDX1, ITGAV, MDM2, RRM2B, TAF9B, BAX, ITS1, MRPL41, RELT, TP53INP1, GDF15 |
| 51172 | 1692 | 17 | 0.03 | 1: negative regulation of nitrogen compound metabolic process (4:6) | TSHZ1, TXNIP, MSL3, GADD45A, TRIM29, PDX1, ITGAV, SMAD7, MDM2, ZFH3, TAF9B, PPP1R14D, LAPTM4B, RIC1, BAX, TFPI, TYMS |
| 6915 | 1426 | 17 | 3.30E-03 | 1: apoptotic process (5:5) | PARP2, TXNIP, FASTK, GADD45A, UNC5B, SLC40A1, PDX1, ITGAV, MDM2, RRM2B, TAF9B, BAX, ITS1, MRPL41, RELT, TP53INP1, GDF15 |
| 10941 | 1214 | 14 | 0.01 | 1: regulation of cell death (4:5) | TXNIP, GADD45A, UNC5B, SLC40A1, PDX1, ITGAV, MDM2, RRM2B, TAF9B, BAX, ITS1, RELT, TP53INP1, GDF15 |
| 42981 | 1116 | 14 | 5.10E-03 | 1: regulation of apoptotic process (6:7) | TXNIP, GADD45A, UNC5B, SLC40A1, PDX1, ITGAV, MDM2, RRM2B, TAF9B, BAX, ITS1, RELT, TP53INP1, GDF15 |
| 43067 | 1130 | 14 | 5.90E-03 | 1: regulation of programmed cell death (5:6) | TXNIP, GADD45A, UNC5B, SLC40A1, PDX1, ITGAV, MDM2, RRM2B, TAF9B, BAX, ITS1, RELT, TP53INP1, GDF15 |
| 44092 | 759 | 11 | 3.50E-03 | 1: negative regulation of molecular function (4:4) | TXNIP, TTBK2, GADD45A, SMAD7, MDM2, PPP1R14D, PTPRF, BAX, LYNX1, TFPI, PKMYT1 |
| 97190 | 483 | 11 | 4.10E-06 | 1: apoptotic signaling pathway (4:6) | PARP2, FASTK, UNC5B, PDX1, ITGAV, MDM2, RRM2B, TAF9B, BAX, RELT, TP53INP1 |
| 51248 | 791 | 10 | 0.02 | 1: negative regulation of <u>protein metabolic process (5:7)</u> | GADD45A, ITGAV, SMAD7, MDM2, PPP1R14D, LAPTM4B, RIC1, BAX, TFPI, TYMS |
| <u>32269</u> | <u>756</u> | <u>9</u> | <u>0.05</u> | <u>1: negative regulation of cellular protein metabolic process (5:8)</u> | <u>GADD45A, SMAD7, MDM2, PPP1R14D, LAPTM4B, RIC1, BAX, TFPI, TYMS</u> |
| <u>43066</u> | <u>617</u> | <u>9</u> | <u>9.40E-03</u> | <u>1: negative regulation of apoptotic process (6:8)</u> | <u>UNC5B, SLC40A1, PDX1, ITGAV, MDM2, RRM2B, TAF9B, BAX, ITSN1</u> |
| <u>43069</u> | <u>631</u> | <u>9</u> | <u>0.01</u> | <u>1: negative regulation of programmed cell death (5:7)</u> | <u>UNC5B, SLC40A1, PDX1, ITGAV, MDM2, RRM2B, TAF9B, BAX, ITSN1</u> |
| <u>60548</u> | <u>692</u> | <u>9</u> | <u>0.03</u> | <u>1: negative regulation of cell death (4:6)</u> | <u>UNC5B, SLC40A1, PDX1, ITGAV, MDM2, RRM2B, TAF9B, BAX, ITSN1</u> |
| <u>70925</u> | <u>668</u> | <u>9</u> | <u>0.02</u> | <u>1: organelle assembly (5:5)</u> | <u>TTBK2, TMOD2, KIF3C, CENPW, SPATA6, CEP70, RAB33B, OFD1, TP53INP1</u> |
| <u>122</u> | <u>592</u> | <u>8</u> | <u>0.03</u> | <u>1: negative regulation of transcription by RNA polymerase II (7:13)</u> | <u>TSHZ1, TXNIP, TRIM29, PDX1, SMAD7, MDM2, ZFXH3, TAF9B</u> |
| <u>2001233</u> | <u>321</u> | <u>8</u> | <u>3.90E-05</u> | <u>1: regulation of apoptotic signaling pathway (5:8)</u> | <u>UNC5B, PDX1, ITGAV, MDM2, RRM2B, TAF9B, BAX, TP53INP1</u> |
| <u>44770</u> | <u>522</u> | <u>8</u> | <u>0.01</u> | <u>1: cell cycle phase transition (4:5)</u> | <u>GADD45A, MDM2, BAX, RRM2, TYMS, CEP70, OFD1, PKMYT1</u> |
| <u>44772</u> | <u>492</u> | <u>8</u> | <u>6.20E-03</u> | <u>1: mitotic cell cycle phase transition (5:6)</u> | <u>GADD45A, MDM2, BAX, RRM2, TYMS, CEP70, OFD1, PKMYT1</u> |
| <u>2001234</u> | <u>174</u> | <u>7</u> | <u>5.70E-08</u> | <u>1: negative regulation of apoptotic signaling pathway (5:9)</u> | <u>UNC5B, PDX1, ITGAV, MDM2, RRM2B, TAF9B, BAX</u> |
| <u>46982</u> | <u>353</u> | <u>6</u> | <u>0.02</u> | <u>2: protein heterodimerization activity (5:5)</u> | <u>GADD45A, GCA, GTF2A2, CENPW, TAF9B, BAX</u> |
| <u>72331</u> | <u>249</u> | <u>6</u> | <u>8.10E-04</u> | <u>1: signal transduction by p53 class mediator (5:7)</u> | <u>GADD45A, MDM2, RRM2B, TAF9B, BAX, TP53INP1</u> |
| <u>104004</u> | <u>231</u> | <u>5</u> | <u>6.90E-03</u> | <u>1: cellular response to environmental stimulus (4:4)</u> | <u>DDB2, GADD45A, MDM2, BAX, TP53INP1</u> |
| <u>2001242</u> | <u>136</u> | <u>5</u> | <u>3.50E-05</u> | <u>1: regulation of intrinsic apoptotic signaling pathway (6:9)</u> | <u>PDX1, MDM2, RRM2B, TAF9B, BAX</u> |
| <u>33218</u> | <u>209</u> | <u>5</u> | <u>3.10E-03</u> | <u>2: amide binding (3:3)</u> | <u>BACE1, SLC40A1, ITGAV, LAPTM4B, TYMS</u> |
| <u>44843</u> | <u>238</u> | <u>5</u> | <u>8.60E-03</u> | <u>1: cell cycle G1/S phase transition (5:6)</u> | <u>GADD45A, MDM2, BAX, RRM2, TYMS</u> |
| <u>71214</u> | <u>231</u> | <u>5</u> | <u>6.90E-03</u> | <u>1: cellular response to abiotic stimulus (4:5)</u> | <u>DDB2, GADD45A, MDM2, BAX, TP53INP1</u> |
| <u>71478</u> | <u>140</u> | <u>5</u> | <u>5.00E-05</u> | <u>1: cellular response to radiation (5:6)</u> | <u>DDB2, GADD45A, MDM2, BAX, TP53INP1</u> |
| <u>82</u> | <u>225</u> | <u>5</u> | <u>5.60E-03</u> | <u>1: G1/S transition of mitotic cell cycle (6:7)</u> | <u>GADD45A, MDM2, BAX, RRM2, TYMS</u> |
| <u>97193</u> | <u>252</u> | <u>5</u> | <u>0.01</u> | <u>1: intrinsic apoptotic signaling pathway (5:7)</u> | <u>PDX1, MDM2, RRM2B, TAF9B, BAX</u> |

**Supplementary Table 7.** GO term analysis Flu<Ctrl

|  |  |  |  |  |  |
| --- | --- | --- | --- | --- | --- |
| #Using 2X2 contingency table analysis (chi-square) with continuity correction |  |  |  |  |  |
| #Expected ratio: 0.00192740122068744 |  |  |  |  |  |
| **#H:goid** | **R#** | **#** | **p_value** | **Goid-Desc** | **GeneSymbols** |
| 10562 | 686 | 6 | 1.80E-04 | 1: positive regulation of phosphorus metabolic process (5:7) | CSNK2B, XRCC6, EHD4, PTPN1, RAP1A, RPS6KA5 |
| 10648 | 933 | 5 | 0.04 | 1: negative regulation of cell communication (4:6) | IGFBP6, PTPN1, RAP1A, SFRP5, NLRX1 |
| 19220 | 1192 | 6 | 0.03 | 1: regulation of phosphate metabolic process (6:7) | CSNK2B, XRCC6, EHD4, PTPN1, RAP1A, RPS6KA5 |
| 1932 | 942 | 6 | 4.40E-03 | 1: regulation of protein phosphorylation (7:9) | CSNK2B, XRCC6, EHD4, PTPN1, RAP1A, RPS6KA5 |
| 1934 | 616 | 6 | 4.80E-05 | 1: positive regulation of protein phosphorylation (7:10) | CSNK2B, XRCC6, EHD4, PTPN1, RAP1A, RPS6KA5 |
| 23057 | 936 | 5 | 0.04 | 1: negative regulation of signaling (3:5) | IGFBP6, PTPN1, RAP1A, SFRP5, NLRX1 |
| 2682 | 916 | 5 | 0.03 | 1: regulation of immune system process (3:4) | XRCC6, PTPN1, RAP1A, NLRX1, RPS6KA5 |
| 31347 | 477 | 5 | 1.40E-04 | 1: regulation of defense response (5:6) | XRCC6, PTPN1, FNDC4, NLRX1, RPS6KA5 |
| 31399 | 1263 | 6 | 0.04 | 1: regulation of protein modification process (6:8) | CSNK2B, XRCC6, EHD4, PTPN1, RAP1A, RPS6KA5 |
| 31401 | 789 | 6 | 8.50E-04 | 1: positive regulation of protein modification process (6:9) | CSNK2B, XRCC6, EHD4, PTPN1, RAP1A, RPS6KA5 |
| 32270 | 1061 | 7 | 1.10E-03 | 1: positive regulation of cellular protein metabolic process (5:8) | CSNK2B, XRCC6, EHD4, PTPN1, RAP1A, EIF3D, RPS6KA5 |
| <u>3676</u> | <u>3219</u> | <u>1</u><br><u>1</u> | <u>0.04</u> | <u>2: nucleic acid binding (4:4)</u> | <u>HNRNPA3, ZBTB12, XRCC6, PRPF31, H1F0, EHD4, CPSF3, PTPN1, RPS11, SLIRP, EIF3D</u> |
| <u>3723</u> | <u>1507</u> | <u>9</u> | <u>4.60E-04</u> | <u>2: RNA binding (5:5)</u> | <u>HNRNPA3, XRCC6, PRPF31, H1F0, CPSF3, PTPN1, RPS11, SLIRP, EIF3D</u> |
| <u>42325</u> | <u>1045</u> | <u>6</u> | <u>0.01</u> | <u>1: regulation of phosphorylation (7:8)</u> | <u>CSNK2B, XRCC6, EHD4, PTPN1, RAP1A, RPS6KA5</u> |
| <u>42327</u> | <u>649</u> | <u>6</u> | <u>9.40E-05</u> | <u>1: positive regulation of phosphorylation (7:9)</u> | <u>CSNK2B, XRCC6, EHD4, PTPN1, RAP1A, RPS6KA5</u> |
| <u>43170</u> | <u>6864</u> | <u>1</u><br><u>9</u> | <u>0.03</u> | <u>1: macromolecule metabolic process (4:4)</u> | <u>CSNK2B, ATN1, HNRNPA3, ZBTB12, XRCC6, PRPF31, H1F0, EHD4, HMMR, IGFBP6, CPSF3, GALNT10, PTPN1, RAP1A, RPS11, SFRP5, SLIRP, EIF3D, RPS6KA5</u> |
| <u>45937</u> | <u>686</u> | <u>6</u> | <u>1.80E-04</u> | <u>1: positive regulation of phosphate metabolic process (6:8)</u> | <u>CSNK2B, XRCC6, EHD4, PTPN1, RAP1A, RPS6KA5</u> |
| <u>48519</u> | <u>3625</u> | <u>1</u><br><u>2</u> | <u>0.04</u> | <u>1: negative regulation of biological process (2:4)</u> | <u>CSNK2B, ATN1, XRCC6, IGFBP6, PTPN1, RAP1A, RPS11, SFRP5, FNDC4, NLRX1, SLIRP, RPS6KA5</u> |
| <u>50776</u> | <u>573</u> | <u>5</u> | <u>9.30E-04</u> | <u>1: regulation of immune response (4:5)</u> | <u>XRCC6, PTPN1, RAP1A, NLRX1, RPS6KA5</u> |
| <u>51174</u> | <u>1194</u> | <u>6</u> | <u>0.03</u> | <u>1: regulation of phosphorus metabolic process (5:6)</u> | <u>CSNK2B, XRCC6, EHD4, PTPN1, RAP1A, RPS6KA5</u> |
| <u>51247</u> | <u>1128</u> | <u>7</u> | <u>2.10E-03</u> | <u>1: positive regulation of protein metabolic process (5:7)</u> | <u>CSNK2B, XRCC6, EHD4, PTPN1, RAP1A, EIF3D, RPS6KA5</u> |
| <u>6807</u> | <u>7468</u> | <u>2</u><br><u>0</u> | <u>0.03</u> | <u>1: nitrogen compound metabolic process (3:3)</u> | <u>CSNK2B, ATN1, HNRNPA3, ZBTB12, XRCC6, PRPF31, H1F0, EHD4, HMMR, IGFBP6, PIPOX, CPSF3, GALNT10, PTPN1, RAP1A, RPS11, SFRP5, SLIRP, EIF3D, RPS6KA5</u> |
| <u>6952</u> | <u>941</u> | <u>5</u> | <u>0.04</u> | <u>1: defense response (4:4)</u> | <u>XRCC6, PTPN1, FNDC4, NLRX1, RPS6KA5</u> |
| <u>6955</u> | <u>1229</u> | <u>6</u> | <u>0.03</u> | <u>1: immune response (3:3)</u> | <u>CSNK2B, XRCC6, PTPN1, RAP1A, NLRX1, RPS6KA5</u> |
| <u>70848</u> | <u>458</u> | <u>5</u> | <u>8.60E-05</u> | <u>1: response to growth factor (5:5)</u> | <u>EHD4, PTPN1, RAP1A, SFRP5, FNDC4</u> |
| <u>7167</u> | <u>706</u> | <u>5</u> | <u>5.50E-03</u> | <u>1: enzyme linked receptor protein signaling pathway (5:7)</u> | <u>CSNK2B, IGFBP6, PTPN1, SFRP5, RPS6KA5</u> |

**Supplementary Table 8.**
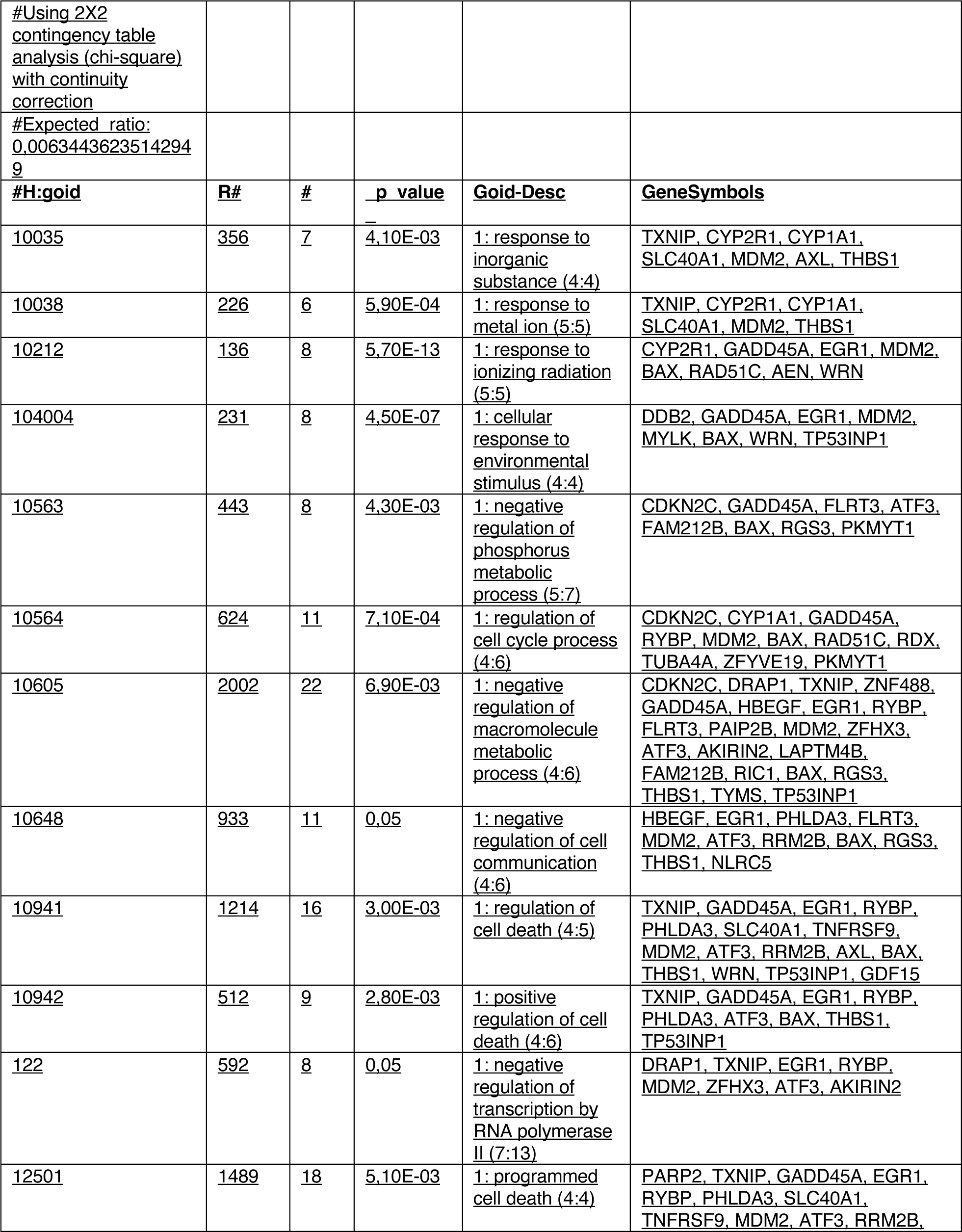
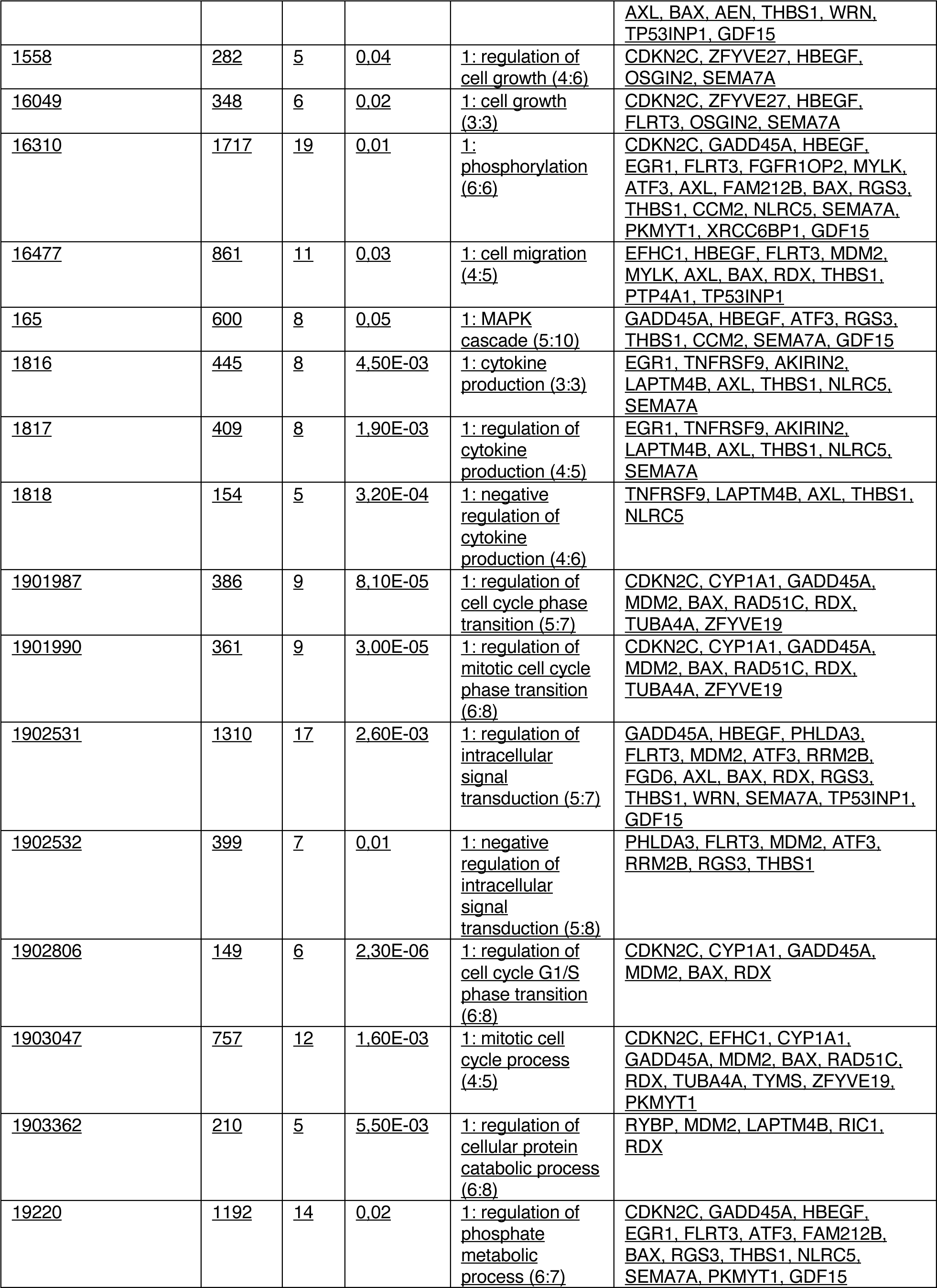
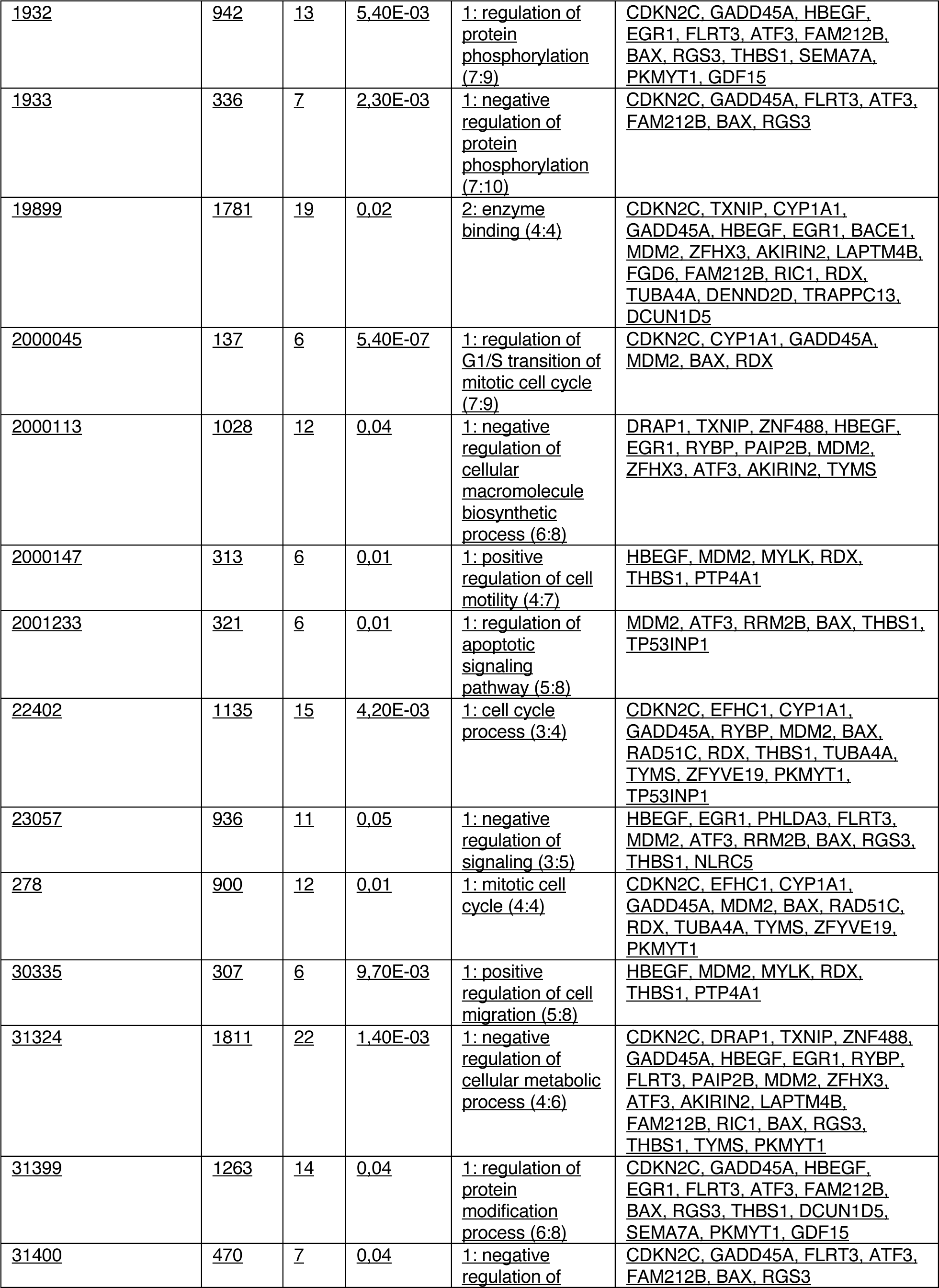
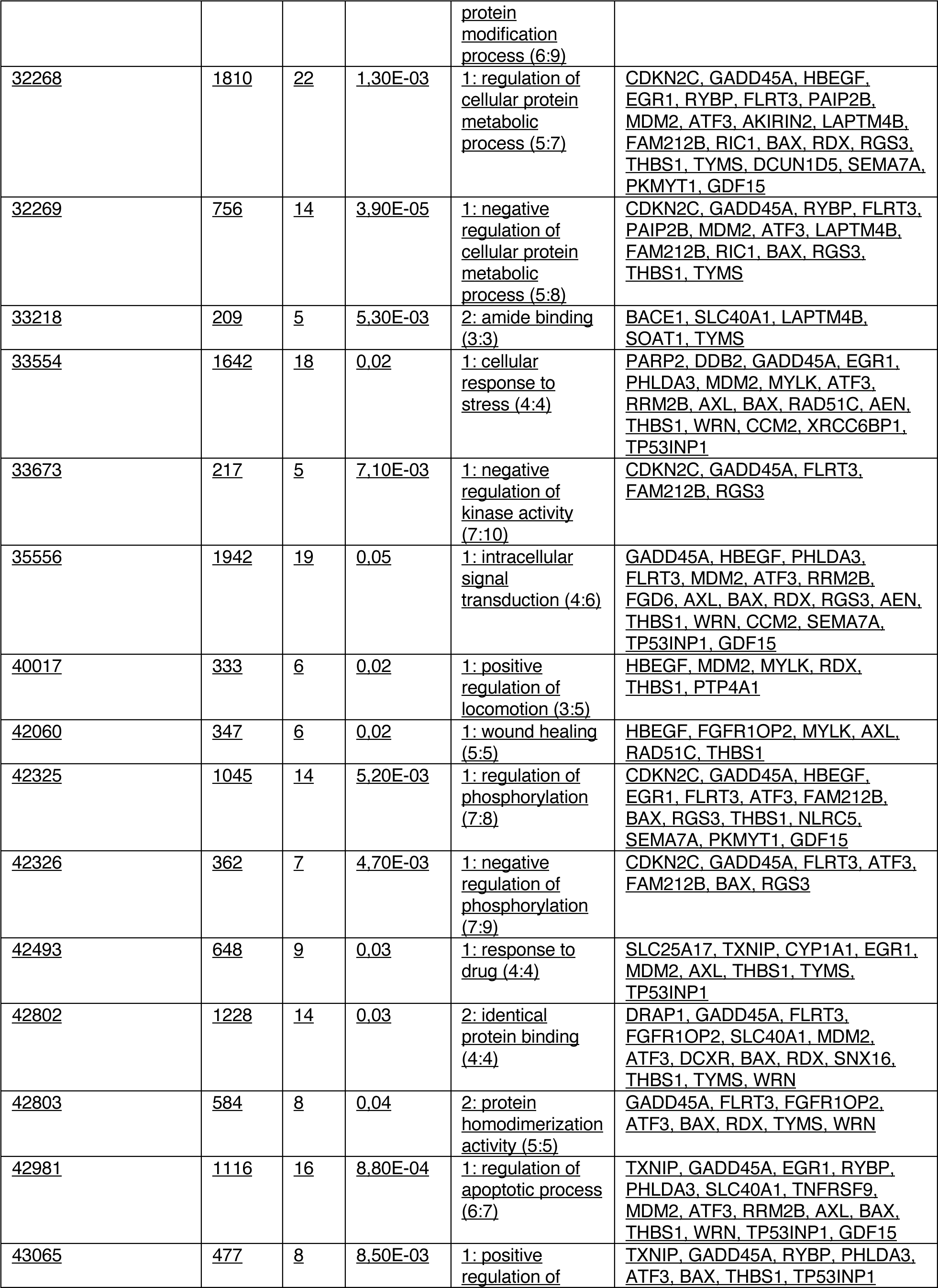
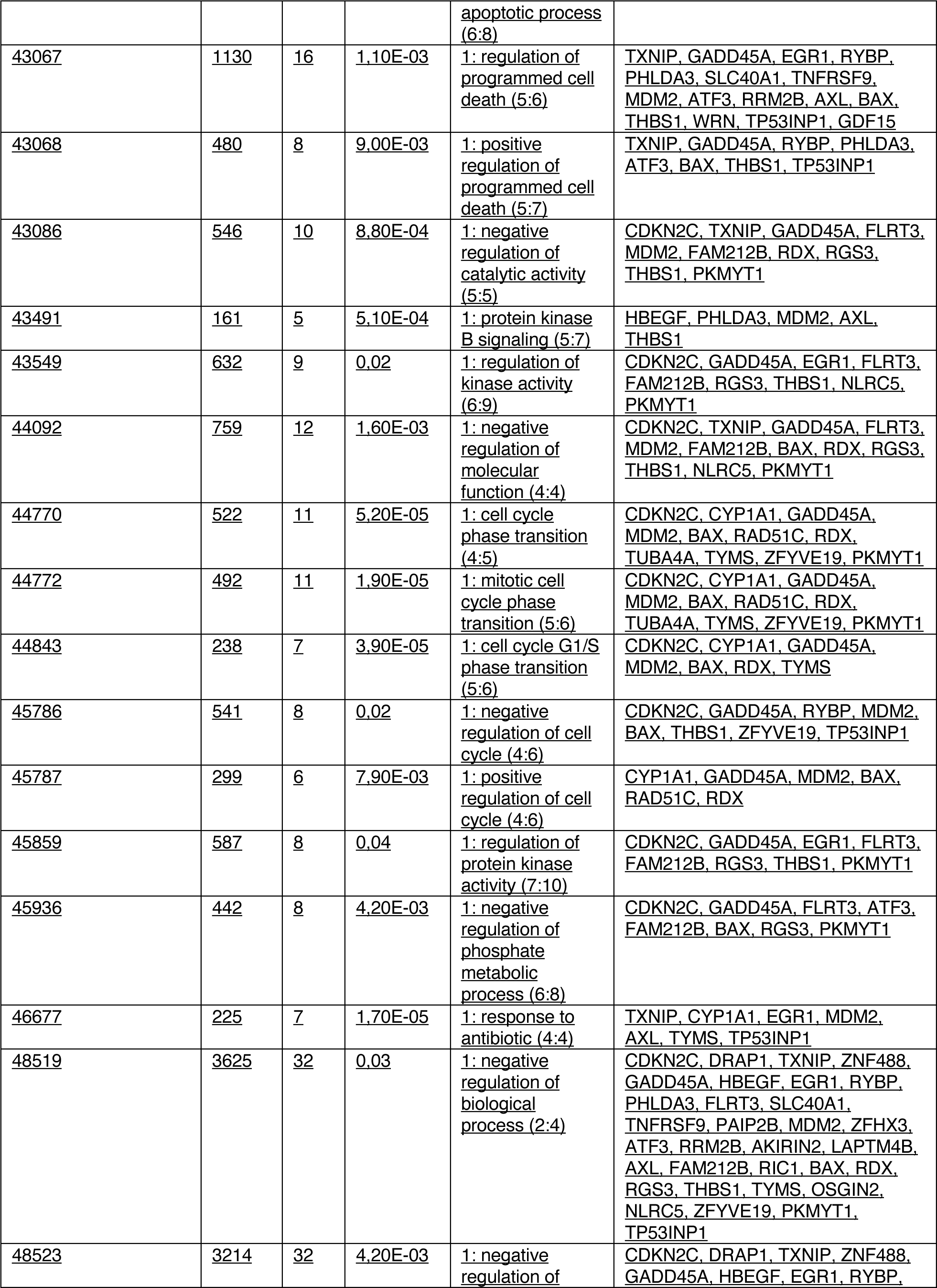
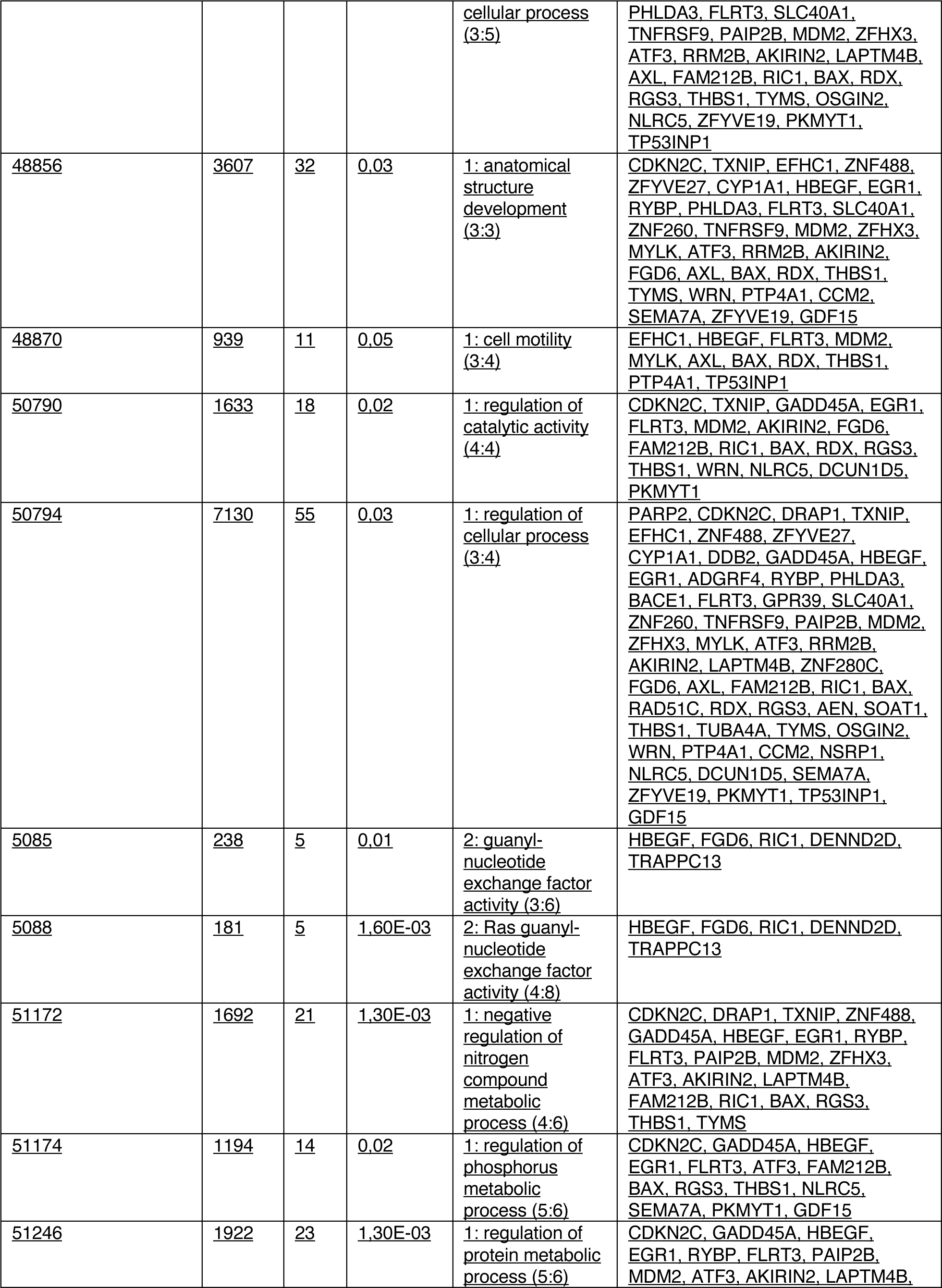
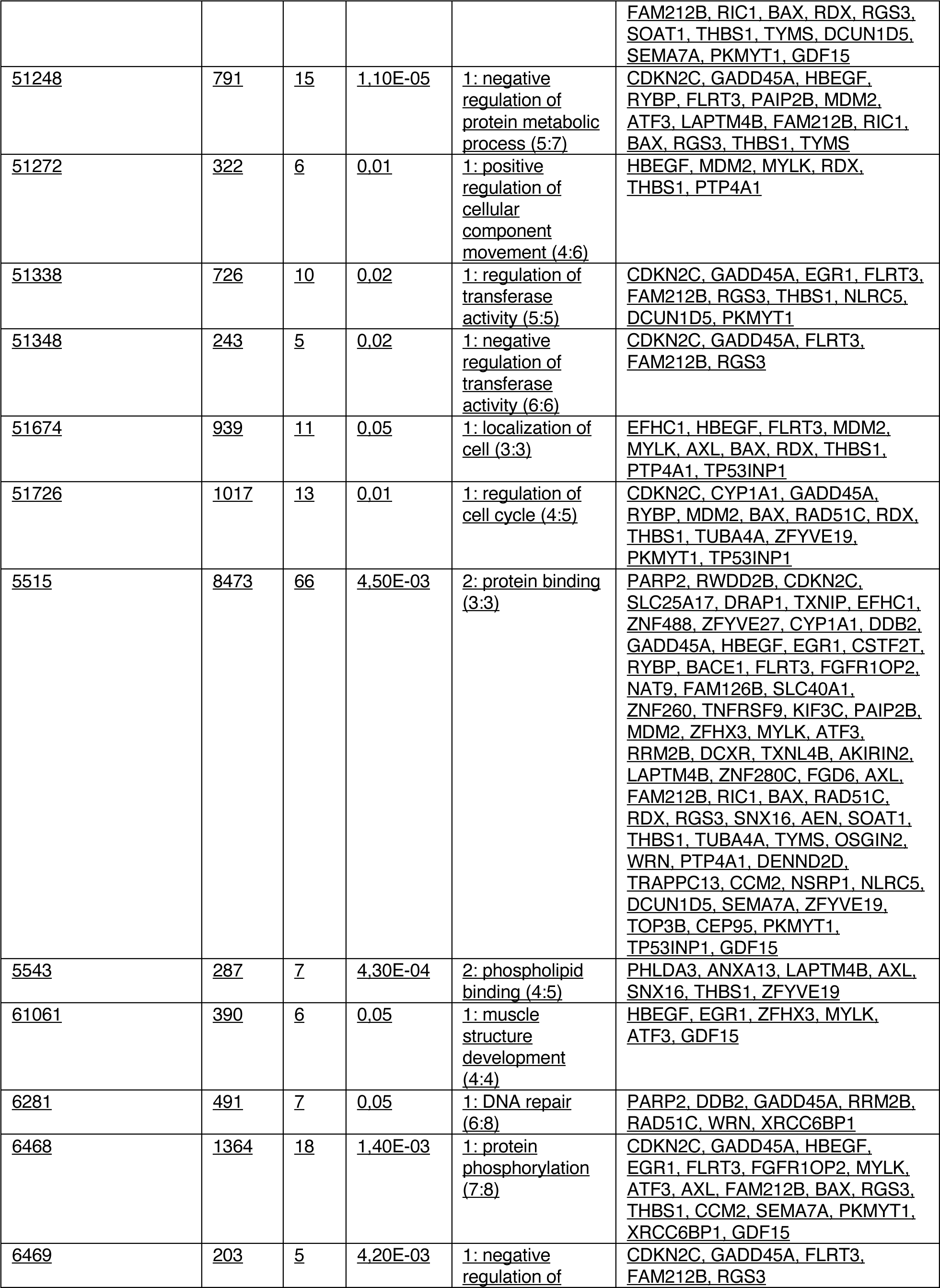
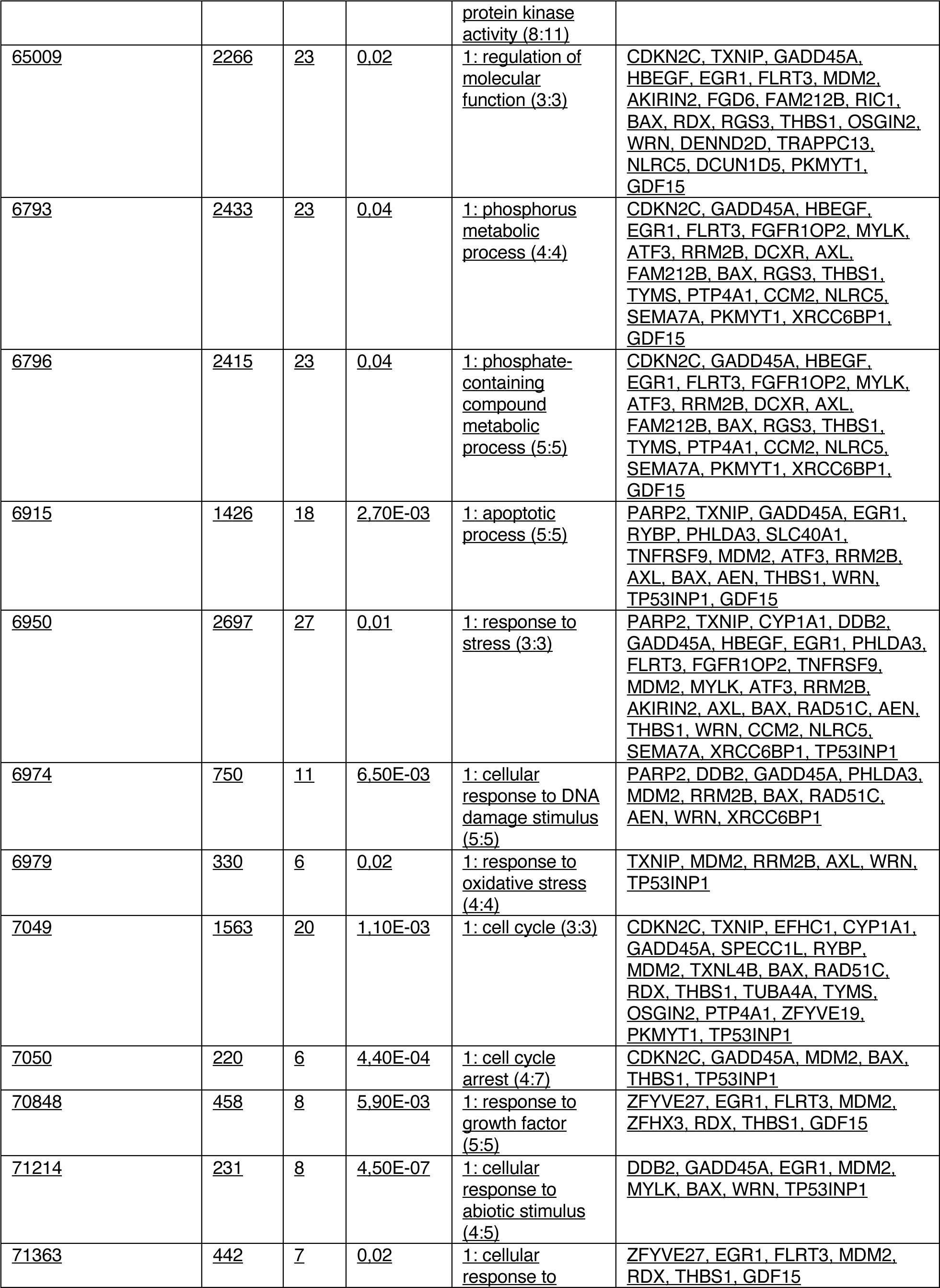
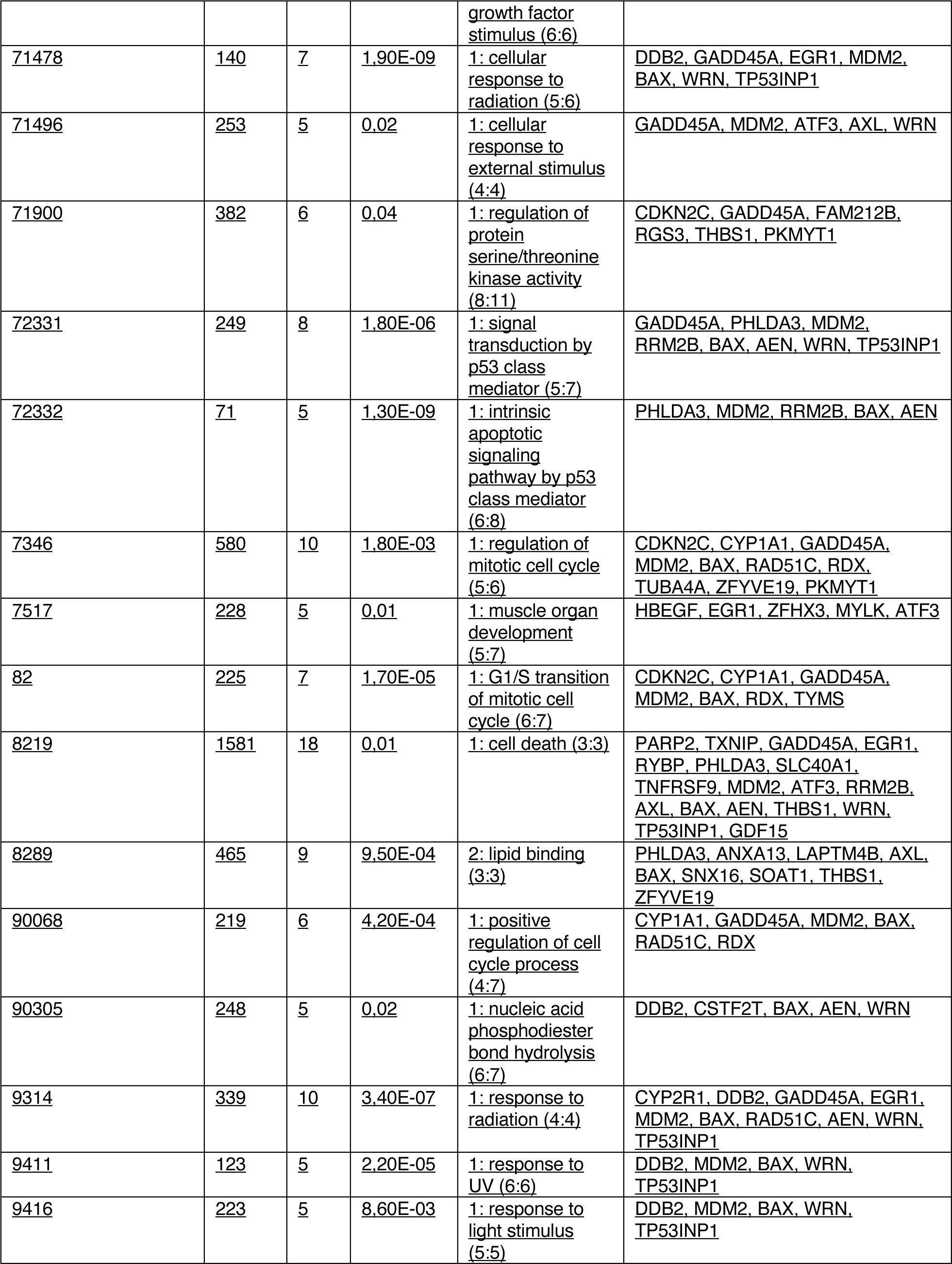
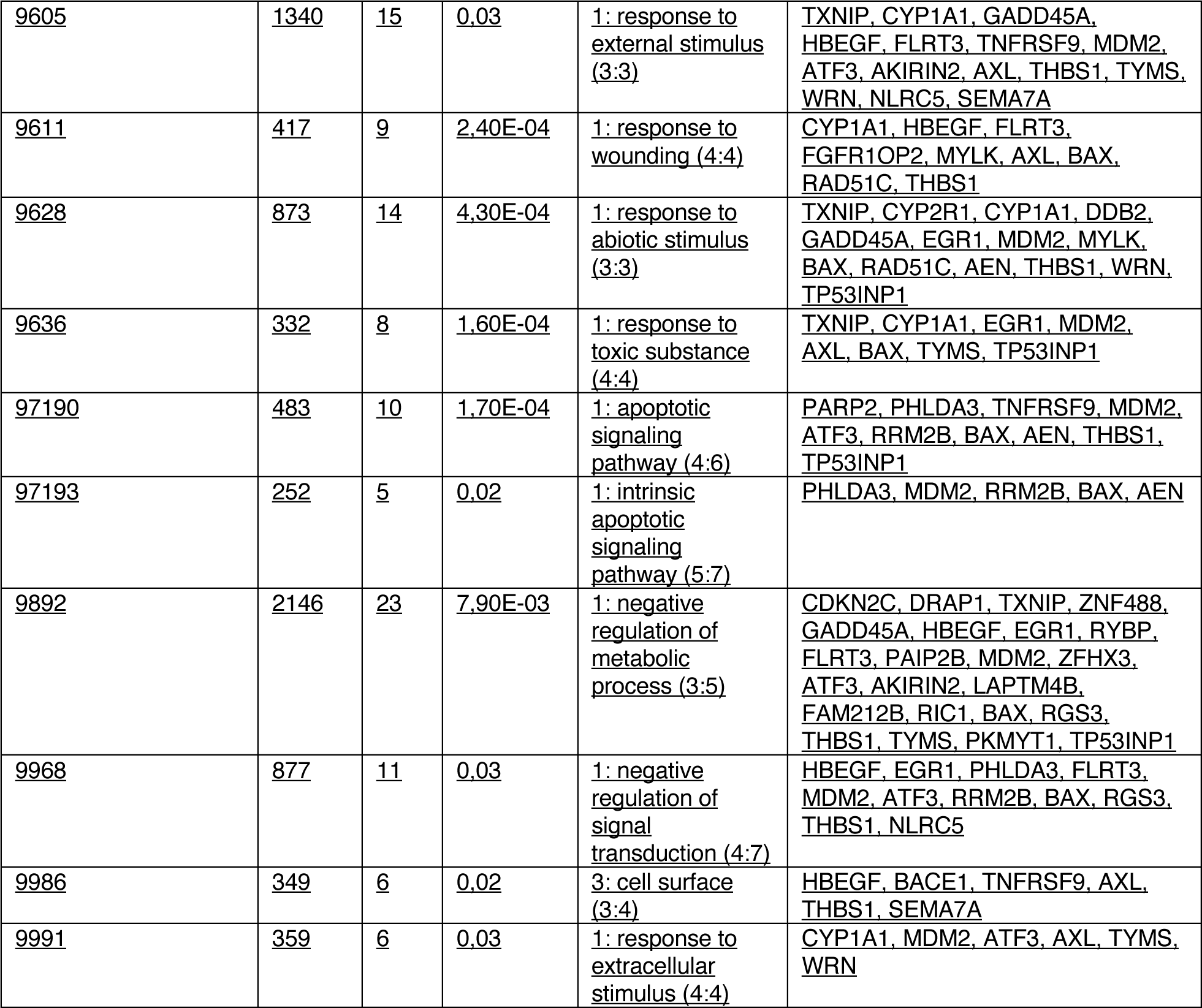
GO term analysis Clo≥Ctrl

**Supplementary Table 9.**
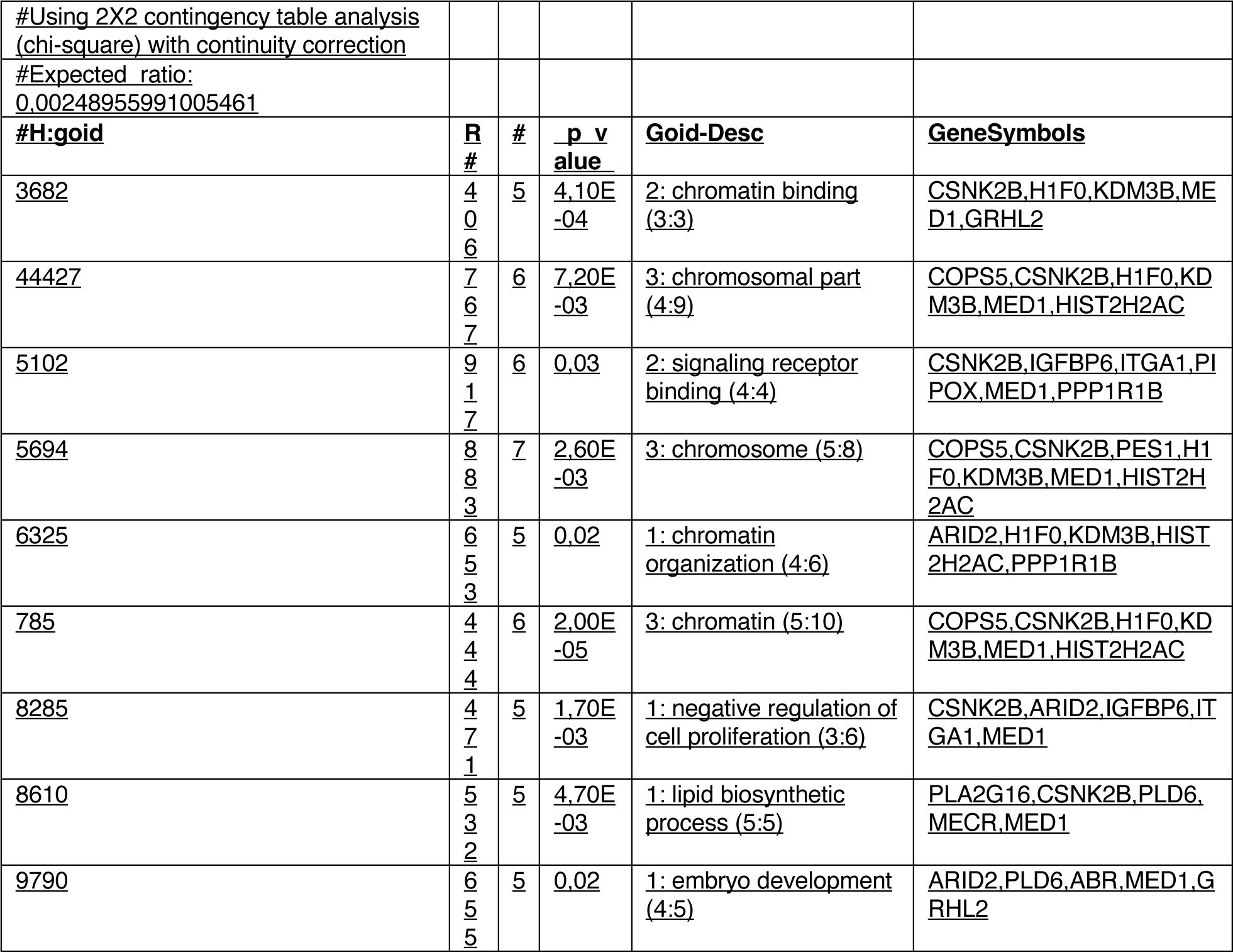
GO term analysis Clo<Ctrl

**Supplementary Table 10.** OLINK proteins immuno-oncology panel

|  |
| --- |
| <u>IL-8</u> |
| <u>TNFRSF9</u> |
| <u>TIE2</u> |
| <u>MCP-3</u> |
| <u>CD40-L</u> |
| <u>IL-1 alpha</u> |
| <u>CD244</u> |
| <u>EGF</u> |
| <u>ANGPT1</u> |
| <u>IL-7</u> |
| <u>PGF</u> |
| <u>IL-6</u> |
| <u>ADGRG1</u> |
| <u>MCP-1</u> |
| <u>CRTAM</u> |
| <u>CXCL11</u> |
| <u>MCP-4</u> |
| <u>TRAIL</u> |
| <u>FGF2</u> |
| <u>CXCL9</u> |
| <u>CD8A</u> |
| <u>CAIX</u> |
| <u>MUC-16</u> |
| <u>ADA</u> |
| <u>CD4</u> |
| <u>NOS3</u> |
| <u>IL-2</u> |
| <u>Gal-9</u> |
| <u>VEGFR-2</u> |
| <u>CD40</u> |
| <u>IL-18</u> |
| <u>GZMH</u> |
| <u>KIR3DL1</u> |
| <u>LAP TGF-beta-1</u> |
| <u>CXCL1</u> |
| <u>TNFSF14</u> |
| <u>IL-33</u> |
| <u>TWEAK</u> |
| <u>PDGF subunit B</u> |
| <u>PDCD1</u> |
| <u>FASLG</u> |
| <u>CD28</u> |
| <u>CCL19</u> |
| <u>MCP-2</u> |
| <u>CCL4</u> |
| <u>IL-15</u> |
| <u>Gal-1</u> |
| <u>PD-L1</u> |
| <u>CD27</u> |
| <u>CXCL5</u> |
| <u>IL-5</u> |
| <u>HGF</u> |
| <u>GZMA</u> |
| <u>HO-1</u> |
| <u>CXCL1</u> |
| <u>CXCL10</u> |
| <u>CD70</u> |
| <u>IL-10</u> |
| <u>TNFRSF12A</u> |
| <u>CCL23</u> |
| <u>CD5</u> |
| <u>CCL3</u> |
| <u>MMP7</u> |
| <u>ARG1</u> |
| <u>NCR1</u> |
| <u>DCN</u> |
| <u>TNFRSF21</u> |
| <u>TNFRSF4</u> |
| <u>MIC-A/B</u> |
| <u>CCL17</u> |
| <u>ANGPT2</u> |
| <u>PTN</u> |
| <u>CXCL12</u> |
| <u>IFN-gamma</u> |
| <u>LAMP3</u> |
| <u>CASP-8</u> |
| <u>ICOSLG</u> |
| <u>MMP12</u> |
| <u>CXCL13</u> |
| <u>PD-L2</u> |
| <u>VEGFA</u> |
| <u>IL-4</u> |
| <u>LAG3</u> |
| <u>IL12RB1</u> |
| <u>IL-13</u> |
| <u>CCL20</u> |
| <u>TNF</u> |
| <u>KLRD1</u> |
| <u>GZMB</u> |
| <u>CD83</u> |
| <u>IL-12</u> |
| <u>CSF-1</u> |

## References

1. Barker N, van Es JH, Kuipers J, et al. Identification of stem cells in small intestine and colon by marker gene Lgr5. Nature. 2007;449(7165):1003-1007. doi:10.1038/nature06196

2. Odenwald MA, Turner JR. The intestinal epithelial barrier: a therapeutic target? Nat Rev Gastroenterol Hepatol. 2017;14(1):9–21. doi:10.1038/nrgastro.2016.169

3. Odenwald MA, Turner JR. Intestinal permeability defects: is it time to treat? Clin Gastroenterol Hepatol. 2013;11(9):1075–1083. doi:10.1016/j.cgh.2013.07.001

4. Turpin W, Lee SH, Raygoza Garay JA, et al. Increased Intestinal Permeability Is Associated With Later Development of Crohn’s Disease. Gastroenterology. 2020;159(6):2092–2100.e5. doi:10.1053/j.gastro.2020.08.005

5. Torres J, Petralia F, Sato T, et al. Serum Biomarkers Identify Patients Who Will Develop Inflammatory Bowel Diseases Up to 5 Years Before Diagnosis. Gastroenterology. 2020;159(1):96–104. doi:10.1053/j.gastro.2020.03.007

6. Abadie V, Kim SM, Lejeune T, et al. IL-15, gluten and HLA-DQ8 drive tissue destruction in coeliac disease. Nature. 2020;578(7796):600-604. doi:10.1038/s41586-020-2003-8

7. Dotsenko V, Oittinen M, Taavela J, et al. Genome-Wide Transcriptomic Analysis of Intestinal Mucosa in Celiac Disease Patients on a Gluten-Free Diet and Postgluten Challenge. Cell Mol Gastroenterol Hepatol. 2021;11(1):13–32. doi:10.1016/j.jcmgh.2020.07.010

8. Nalle SC, Zuo L, Ong MLDM, et al. Graft-versus-host disease propagation depends on increased intestinal epithelial tight junction permeability. J Clin Invest. 2019;129(2):902–914. doi:10.1172/JCI98554

9. Jansen SA, Nieuwenhuis EES, Hanash AM, Lindemans CA. Challenges and opportunities targeting mechanisms of epithelial injury and recovery in acute intestinal graft-versus-host disease. Mucosal Immunol. 2022;15(4):605–619. doi:10.1038/s41385-022-00527-6

10. van Wijk F, Cheroutre H. Mucosal T cells in gut homeostasis and inflammation. Expert Rev Clin Immunol. 2010;6(4):559–566. doi:10.1586/eci.10.34

11. Boussios S, Pentheroudakis G, Katsanos K, Pavlidis N. Systemic treatment-induced gastrointestinal toxicity: incidence, clinical presentation and management. Ann Gastroenterol. 2012;25(2):106–118.

12. Elting LS, Cooksley C, Chambers M, Cantor SB, Manzullo E, Rubenstein EB. The burdens of cancer therapy. Clinical and economic outcomes of chemotherapy-induced mucositis. Cancer. 2003;98(7):1531–1539. doi:10.1002/cncr.11671

13. Fischer JC, Wintges A, Haas T, Poeck H. Assessment of mucosal integrity by quantifying neutrophil granulocyte influx in murine models of acute intestinal injury. Cell Immunol. 2017;316:70–76. doi:10.1016/j.cellimm.2017.04.003

14. Johansson JE, Brune M, Ekman T. The gut mucosa barrier is preserved during allogeneic, haemopoietic stem cell transplantation with reduced intensity conditioning. Bone Marrow Transplant. 2001;28(8):737–742. doi:10.1038/sj.bmt.1703230

15. Johansson JE, Ekman T. Gut toxicity during hemopoietic stem cell transplantation may predict acute graft-versus-host disease severity in patients. Dig Dis Sci. 2007;52(9):2340–2345. doi:10.1007/s10620-006-9404-x

16. Ferrara JLM, Levine JE, Reddy P, Holler E. Graft-versus-host disease. Lancet. 2009;373(9674):1550-1561. doi:10.1016/S0140-6736(09)60237-3

17. Zeiser R, Blazar BR. Acute Graft-versus-Host Disease. N Engl J Med. 2018;378(6):586. doi:10.1056/NEJMc1716969

18. Jagasia M, Arora M, Flowers MED, et al. Risk factors for acute GVHD and survival after hematopoietic cell transplantation. Blood. 2012;119(1):296–307. doi:10.1182/blood-2011-06-364265

19. Nakasone H, Fukuda T, Kanda J, et al. Impact of conditioning intensity and TBI on acute GVHD after hematopoietic cell transplantation. Bone Marrow Transplant. 2015;50(4):559-565. doi:10.1038/bmt.2014.293

20. Flowers MED, Inamoto Y, Carpenter PA, et al. Comparative analysis of risk factors for acute graft-versus-host disease and for chronic graft-versus-host disease according to National Institutes of Health consensus criteria. Blood. 2011;117(11):3214-3219. doi:10.1182/blood-2010-08-302109

21. Liu D, Yan C, Xu L, et al. Diarrhea during the conditioning regimen is correlated with the occurrence of severe acute graft-versus-host disease through systemic release of inflammatory cytokines. Biol Blood Marrow Transplant. 2010;16(11):1567-1575. doi:10.1016/j.bbmt.2010.05.001

22. Koyama M, Kuns RD, Olver SD, et al. Recipient nonhematopoietic antigen-presenting cells are sufficient to induce lethal acute graft-versus-host disease. Nat Med. 2011;18(1):135–142. doi:10.1038/nm.2597

23. Koyama M, Hill GR. The primacy of gastrointestinal tract antigen-presenting cells in lethal graft-versus-host disease. Blood. 2019;134(24):2139–2148. doi:10.1182/blood.2019000823

24. Koyama M, Mukhopadhyay P, Schuster IS, et al. MHC Class II Antigen Presentation by the Intestinal Epithelium Initiates Graft-versus-Host Disease and Is Influenced by the Microbiota. Immunity. 2019;51(5):885–898.e7. doi:10.1016/j.immuni.2019.08.011

25. Yu Y, Jin QR, Mi Y, et al. Intestinal Epithelial Cell-Derived CD83 Contributes to Regulatory T-Cell Generation and Inhibition of Food Allergy. J Innate Immun. 2021;13(5):295–305. doi:10.1159/000515332

26. Biton M, Haber AL, Rogel N, et al. T Helper Cell Cytokines Modulate Intestinal Stem Cell Renewal and Differentiation. Cell. 2018;175(5):1307–1320.e22. doi:10.1016/j.cell.2018.10.008

27. Harrison OJ, Srinivasan N, Pott J, et al. Epithelial-derived IL-18 regulates Th17 cell differentiation and Foxp3^+^ Treg cell function in the intestine. Mucosal Immunol. 2015;8(6):1226–1236. doi:10.1038/mi.2015.13

28. Schiering C, Krausgruber T, Chomka A, et al. The alarmin IL-33 promotes regulatory T- cell function in the intestine. Nature. 2014;513(7519):564-568. doi:10.1038/nature13577

29. Watanabe M, Ueno Y, Yajima T, et al. Interleukin 7 is produced by human intestinal epithelial cells and regulates the proliferation of intestinal mucosal lymphocytes. J Clin Invest. 1995;95(6):2945–2953. doi:10.1172/JCI118002

30. Fu YY, Egorova A, Sobieski C, et al. T Cell Recruitment to the Intestinal Stem Cell Compartment Drives Immune-Mediated Intestinal Damage after Allogeneic Transplantation. Immunity. 2019;51(1):90–103.e3. doi:10.1016/j.immuni.2019.06.003

31. Sujino T, London M, Hoytema van Konijnenburg DP, et al. Tissue adaptation of regulatory and intraepithelial CD4^+^ T cells controls gut inflammation. Science. 2016352(6293):1581-1586. doi:10.1126/science.aaf3892

32. Hoytema van Konijnenburg DP, Reis BS, Pedicord VA, Farache J, Victora GD, Mucida D. Intestinal Epithelial and Intraepithelial T Cell Crosstalk Mediates a Dynamic Response to Infection. Cell. 2017;171(4):783–794.e13. doi:10.1016/j.cell.2017.08.046

33. Takashima S, Martin ML, Jansen SA, et al. T cell-derived interferon-gamma programs stem cell death in immune-mediated intestinal damage. Sci Immunol. 2019;4(42). doi:10.1126/sciimmunol.aay8556

34. Sato T, Vries RG, Snippert HJ, et al. Single Lgr5 stem cells build crypt-villus structures in vitro without a mesenchymal niche. Nature. 2009459(7244):262-265. doi:10.1038/nature07935

35. Sato T, Stange DE, Ferrante M, et al. Long-term expansion of epithelial organoids from human colon, adenoma, adenocarcinoma, and Barrett’s epithelium. Gastroenterology. 2011;141(5):1762–1772. doi:10.1053/j.gastro.2011.07.050

36. Lindemans CA, Calafiore M, Mertelsmann AM, et al. Interleukin-22 promotes intestinal- stem-cell-mediated epithelial regeneration. Nature. 2015528(7583):560-564. doi:10.1038/nature16460

37. Bar-Ephraim YE, Kretzschmar K, Clevers H. Organoids in immunological research. Nat Rev Immunol. 2020;20(5):279–293. doi:10.1038/s41577-019-0248-y

38. Kelsen JR, Dawany N, Conrad MA, et al. Colonoids From Patients With Pediatric Inflammatory Bowel Disease Exhibit Decreased Growth Associated With Inflammation Severity and Durable Upregulation of Antigen Presentation Genes. Inflamm Bowel Dis. 2021;27(2):256–267. doi:10.1093/ibd/izaa145

39. Rana N, Privitera G, Kondolf HC, et al. GSDMB is increased in IBD and regulates epithelial restitution/repair independent of pyroptosis. Cell. 2022;185(2):283–298.e17. doi:10.1016/j.cell.2021.12.024

40. Li H, Durbin R. Fast and accurate short read alignment with Burrows-Wheeler transform. Bioinformatics. 2009;25(14):1754–1760. doi:10.1093/bioinformatics/btp324

41. Love MI, Huber W, Anders S. Moderated estimation of fold change and dispersion for RNA-seq data with DESeq2. Genome Biol. 2014;15(12):550. doi:10.1186/s13059-014-0550-8

42. Subramanian A, Tamayo P, Mootha VK, et al. Gene set enrichment analysis: a knowledge-based approach for interpreting genome-wide expression profiles. Proc Natl Acad Sci U S A. 2005;102(43):15545–15550. doi:10.1073/pnas.0506580102

43. Assarsson E, Lundberg M, Holmquist G, et al. Homogenous 96-plex PEA immunoassay exhibiting high sensitivity, specificity, and excellent scalability. PLoS One. 2014;9(4):e95192. doi:10.1371/journal.pone.0095192

44. Kebriaei P, Bassett R, Lyons G, et al. Clofarabine Plus Busulfan is an Effective Conditioning Regimen for Allogeneic Hematopoietic Stem Cell Transplantation in Patients with Acute Lymphoblastic Leukemia: Long-Term Study Results. Biol Blood Marrow Transplant. 2017;23(2):285–292. doi:10.1016/j.bbmt.2016.11.001

45. Alatrash G, Thall PF, Valdez BC, et al. Long-Term Outcomes after Treatment with Clofarabine ± Fludarabine with Once-Daily Intravenous Busulfan as Pretransplant Conditioning Therapy for Advanced Myeloid Leukemia and Myelodysplastic Syndrome. Biol Blood Marrow Transplant. 2016;22(10):1792–1800. doi:10.1016/j.bbmt.2016.06.023

46. Kebriaei P, Basset R, Ledesma C, et al. Clofarabine combined with busulfan provides excellent disease control in adult patients with acute lymphoblastic leukemia undergoing allogeneic hematopoietic stem cell transplantation. Biol Blood Marrow Transplant. 2012;18(12):1819–1826. doi:10.1016/j.bbmt.2012.06.010

47. Valdez BC, Li Y, Murray D, Champlin RE, Andersson BS. The synergistic cytotoxicity of clofarabine, fludarabine and busulfan in AML cells involves ATM pathway activation and chromatin remodeling. Biochem Pharmacol. 2011;81(2):222–232. doi:10.1016/j.bcp.2010.09.027

48. Ben Hassine K, Powys M, Svec P, et al. Total Body Irradiation Forever? Optimising Chemotherapeutic Options for Irradiation-Free Conditioning for Paediatric Acute Lymphoblastic Leukaemia. Front Pediatr. 2021;9:775485. doi:10.3389/fped.2021.775485

49. Bartelink IH, van Reij EML, Gerhardt CE, et al. Fludarabine and exposure-targeted busulfan compares favorably with busulfan/cyclophosphamide-based regimens in pediatric hematopoietic cell transplantation: maintaining efficacy with less toxicity. Biol Blood Marrow Transplant. 2014;20(3):345–353. doi:10.1016/j.bbmt.2013.11.027

50. Versluys AB, Boelens JJ, Pronk C, et al. Hematopoietic cell transplant in pediatric acute myeloid leukemia after similar upfront therapy; a comparison of conditioning regimens. Bone Marrow Transplant. 2021;56(6):1426–1432. doi:10.1038/s41409-020-01201-w

51. Versluijs AB, de Koning CCH, Lankester AC, et al. Clofarabine-fludarabine-busulfan in HCT for pediatric leukemia: an effective, low toxicity, TBI-free conditioning regimen. Blood Adv. 2022;6(6):1719–1730. doi:10.1182/bloodadvances.2021005224

52. Iwamoto T, Hiraku Y, Oikawa S, Mizutani H, Kojima M, Kawanishi S. DNA intrastrand cross-link at the 5’-GA-3’ sequence formed by busulfan and its role in the cytotoxic effect. Cancer Sci. 2004;95(5):454–458. doi:10.1111/j.1349-7006.2004.tb03231.x

53. Ewald B, Sampath D, Plunkett W. Nucleoside analogs: molecular mechanisms signaling cell death. Oncogene. 2008;27(50):6522–6537. doi:10.1038/onc.2008.316

54. Andersson BS, Valdez BC, de Lima M, et al. Clofarabine ± fludarabine with once daily i.v. busulfan as pretransplant conditioning therapy for advanced myeloid leukemia and MDS. Biol Blood Marrow Transplant. 2011;17(6):893–900. doi:10.1016/j.bbmt.2010.09.022

55. Mah LJ, El-Osta A, Karagiannis TC. gammaH2AX: a sensitive molecular marker of DNA damage and repair. Leukemia. 2010;24(4):679–686. doi:10.1038/leu.2010.6

56. Schnalzger TE, de Groot MH, Zhang C, et al. 3D model for CAR-mediated cytotoxicity using patient-derived colorectal cancer organoids. EMBO J. 2019;38(12). doi:10.15252/embj.2018100928

57. Jacob F, Ming GL, Song H. Generation and biobanking of patient-derived glioblastoma organoids and their application in CAR T cell testing. Nat Protoc. 2020;15(12):4000–4033. doi:10.1038/s41596-020-0402-9

58. Larson RC, Kann MC, Bailey SR, et al. CAR T cell killing requires the IFNγR pathway in solid but not liquid tumours. Nature. Published online April 2022. doi:10.1038/s41586-022-04585-5

59. Dijkstra KK, Cattaneo CM, Weeber F, et al. Generation of Tumor-Reactive T Cells by Co- culture of Peripheral Blood Lymphocytes and Tumor Organoids. Cell. 2018;174(6):1586–1598.e12. doi:10.1016/j.cell.2018.07.009

60. Griffin JM, Healy FM, Dahal LN, Floisand Y, Woolley JF. Worked to the bone: antibody- based conditioning as the future of transplant biology. J Hematol Oncol. 2022;15(1):65. doi:10.1186/s13045-022-01284-6

61. Wada J, Kanwar YS. Identification and characterization of galectin-9, a novel beta- galactoside-binding mammalian lectin. J Biol Chem. 1997;272(9):6078–6086. doi:10.1074/jbc.272.9.6078

62. Türeci O, Schmitt H, Fadle N, Pfreundschuh M, Sahin U. Molecular definition of a novel human galectin which is immunogenic in patients with Hodgkin’s disease. J Biol Chem. 1997;272(10):6416–6422. doi:10.1074/jbc.272.10.6416

63. Spitzenberger F, Graessler J, Schroeder HE. Molecular and functional characterization of galectin 9 mRNA isoforms in porcine and human cells and tissues. Biochimie. 2001;83(9):851–862. doi:10.1016/s0300-9084(01)01335-9

64. Brewer CF, Miceli MC, Baum LG. Clusters, bundles, arrays and lattices: novel mechanisms for lectin–saccharide-mediated cellular interactions. Curr Opin Struct Biol. 2002;12(5):616–623. doi:10.1016/S0959-440X(02)00364-0

65. Zhu C, Anderson AC, Schubart A, et al. The Tim-3 ligand galectin-9 negatively regulates T helper type 1 immunity. Nat Immunol. 2005;6(12):1245–1252. doi:10.1038/ni1271

66. Madireddi S, Eun SY, Mehta AK, et al. Regulatory T Cell-Mediated Suppression of Inflammation Induced by DR3 Signaling Is Dependent on Galectin-9. J Immunol. 2017;199(8):2721–2728. doi:10.4049/jimmunol.1700575

67. Madireddi S, Eun SY, Lee SW, et al. Galectin-9 controls the therapeutic activity of 4-1BB- targeting antibodies. J Exp Med. 2014;211(7):1433–1448. doi:10.1084/jem.20132687

68. Wu C, Thalhamer T, Franca RF, et al. Galectin-9-CD44 interaction enhances stability and function of adaptive regulatory T cells. Immunity. 2014;41(2):270–282. doi:10.1016/j.immuni.2014.06.011

69. Bi S, Hong PW, Lee B, Baum LG. Galectin-9 binding to cell surface protein disulfide isomerase regulates the redox environment to enhance T-cell migration and HIV entry. Proc Natl Acad Sci U S A. 2011;108(26):10650–10655. doi:10.1073/pnas.1017954108

70. Yang R, Sun L, Li CF, et al. Galectin-9 interacts with PD-1 and TIM-3 to regulate T cell death and is a target for cancer immunotherapy. Nat Commun. 2021;12(1):832. doi:10.1038/s41467-021-21099-2

71. Gooden MJM, Wiersma VR, Samplonius DF, et al. Galectin-9 activates and expands human T-helper 1 cells. PLoS One. 2013;8(5):e65616. doi:10.1371/journal.pone.0065616

72. Kashio Y, Nakamura K, Abedin MJ, et al. Galectin-9 induces apoptosis through the calcium-calpain-caspase-1 pathway. J Immunol. 2003;170(7):3631–3636. doi:10.4049/jimmunol.170.7.3631

73. Lu LH, Nakagawa R, Kashio Y, et al. Characterization of galectin-9-induced death of Jurkat T cells. J Biochem. 2007;141(2):157–172. doi:10.1093/jb/mvm019

74. Lhuillier C, Barjon C, Niki T, et al. Impact of Exogenous Galectin-9 on Human T Cells: CONTRIBUTION OF THE T CELL RECEPTOR COMPLEX TO ANTIGEN- INDEPENDENT ACTIVATION BUT NOT TO APOPTOSIS INDUCTION. J Biol Chem. 2015;290(27):16797–16811. doi:10.1074/jbc.M115.661272

75. Lee J, Park EJ, Noh JW, et al. Underexpression of TIM-3 and blunted galectin-9-induced apoptosis of CD4+ T cells in rheumatoid arthritis. Inflammation. 2012;35(2):633–637. doi:10.1007/s10753-011-9355-z

76. Chen HY, Wu YF, Chou FC, et al. Intracellular Galectin-9 Enhances Proximal TCR Signaling and Potentiates Autoimmune Diseases. J Immunol. 2020;204(5):1158–1172. doi:10.4049/jimmunol.1901114

77. Su W, Zhang J, Yang S, et al. Galectin-9 contributes to the pathogenesis of atopic dermatitis via T cell immunoglobulin mucin-3. Front Immunol. 2022;13:952338. doi:10.3389/fimmu.2022.952338

78. Sakai K, Kawata E, Ashihara E, et al. Galectin-9 ameliorates acute GVH disease through the induction of T-cell apoptosis. Eur J Immunol. 2011;41(1):67–75. doi:10.1002/eji.200939931

79. Veenstra RG, Taylor PA, Zhou Q, et al. Contrasting acute graft-versus-host disease effects of Tim-3/galectin-9 pathway blockade dependent upon the presence of donor regulatory T cells. Blood. 2012;120(3):682–690. doi:10.1182/blood-2011-10-387977

80. Kandel S, Adhikary P, Li G, Cheng K. The TIM3/Gal9 signaling pathway: An emerging target for cancer immunotherapy. Cancer Lett. 2021;510:67–78. doi:10.1016/j.canlet.2021.04.011

81. Yang R, Sun L, Li CF, et al. Galectin-9 interacts with PD-1 and TIM-3 to regulate T cell death and is a target for cancer immunotherapy. Nat Commun. 2021;12(1):832. doi:10.1038/s41467-021-21099-2

82. Yang R, Sun L, Li CF, et al. Development and characterization of anti-galectin-9 antibodies that protect T cells from galectin-9-induced cell death. J Biol Chem. 2022;298(4):101821. doi:10.1016/j.jbc.2022.101821

## References Supplemental Methods

1. Segeren HA, van Liere EA, Riemers FM, de Bruin A, Westendorp B. Oncogenic RAS sensitizes cells to drug-induced replication stress via transcriptional silencing of P53. Oncogene. 2022;41(19):2719–2733. doi:10.1038/s41388-022-02291-0

2. de Jager W, Prakken BJ, Bijlsma JWJ, Kuis W, Rijkers GT. Improved multiplex immunoassay performance in human plasma and synovial fluid following removal of interfering heterophilic antibodies. J Immunol Methods. 2005;300(1- 2):124–135. doi:10.1016/j.jim.2005.03.009

